# Multinuclear non-heme iron dependent oxidative enzymes: Landscape of their substrates, partner proteins and biosynthetic gene clusters

**DOI:** 10.1101/2025.01.10.632315

**Authors:** R. Antoine, L. Leprevost, S. Jünger, S. Zirah, G. Lippens, Y. Li, S. Dubiley, F. Jacob-Dubuisson

## Abstract

Proteins of the multinuclear non-heme iron-dependent oxidative (MNIO) enzyme superfamily catalyze various modification reactions on the precursors of ribosomally synthesized, post-translationally modified peptides (RiPPs). We recently identified two large families of MNIO-modified RiPPs called bufferins, which enhance bacterial growth under copper stress by chelating the excess metal ions. Here, we explored the diversity of potential MNIO substrates by performing extensive *in silico* studies. Analyses of MNIO-coding biosynthetic gene clusters (BGCs) identified various groups of putative precursors most of which are characterized by specific Cys-containing motifs, throughout the eubacterial phylogenetic tree. The precursors of most MNIO-modified RiPPs harbor N-terminal Sec-dependent signal peptides, a rare feature among bacterial RiPPs. Some precursors are very long relative to those of typical RiPPs, indicating that MNIO enzymes could modify both peptide and protein substrates. We also identified a distinct family of integral membrane proteins with large predicted extra-cytoplasmic domains mostly found in Actinomycetota, frequently but not systematically associated with MNIOs. Most MNIO BGCs harbor genes coding for DUF2063 domain-containing proteins or structurally related proteins, serving as partners of the enzymes for precursor modification. We uncovered a correlation between the presence or the absence of Sec signal peptides in the precursors and the types of partner proteins of the MNIO enzymes. This study depicts the global landscape of potential MNIO-dependent natural products by unveiling groups of peptides and proteins genetically associated with MNIOs. It reveals a treasure trove of potential new RiPP precursors which likely represent a widespread bacterial strategy to deal with copper stress, and most likely other stresses, in natural environments.

**Impact statement:** The multinuclear non-heme iron-dependent oxidative (MNIO) enzymes belong to an emerging superfamily of modification enzymes that catalyze various oxidation reactions on ribosomally synthesized post-translationally modified peptides (RiPP) precursors.

The largest families of MNIO-modified RiPPs, called bufferins, are involved in copper homeostasis. In this work we performed extensive *in silico* analyses to explore the diversity of RiPP precursors genetically associated with MNIO enzymes and identified major families. Some precursors are much larger than typical RiPP precursors, indicating that MNIO enzymes may also modify proteins. We defined subtypes of MNIO enzymes’ partner proteins dedicated to specific families of precursors. Our analyses of the biosynthetic gene clusters unveiled functions beyond copper homeostasis, likely for the response to other metal stresses. The global landscape of MNIO-modified RiPP precursors will be a basis for investigations into new RiPP families and will undoubtedly lead to the discovery of new modifications and new functions.

## Introduction

Peptide-based natural products can notably be synthesized by ribosomes and then enzymatically modified like ribosomally synthesized post-translationally modified peptides (RiPPs) or fully synthesized by enzymatic pathways (1, 2). RiPPs are highly prevalent natural products in bacteria, with several dozen families described over the last decades (2–5). Some eukaryotic organisms, particularly plants, cone snails and fungi, also produce RiPPs (6).

Bacterial RiPPs are encoded in biosynthetic gene clusters (BGCs) that comprise at least one precursor gene and one or more gene(s) coding for modification enzymes. The RiPP precursors harbor a core region that undergoes enzymatic installation of post-translational modifications (PTMs), generally flanked at its N-terminus by a leader peptide (7), or more rarely at its C terminus by a follower peptide (8). In many instances the leader peptide is recognized by a system-specific helix-winged-helix motif-containing ‘RiPP recognition element’ (RRE) that can be a discrete domain or the C-or N-terminal domain of a larger protein (9–11) and that presents the substrate to the modification enzyme(s) (9, 12). Several modes of interaction between the RiPP precursor and the RRE domain as well as RRE domain-independent modes of precursor recognition have been reported (9, 11, 13–17). RiPP BGCs also frequently comprise genes coding for maturation proteases and export systems (18, 19), as well as for cognate immunity proteins in the case of RiPP bacteriocins (20).

RiPPs modified by members of the multinuclear non-heme iron oxidative (MNIO) enzymes (Pfam short name MbnB_TglH_ChrH; Pfam entry PF05114) superfamily represent a vast, emerging RiPP class, as described in a recent review (21) (Table 1). MNIO enzymes catalyze various 2-or 4-electron oxidation reactions targeting mostly Cys residues and frequently work with partner proteins, named DUF2063 (Domain of Unknown Function) (Pfam entry PF09836), Anthrone_oxy (Pfam entry PF08592) or methbact_MbnC (NCBIFAM entry TIGR04160) (22–24). The first MNIO-modified RiPPs identified, methanobactins, are chalkophores that scavenge copper in the milieu for the assembly of cuproenzymes called methane monooxygenases in Methanobacteria (24–26). Methanobactins were identified in non-methanogenic bacteria as well (27). In complex with its partner protein MbnC, the MNIO enzyme MbnB oxidizes specific internal Cys residues of the MbnA methanobactin precursor to form copper-chelating oxazolone thioamide groups (26). Other MNIO enzymes modify C-terminal Cys residues first added to precursor scaffold peptides, thus generating thia-amino acid derivatives called ‘pearlins’ (28, 29). Yet another type of MNIO-modified RiPP, chryseobasin harbors an imidazolidinedione heterocycle, a macrocycle, thioaminals and a thiomethyl group (23). Non-Cys modifying MNIO enzymes were described to exert their catalytic activity on C-terminal Asp or Asn residues, generating C-terminally amidated peptides or aminopyruvic acid, respectively (27, 30).

**Table 1.**
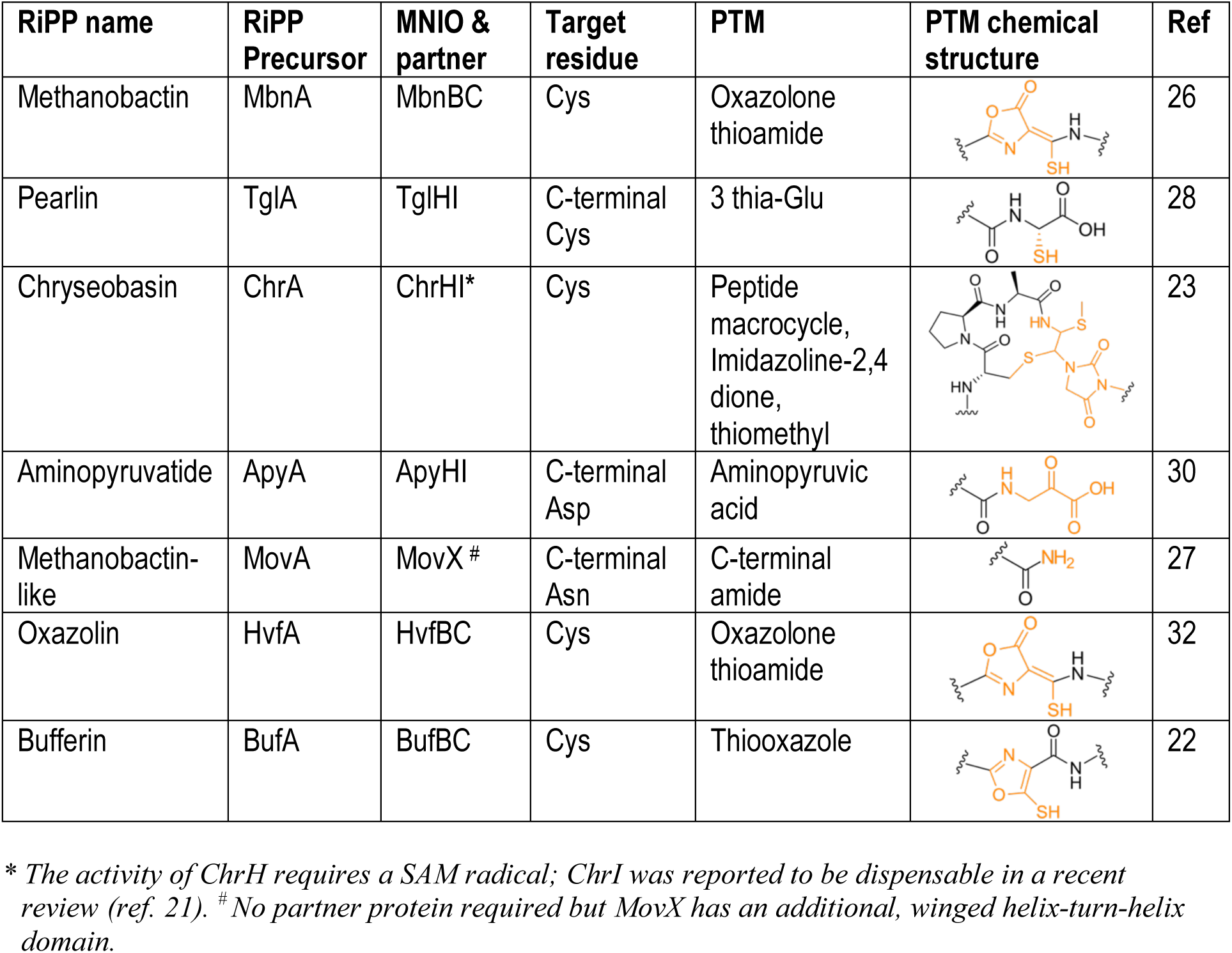
Known types of MNIO-modified RiPPs.

We have described two large families of RiPPs modified by MNIO enzymes involved in copper homeostasis and called them bufferins (22). Bufferins belong to the Buf1 (named DUF2282; Pfam entry PF10048) or Buf2 (which we previously called Buf_6/12Cys) families. Copper is a necessary transition metal that is toxic in excess (31), and bufferins use Cys-derived thiooxazole heterocycles to sequester this metal in the periplasm (22). The recently reported oxazolin represents another family of MNIO-modified RiPPs that bind copper, but it appears to do so using oxazolone thioamide groups, similar to methanobactins (32).

Unlike the precursors of methanobactins, pearlins and aminopyruvatives, those of bufferins and oxazolins feature Sec-dependent N-terminal signal peptides rather than typical leader peptides, which is highly unusual for bacterial RiPP precursors (22). Although the presence of Sec signal peptides was suggested in a few other RiPP precursors, the use of the general Sec export machinery in the natural host has only been shown experimentally for bufferins (22, 33, 34). Thus, this appears to be an original feature among RiPP precursors. The partners of the bufferin-modifying MNIO enzymes are proteins composed of a DUF2063 domain fused to an RRE-type domain, another specific feature of this RiPP class (22).

Thousands of MNIOs have been identified *in silico* (22, 23, 27, 35, 36). In this work we explored the global diversity of the MNIO BGCs by genome mining approaches and identified new potential MNIO substrates including *bona fide* proteins, and new partner proteins.

## Results

### Search for MNIO-associated RiPP precursors

In order to identify the various types of putative MNIO substrates, we collected the members of the MNIO family (approx. 14,000 proteins at < 90% sequence identity) using the Pfam MbnB_TglH_ChrH signature. These enzymes are distributed across a number of bacterial taxonomic groups (Fig. 1A). Sequence similarity network (SSN) analyses showed that Buf1- and Buf2-associated MNIO enzymes represent a sizeable fraction of the largest sequence cluster of the representative node network (red and yellow dots in Fig. 1B). MNIOs genetically associated with oxazolins (hereafter called Buf_EGKCG/oxazolins, see below) are also part of this large cluster (pale blue dots in Fig. 1B). Some of the small MNIO sequence clusters contain enzymes associated with methanobactins, pearlins, methanobactin-like peptides from *Vibrio*, chryseobasins and aminopyruvatides (23, 26–28, 30). As evidenced from this analysis, many other types of potential MNIO substrates remain to be described. It is likely that the small groups of MNIO enzymes mediate the installation of a diversity of post-translational modifications.

**Figure 1.**
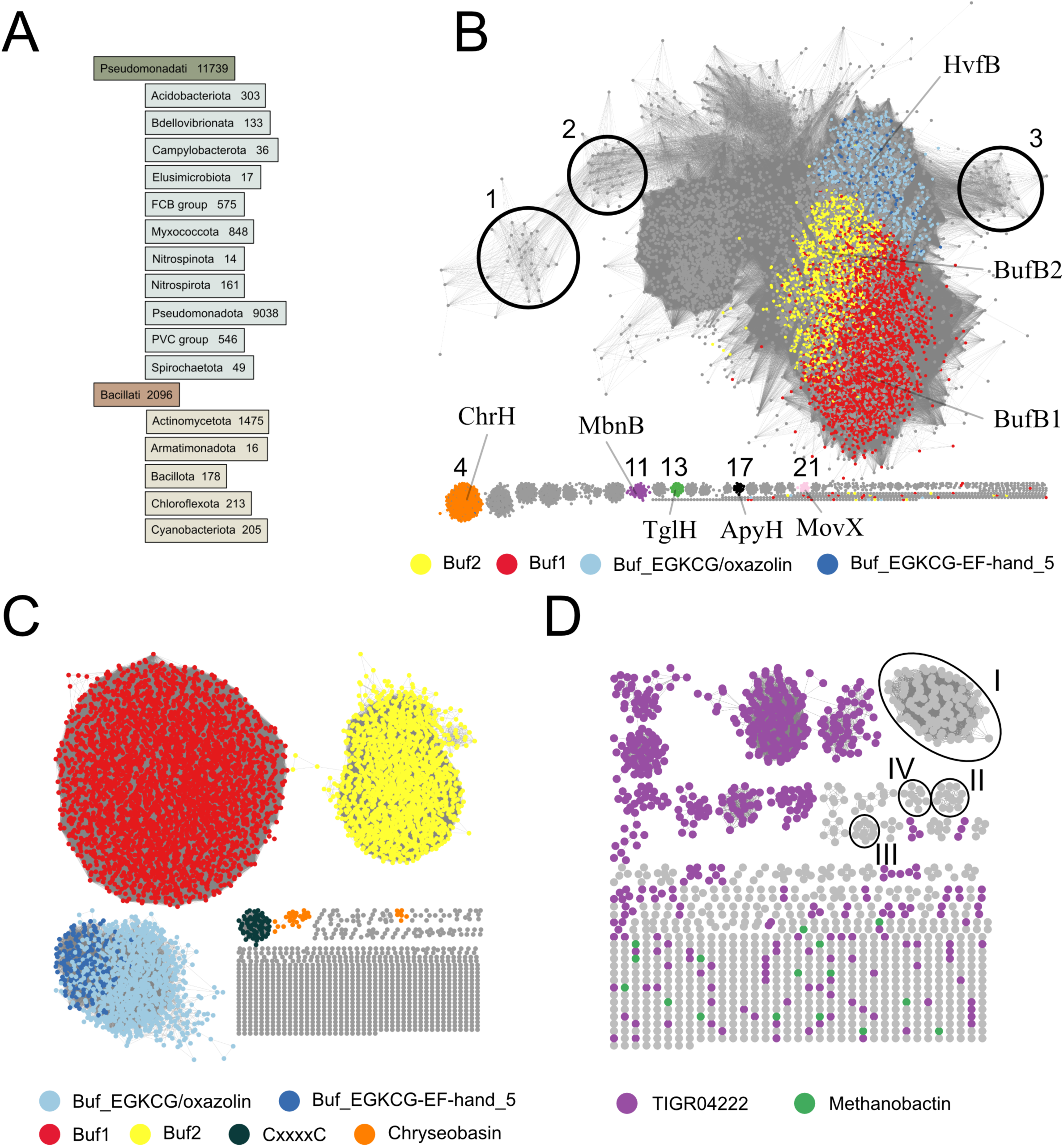
The MNIO superfamily and associated families of precursors. **A.** Taxonomic distribution of the MNIO enzymes. The absolute numbers of proteins that belong to each taxonomic group are provided. For the sake of clarity, only the groups that contain more than ten members are shown. **B,C,D.** SSN analyses. **B.** Representative node network of MNIO enzymes (alignment score threshold (AST) of 50). MNIO enzymes genetically associated with Buf1 (DUF2282), Buf2 (BUF_6/12Cys), Buf_EGKCG/oxazolins (see below) and Buf_EGKC|EF-hand_5 (see below) peptides are colored red, yellow, light blue and dark blue, respectively. The small clusters containing MNIO enzymes that modify the precursors of chryseobasins (#4), methanobactins (#11), pearlins (#13), aminopyruvatides (#17) and *Vibrio* methanobactin-like RiPPs (#21) are indicated. The clusters numbered 1 to 21 are further described later in the text and in Fig. 5. **C**. Representative node network of putative precursors genetically associated with MNIO enzymes and harboring predicted signal-peptides (AST=10). The largest clusters by decreasing sizes correspond to precursors of the Buf1 (DUF2282), Buf2 (Buf_6/12_Cys), Buf_EGKCG/oxazolin, CxxxxC (see below) and chryseobasin families. **D**. Representative node network of putative precursors genetically associated with MNIO enzymes and lacking predicted signal-peptides (AST=10). TIGR04222 proteins (purple nodes) and methanobactins (green nodes) were notably found in this search. Methanobactins do not form clusters in the node network, most likely because of their small size and low sequence identity. The largest four sequence clusters of unknown proteins (labeled I to IV) are further described in the text and in Table 2.

**Table 2.**
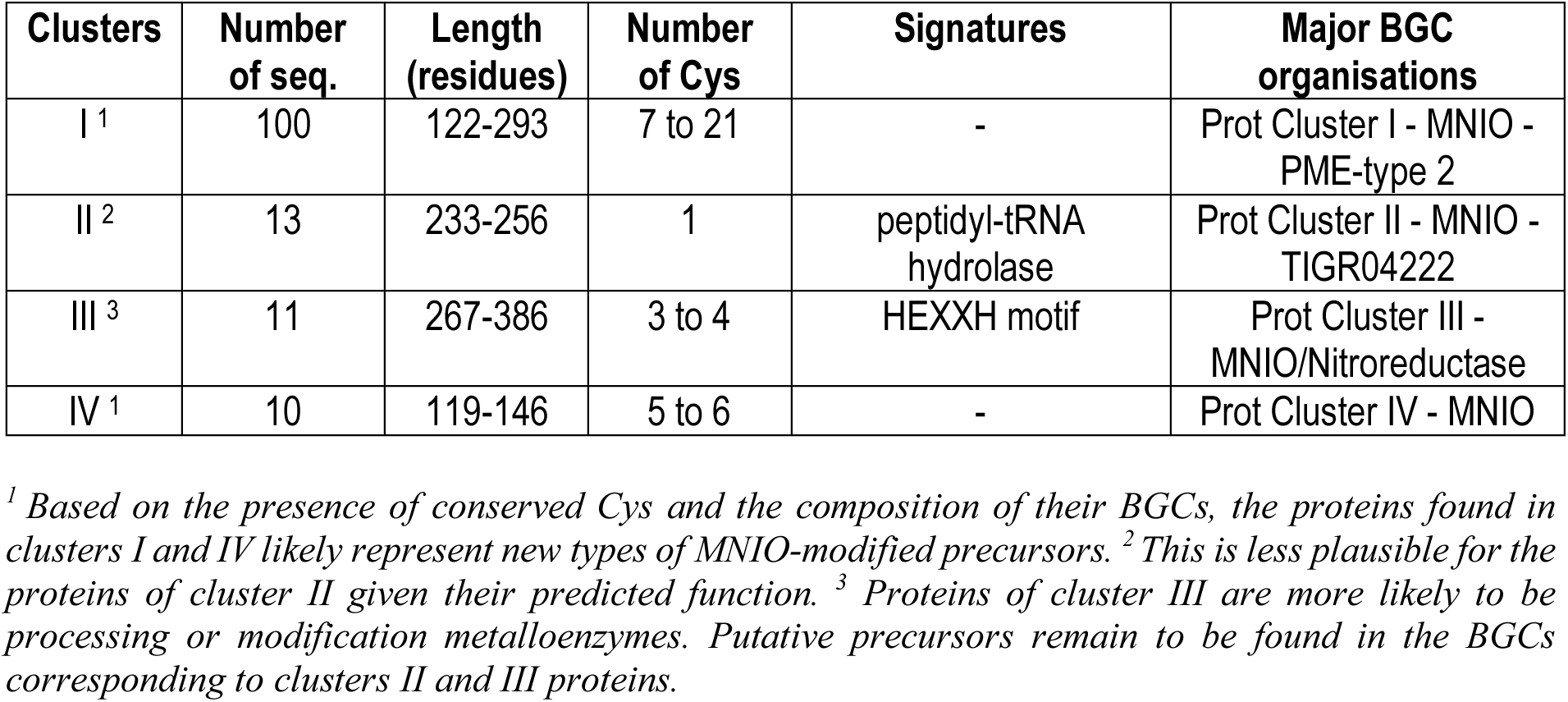
Main features of the proteins found in clusters I to IV of Fig. 1D.

We utilized a home-made RODEO-type approach (37, 38) to collect the BGCs of all these MNIO enzymes, and we analyzed their compositions (Table S1). Known domains frequently encoded in the vicinity of MNIO-coding genes notably include DUF2282, DUF2063, DoxX (Pfam entry PF07681), Sigma-70 region 4 (Pfam entry PF08281), Sigma-70 region 2 (Pfam entry PF04542), and NrsF (Pfam entry PF06532), all found in bufferin BGCs (22). Because no signatures were available yet for Buf2 or for Buf_EGKCG/oxazolins, we generated new hmm profiles for these two groups (Suppl. Files S1 and S2).

Next, we performed two-step analyses to characterize the various types of putative RiPP precursors (Fig. 2A). We first identified potential precursors in all BGCs collected, as described below, sorted them through SSN analyses and where possible we built hmm profiles for new protein families. Then, we used the available or the newly generated profiles to collect all members of each family from the NCBI nr Protein database, irrespective of their genetic environment, retrieved the corresponding BGCs and analyzed their composition.

**Figure 2.**
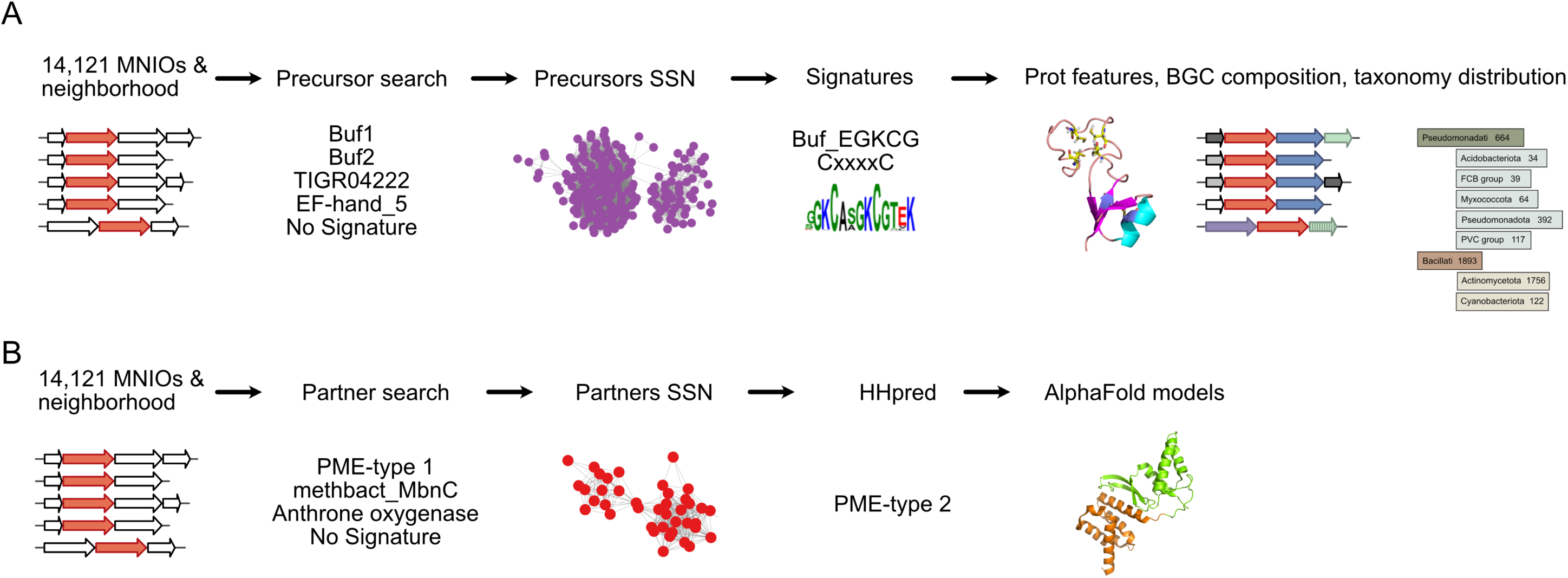
Flow chart for the identification of putative MNIO substrates and partners. **A**. Search for MNIO substrates (putative RiPP precursors). After collecting all MNIO ORFs, the ORFs that precede them were collected and analyzed for domain signatures. SSN analyses revealed new putative families, for which hmm signatures were generated. Available and new signatures were used to collect all family members in the NCBI nr protein database, irrespective of their genetic environments. These proteins and their BCGs were analyzed. **B**. Search for partner proteins of MNIO enzymes. The ORFs that follow MNIO ORFs were analyzed for domain signatures, and SSN analyses were performed with the unknown proteins. HHpred analyses were carried out for all proteins from the various sequence clusters of the SSN network, and AlphaFold2 models were generated for a few of them. Only proteins less than 90% identical in sequence were used in all analyses.

From our initial set of BGCs, we first retrieved putative precursors shorter than 200 residues, harboring Cys residues and a predicted Sec signal-peptide and coded by genes that precede the MNIO genes. These criteria were motivated by the fact that the overwhelming majority of MNIO substrates appear to modify Cys residues (21) (Fig. 1B), and by preliminary analyses showing the strict conservation of the gene order *bufABC* (encoding the bufferin precursor, the MNIO enzyme and the partner protein, respectively) in *bufferin* BGCs (22). Classification of the >11,000 identified proteins by SSN analyses revealed large sequence clusters of Buf1, Buf2 and Buf_EGKCG/oxazolin homologs (Fig. 1C). In the latter cluster a sizeable proportion of proteins correspond to the EF-hand_5 name (Pfam entry PF13202) (see below). Most other proteins, which belong to small sequence clusters, lacked known domain signatures.

To enlarge our range of putative MNIO substrates, we also retrieved from our initial BGC collection all proteins with Cys residues but devoid of predicted Sec signal-peptides and without size restriction, whose genes precede MNIO genes. This yielded >1000 proteins, 40% of which correspond to the NCBIfam entry TIGR04222. An SSN analysis with all these proteins revealed a very scattered node network, with many sequence clusters harboring TIGR04222 proteins, indicating the considerable sequence diversity of the latter (purple dots in Fig. 1D).

### General features of bufferins and their BGCs

Next, we characterized the various groups of proteins in more detail, starting with *bona fide* bufferins. We collected thousands Buf1 and Buf2 bufferins (at < 90% redundancy) from the NCBI nr Protein database, in various taxonomic groups (Fig. 3A,B). Of note, Buf2 BGCs were reported in uncultured Antarctic soil bacteria (39), suggesting their large environmental distribution.

**Figure 3.**
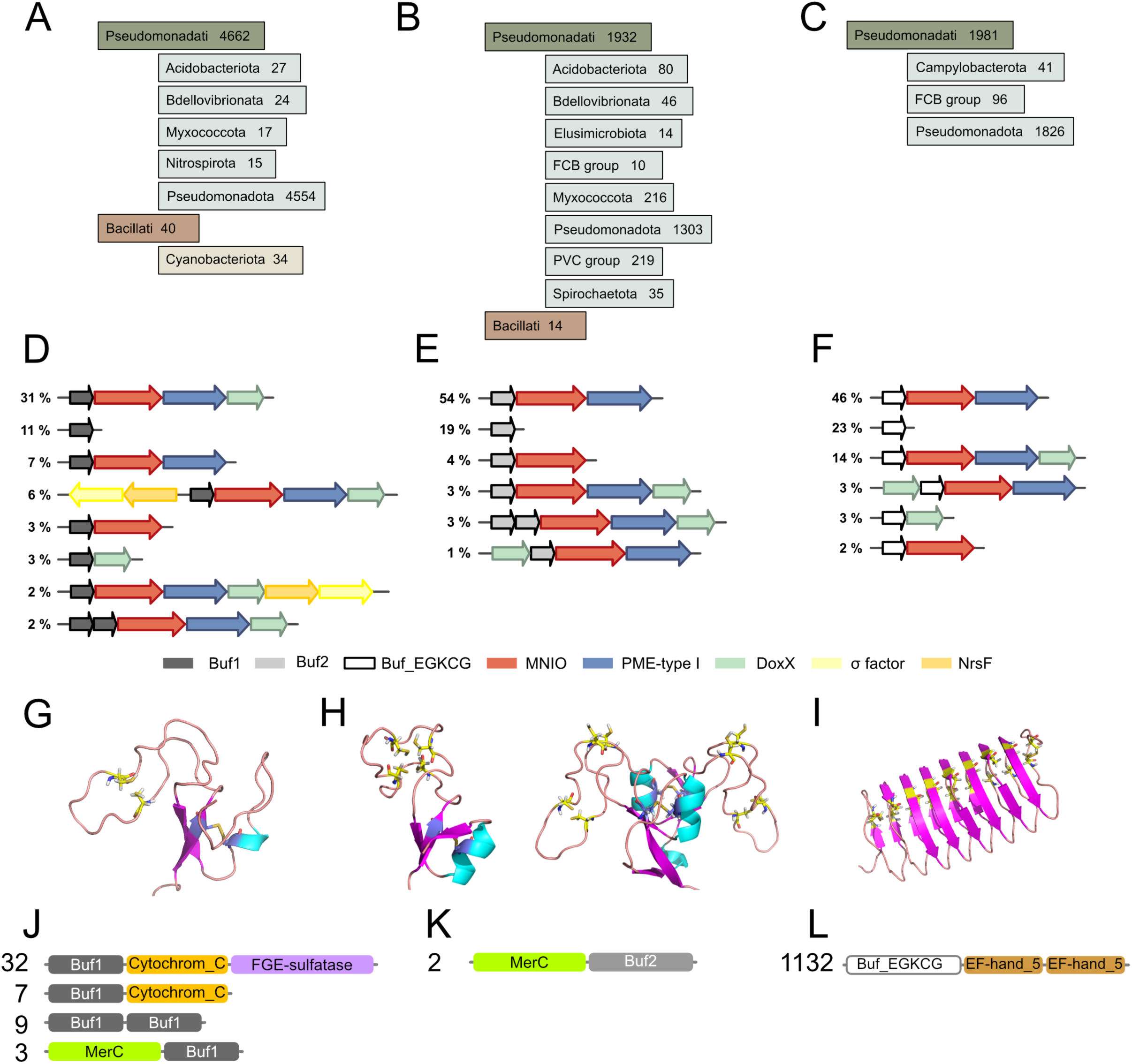
Features of precursors with signal peptides. **A-C**. Taxonomic distributions of the Buf1 (DUF2282) bufferins (A), Buf2 (Buf_6/12Cys) bufferins (B), and Buf-EGKCG/oxazolins (C). **D-F.** Major types of genetic organizations with their relative frequencies in the NCBI nr protein database. Some of the stand-alone precursor genes genuinely form single-gene transcriptional units, whereas others are flanked by genes that are not conserved to a significant extent and are thus not shown. **G-I.** Structural models of a Buf1 bufferin (WP_011519119 of *Cupriavidus metallidurans*; G), two Buf2 bufferins (WP_010947964 of *Legionella pneumophila* subsp. *fraseri* and WP_077285100 of *Cognaticolwellia aestuarii*; H) and a mid-size Buf_EGKCG protein (WP_070991527 of *Pseudoalteromonas byunsanensis*; I). The Cys residues are shown in stick representation. In Buf1, the central Cys residues are the targets of PTMs, whereas the first and last ones form a disulfide bond. In the Buf_EGKCG/oxazolin protein, the Cys side chains that might be the targets of PTMs are predicted to point toward the inside of the helix. The Buf_EGKC proteins that form the N-terminal domains of the Buf_EGKC|EF-hand_5 chimeras are predicted to be unstructured and are thus not represented. **J-L**. Numbers of occurrences and schematic representation of the fusion proteins harboring Buf1 (J), Buf2 domains (K) and Buf_EGKCG domains (L). Examples of these proteins include WP_160059411.1 (Buf1/Cytochrom_C/FGE-sulfatase); WP_017136223.1 (Buf1/Cytochrom_C); OAD22792.1 (Buf1/Buf1); WP_200808486.1 (MerC/Buf1); OOO01496.1 (MerC/Buf2) and WP_187711723.1 (Buf_EGKCG/Ef-hand_5/ Ef-hand_5). The MerC fusions are predicted to harbour five transmembrane segments, with the C-terminal bufferin domains of the chimeras on the periplasmic side of the membrane.

Genes for sigma factors and anti-sigma NsrF proteins were found in Buf1 but not in Buf2 BGCs (Fig. 3D,E). In both subfamilies, sizeable proportions of precursors are coded by genes not genetically associated with MNIO genes. We also identified BGCs with tandem precursor genes, suggesting that some MNIO enzymes might have more than one substrate. Additional conserved genes were found in extremely low proportions, confirming that the bufferin precursor-, MNIO-, DUF2063- and DoxX-coding genes constitute the cores of the bufferin BGCs.

Buf1 precursors range from 30 to 835 residues and harbor the four conserved Cys residues present in the founding member (22) (Fig. 3G; Suppl. Fig. S1). It is likely however that the smallest of these proteins are truncated. Buf2 precursors are between 51 and 367 residues long, with six conserved Cys residues in 90% of cases and twelve Cys in the remainders. An AlphaFold2 model of one of the latter indicates that it consists of duplicated 6-Cys-containing domains (Fig. 3H; Suppl. Fig. S1). Approximately 16% Buf2 bufferins are predicted to harbor lipoprotein signal peptides and might therefore be membrane-anchored after Sec-dependent export. In both families we identified a few instances of large bufferin domain-containing fusion proteins, including Buf1|Cytochrom_C|FGE-sulfatase (Pfam entries of the latter two domains PF00034 and PF03781), MerC|Buf1 (Pfam entry of the former domain PF03203) and MerC|Buf2 chimeras (Fig. 3J,K). The formylglycine-generating enzymes (FGE) are cuproenzymes (40), consistent with copper-related functions. MerC-containing fusions are found in BGCs related to mercury detoxification (41), indicating that some bufferins may participate in stress responses to other metals.

### Other signal peptide-containing precursors

Proteins of the third largest cluster of the putative RiPP precursors node network (Fig. 1C) contain oxazolins (32). The newly generated hmm profile for these proteins was called ‘Buf_EGKCG/oxazolin’ (Suppl. File S2), as they are characterized by repeated Glu-Gly-Lys-Cys-Gly motifs. The majority of the approximately 2,000 proteins collected with this new signature are mostly found in *Pseudomonadota* (Fig. 3C). Analyses of their BGCs revealed that most are composed of the core RiPP-, MNIO-, DUF2063 and DoxX-coding genes in various combinations (Fig. 3F). Small numbers of BGCs harbor additional genes that code for proteins with the names oxidored_molybd (Pfam entry PF00174) and Ferric reductase (Panther Database entry PTHR11972) domains, suggesting metal-related or redox functions.

The Buf_EGKCG/oxazolin precursors are 31 to 506 residues long and are of two distinct types. Proteins of the first type contain variable numbers of 17-residue repeats with two EGKCG motifs each. An AlphaFold2-generated structural model of a mid-size protein showed a β-helix fold (Fig. 3I; Suppl. Fig. S1), with the Cys residues putative targets of the MNIO enzyme predicted to be oriented towards the inside of the helix, reminiscent of copper storage proteins (42). The longest protein of the family, from *Pseudoalteromonas piscicida,* harbors 58 Cys (strain JCM_20779, protein acc. ATD05916.1). Proteins of the second type harbor two Ca^+2^-binding EF-hand_5 domains fused to the C terminus of a short Buf_EGKCG domain devoid of 17-residue repeats and preceded by a predicted transmembrane helix (Fig. 3L). AlphaFold2 predicted that the N-terminal Buf_EGKCG domains of the second type of proteins are unstructured.

We also built an hmm profile called ‘CxxxxC’ for the fourth largest cluster of precursor proteins (black dots in Fig. 1C), which notably contain signal peptides and conserved CxxxxC motifs (Suppl. File S3). The newly built signature collected 76 proteins at < 90% sequence identity, all from *Pseudomonadati*. In addition to the core RiPP-, MNIO, DUF2063- and DoxX-coding genes, 54% of these BGCs contain genes for proteins named Lactamase_B_2 (Pfam entry PF12706), which are metalloproteins.

Last, we searched for precursors in the BGCs of the approximately 4000 MNIO enzymes of the SSN mega-cluster in Fig. 1B that have no Buf1, Buf2, Buf_EGKCG/oxazolin or CxxxxC precursors encoded next to them. We identified hundreds of small annotated ORFs immediately preceding the MNIO genes and coding for peptides containing between 1 and 18 Cys with predicted signal peptides (Suppl. Table S2). These ORFs probably represent new precursors and might correspond to several small sequence clusters of Fig. 1C. For the remaining BGCs, we performed RODEO analyses to detect other unannotated ORFs, which we subjected to SSN analyses. However, these analyses did not group them into conserved families. Which of these might be new precursors is currently difficult to establish.

### TIGR04222 proteins

Using the TIGR04222 signature, > 3500 proteins were collected at < 90% sequence identity from the nr NCBI database, most of them from Actinomycetota (Fig. 4A). Analyses of the genetic environments of the TIGR04222-coding genes revealed a sizeable proportion of BGCs devoid of MNIO genes (Fig. 4B). Furthermore, in more than 10% cases TIGR04222 genes follow MNIO genes, in contrast with bufferin BGCs. No DUF2063-coding genes were found in the TIGR04222 BGCs, some of which harbor other conserved genes of unknown function.

**Figure 4.**
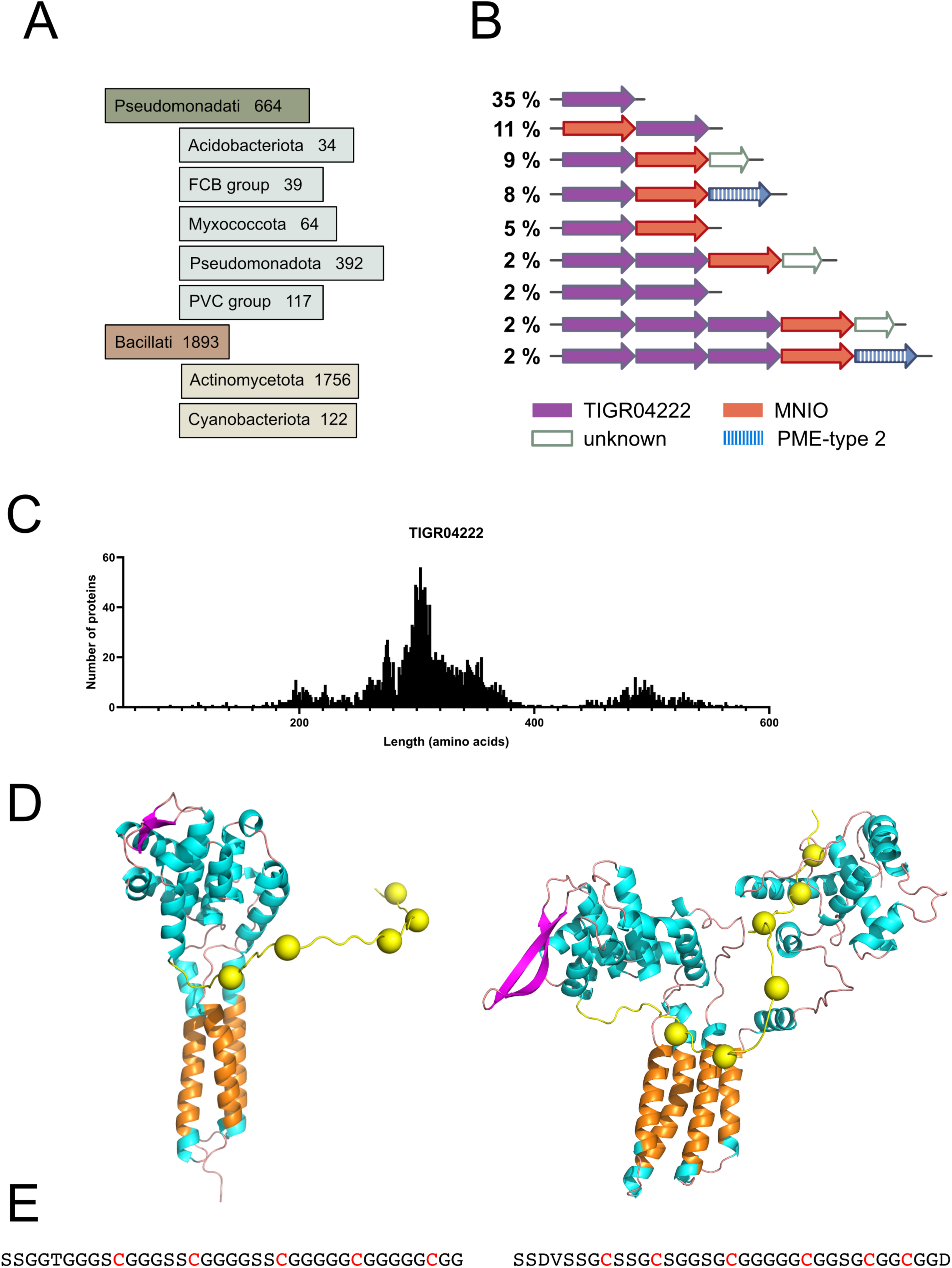
Features of TIGR04222 proteins. **A.** Taxonomic distribution. **B.** Major types of genetic organizations with their relative frequencies in the NCBI nr protein database. For this analysis only the TIGR04222 proteins harboring Cys residues were considered. See below (text and Fig. 6) for the PME-type 2 proteins. **C.** Size distribution of the TIGR04222 proteins showing the wide range of lengths. **D.** AlphaFold2 models of selected TIGR04222 proteins (WP_179806563 of *Micromonospora purpureochromogenes*, 300 residues, and ADB18956 of *Pirellula staley*i DSM 6068, 516 residues). The transmembrane segments are in orange, and the Cys residues are shown as yellow balls. E. Sequences of the C-terminal Cys-rich domains of the latter two proteins, which are predicted to be exposed to the periplasm.

The sizes of TIGR04222 proteins range from 70 to 643 residues (Fig. 4C). Alphafold2 models of selected proteins showed that they are composed of several transmembrane segments and large periplasmic domain(s), with unstructured, C-terminal Ser- and Gly-rich regions interspersed with Cys residues (Fig. 4D,E; Suppl. Fig. S1). Interestingly, these low-complexity regions are predicted to be exported to the periplasm, like the bufferins. Together with their frequent genetic association with MNIO enzymes, these features are consistent with TIGR04222 proteins also being MNIO substrates. Nevertheless, more than a quarter of them are devoid of C-proximal Cys residues, and many are not genetically associated with MNIO enzymes. Thus, only some TIGR04222 proteins might represent new MNIO-modified precursors.

### Other types of putative precursors

The SSN analysis of the MNIO enzymes has revealed that bufferins and related proteins are the majority substrates. Nevertheless, more diversity can be expected from the small, uncharacterized sequence clusters. To expand our range of new potential MNIO-modified precursors, we focused both on minor groups of MNIO enzymes, corresponding to the small sequence clusters labeled 1 to 21 in Fig. 1B, and on the non-characterized mid-size sequence clusters of putative precursors labelled I to IV in Fig. 1D.

Analyses of the BGCs that correspond to subclusters numbered 1 to 3 of the MNIO node network identified new families of precursors with Cys-containing motifs (Fig. 5A). Those genetically associated with the MNIO enzymes of the subcluster #3 harbor predicted Sec signal peptides, unlike those of clusters #1 and #2. Their BGCs are similar to those of bufferins.

**Figure 5.**
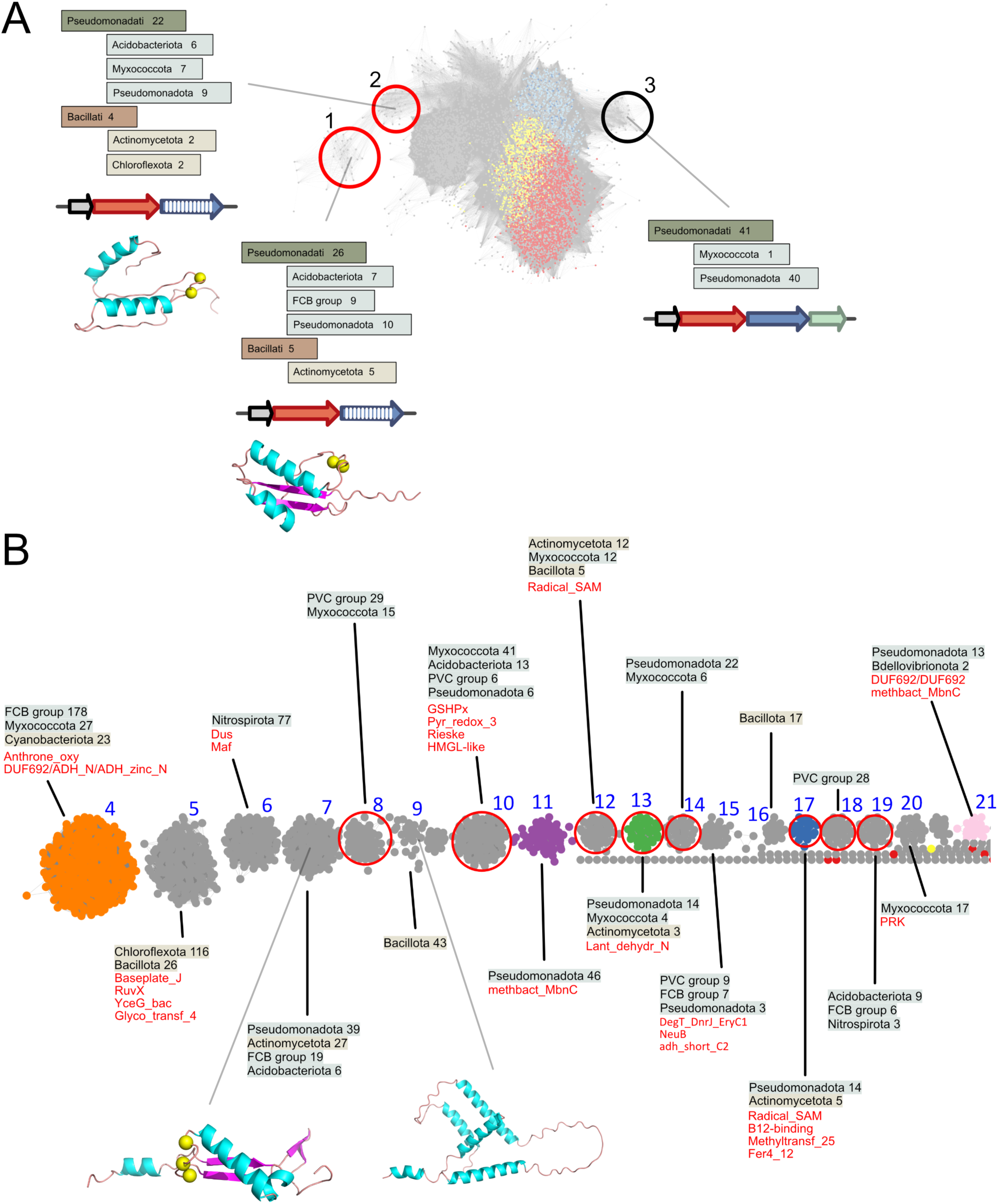
Characterization of small sequence clusters of MNIO enzymes to identify new precursors. **A, B.** Analyses of the MINO sequence clusters denoted #1, #2 and #3 (**A**) and zoom on the sequence clusters #4 to #21 (**B**) of Fig. 1B. The taxonomic groups of these MNIO enzymes and the salient features of their BGCs are shown, including the proteins frequently found encoded in these BGCs (Pfam short names in red, panel **B**). In **A**, the AlphaFold2 models of representative RiPPs are those of WP_073215760 *Massilia sp*. CF038 (cluster #1) and WP_127967852.1 *Hahella sp*. KA22 (cluster #2), with the Cys residues shown as yellow spheres. The precursors genetically associated with MNIO enzymes of sequence cluster #3 are predicted to be unstructured, and therefore no model is shown. **B.** The sequence clusters #4, #11, #13, #17 and #21 contain MNIO enzymes that modify the precursors of chryseobasins, methanobactins, pearlins, aminopyruvatides and *Vibrio* methanobactin-like RiPPs, respectively. In addition, new putative precursors were identified for MNIO enzymes of clusters #7 and #9. AlphaFold models of representative proteins are shown (WP_078758616 of *Novilysobacter spongiicola* and WP_053243439 of *Clostridium sp.,* respectively). Note that no Cys residues are present in the latter. Among these clusters of MNIO enzymes, only the BGCs of the cluster #3 (panel **A**) encode *bona fide* DUF2063 domain-containing partner proteins (called PME-type 1; see below). In contrast, the BGCs of the ten clusters circled in red (panels **A** and **B**), including those of TglH and ApyH for the biosynthesis of pearlins and aminopyruvatides, respectively, encode PME-type 2 proteins (see below).

We next analyzed the BGCs of MNIO enzymes found in clusters 4 to 21 (Fig. 5B). These BGCs are extremely diverse, even within each sequence cluster. Other than the already described precursors of chryseobasins, methanobactins, aminopyruvatides and pearlins, (23, 24, 26–28, 30), we identified potential signal peptide-less RiPP precursors in the BGCs of two other small clusters. MNIO enzymes of cluster #7 are genetically associated with putative precursors harboring a CxxC motif (Fig. 5B; Suppl. Fig. S1). Most BGCs corresponding to MNIO enzymes of cluster #9 also code for potential RiPP precursors devoid of Cys residues. Proteins with putative functions in RiPP biosynthesis were encoded in many of these BGCs (Pfam names listed in red in Fig. 5B). Thus, based on these analyses, many minor types of MNIO-modified RiPP precursors remain to be identified, but this is not straightforward as already acknowledged (43). In the absence of small ORFs it is hazardous to assign a precursor status to unknown ORFs with no distinctive features.

We also analyzed four sequence clusters of putative precursors, labeled I to IV in Fig. 1D. With 7 to 21 Cys residues, proteins of cluster #I plausibly represent a new group of precursors (Table 2). Similarly, those of cluster #IV harbor a CxxCC motif and might also be a new group of RiPP precursors. In contrast, no precursor genes were identified in the BGCs corresponding to clusters #II and #III. The genes that precede the MNIO enzyme genes in these latter two clusters code for peptidyl-tRNA hydrolases (Pfam entry PF01195), and for putative metalloproteins with a HExxH motif, respectively. Both types of proteins are thus more likely to be enzymes than precursors (28, 29, 44). In the BGCs encoding cluster #III proteins, we found genes coding for MNIO|Nitroreductase fusion proteins (Pfam entry of the latter: PF00881), suggesting the possibility of new chemical modifications.

### Identification of MNIO partner proteins

Partners of MNIO enzymes include proteins that possess DUF2063, Anthrone_oxyd or methbact_MbnC Pfam signatures. The former proteins are by far the most frequent partner proteins of MNIO enzymes (approx. 60%), and we propose to name them PME-type 1, for Partner protein of MNIO Enzyme, type 1. To identify other types of partner proteins, we analyzed the proteins coded by genes that follow the MNIO genes, based on observations that this is the most frequent gene order (Fig. 2B). Among the proteins thus collected, 72% possess the DUF2063 signature, others have Anthrone_oxyd, methbact_MbnC, DoxX or MNIO signatures, and 1700 others have no Pfam signature. SSN analyses of these latter sequences revealed that they could be sorted into several sequence clusters (Fig. 6A).

**Figure 6.**
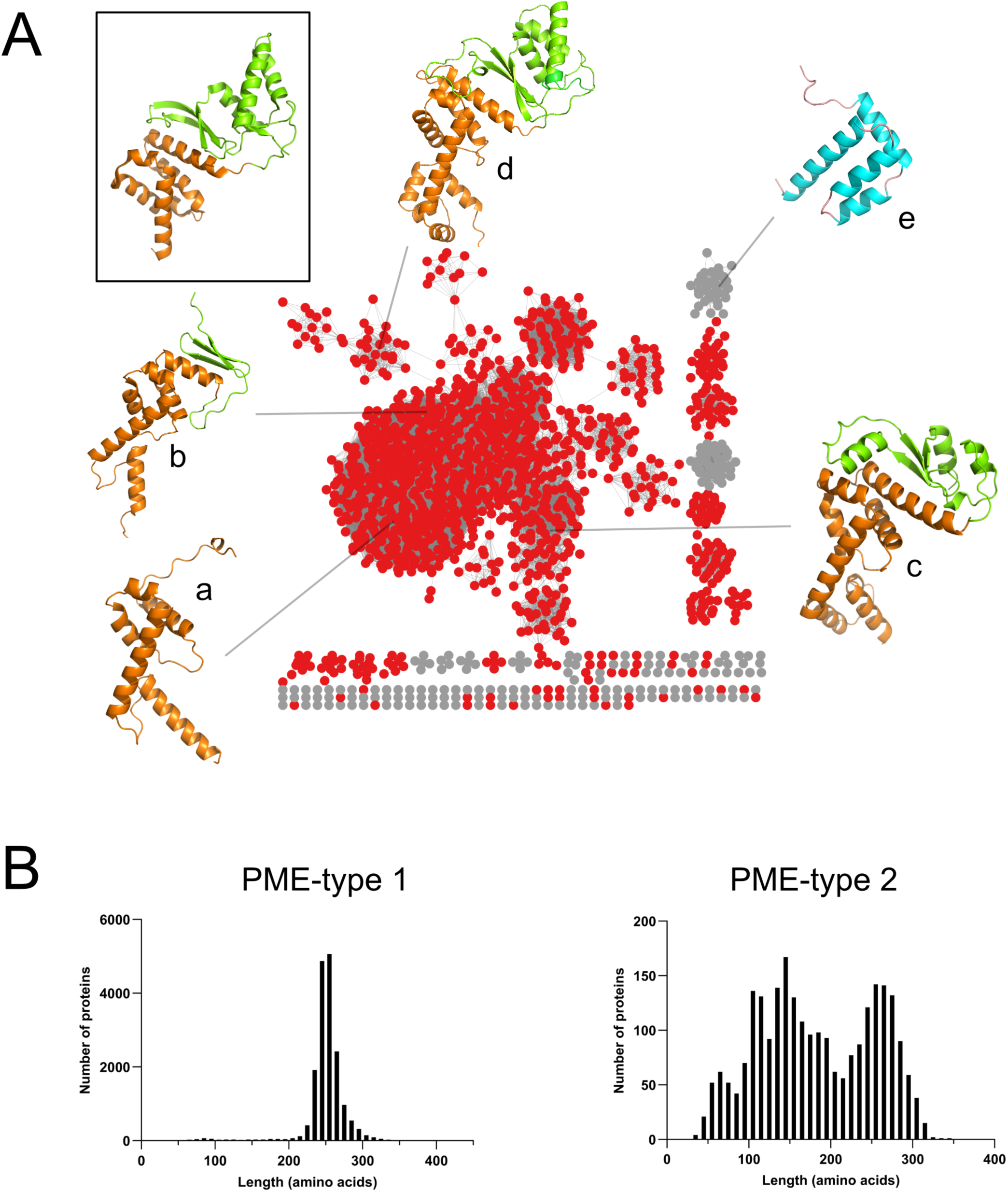
Characterization of new MNIO partner proteins. **A**. Representative node network (AST= 5) and structural models of the unknown proteins encoded by genes that follow the MNIO-coding genes. The nodes of proteins found to be structurally related to the PME-type 1 *NGO1945* protein of *Neisseria* (pdb #3dee) signature but that do not possess the Pfam DUF2063 signature are colored in red. They are hereafter called PME-type 2 proteins. The inset shows the X-ray structure of the *NGO1945* protein, with its DUF2063 and RRE domains colored orange and green, respectively. The Alphafold models a to d of representative PME-type 2 proteins found in the major sequence cluster are shown with the same color code: a, WP_033442786 of *Saccharothrix* sp. NRRL B-16314 (UNIPROT A0A7W9M5Z5, 115 residues); b, WP_091237309 of *Micromonospora matsumotoense* (UNIPROT A0A246RJP9, 158 residues); c, WP_151034108 of *Cellvibrio* sp. KY-GH-1 (UNIPROT A0A5J6PC27, 215 residues); d, WP_006975235 of *Plesiocystis pacifica* (UNIPROT A6GEU5, 276 residues). The blue model shown in e (WP_035891099 of *Legionella norrlandica*) represents a potential new type of partner proteins that belong neither to PME-type 1 nor to PME-type 2 proteins. **B.** Size distribution of the PME-type 1and PME-type 2 proteins. The majority length of 250-260 residues of PME-type 1 proteins in the left-hand graph corresponds to a fusion between a DUF2063 domain and a complete RRE-fold type domain. In the right-hand graph, the shorter proteins that are heterogeneous in length most likely do not contain full-length RRE-type domains.

To detect low-level similarity between these uncharacterized proteins and known domains, we performed HHpred analyses to identify structural homologs. For many of them the *Neisseria gonorrheae* protein NGO1945 (pdb# 3dee) and TglI (pdb#8hi7), the partner of the MNIO enzyme TglH (28) were among the best hits. NGO1945 contains both a DUF2063 domain and an RRE-type domain and is therefore a typical MNIO partner protein, although it was previously predicted to be a transcription factor (45) (Fig. 6A, inset). Alphafold2 models of a few of these uncharacterized proteins revealed that they harbor N-terminal α helix-rich domains followed in some cases by RRE-fold type domains or truncated versions thereof (Fig. 6A, models a to d). Therefore, many BGCs without *bona fide* DUF2063 genes encode DUF2063-like proteins, missed by the DUF2063 signature because of a lack of sequence similarity. We propose to call them PME-type 2 proteins. Note that PME-type 2 proteins appear to encompass a variety of subtypes, and therefore future studies will be necessary to refine their classification. The SSN analysis presented in Fig. 6A also indicated that yet additional types of partner proteins likely exist. We analyzed the largest two sequence clusters of proteins not included in PME-type 2 proteins (sequence clusters in gray in Fig. 6A). One of them contains small, all α-helical proteins with no structural homology with other proteins (Fig. 6A, model e). They might represent a minor type of MNIO partners, as most of the corresponding BGCs also encode putative precursors, notably with C-terminal CxxC motifs. The proteins of the other large non-PME type 2 cluster are coded by genes inserted between the MNIO and the *bona fide* PME-type 1 genes in otherwise classical *buf1* operons. The function of these proteins in these RiPP systems is unknown, but they are probably enzymes that introduce additional modifications.

We reanalyzed the products of unknown genes in the TIGR04222 BGCs. HHpred analyses predicted that at least 10% of TIGR04222 BGCs code for PME-type 2 proteins. Similarly, BGCs encoding MNIO enzymes from several small sequence clusters also encode PME-type 2 proteins, including those of pearlins and aminopyruvatides (circled in red in Fig. 5B).

We next determined the length distributions of the major two types of putative MNIO partner proteins. PME-type 1 proteins have a narrow size distribution corresponding to full-length DUF2063-RRE chimeras (Fig. 6B). In contrast, PME-type 2 proteins are very heterogeneous in length. The group of proteins with similar lengths as the PME-type 1, around 260 residues, most likely harbor two full-length domains. However, a large proportion of PME-type 2 proteins are too short to comprise both domains and probably consist of a DUF2063-related domain alone or followed by a short putative RRE-type truncate or variant.

### Specificity of MNIO enzymes for RiPP precursors and partner proteins

The discovery of other types of partner proteins prompted us to investigate specific associations between the subgroups of MNIO enzymes, the families of RiPP precursors, and the types of partner proteins. To facilitate visualization of these relationships, we colored the nodes of the large sequence cluster of MNIO enzymes of Fig. 1B according to the types of precursors and partner proteins encoded in the same BGCs. This confirmed that specific areas of this large cluster of MNIO enzymes correspond to specific precursor families (Fig. 7A). In addition, coloring of the nodes of MNIO enzymes as a function of their partner proteins showed that those genetically associated with Sec signal peptide-containing precursors (i.e., the Buf1, Buf2, Buf_EGKCG/oxazolins and CxxxxC families) have authentic PME-type 1 proteins (Fig. 7B). In contrast, those associated with signal peptide-less TIGR04222 proteins appear rather to have PME-type 2 partners.

**Figure 7.**
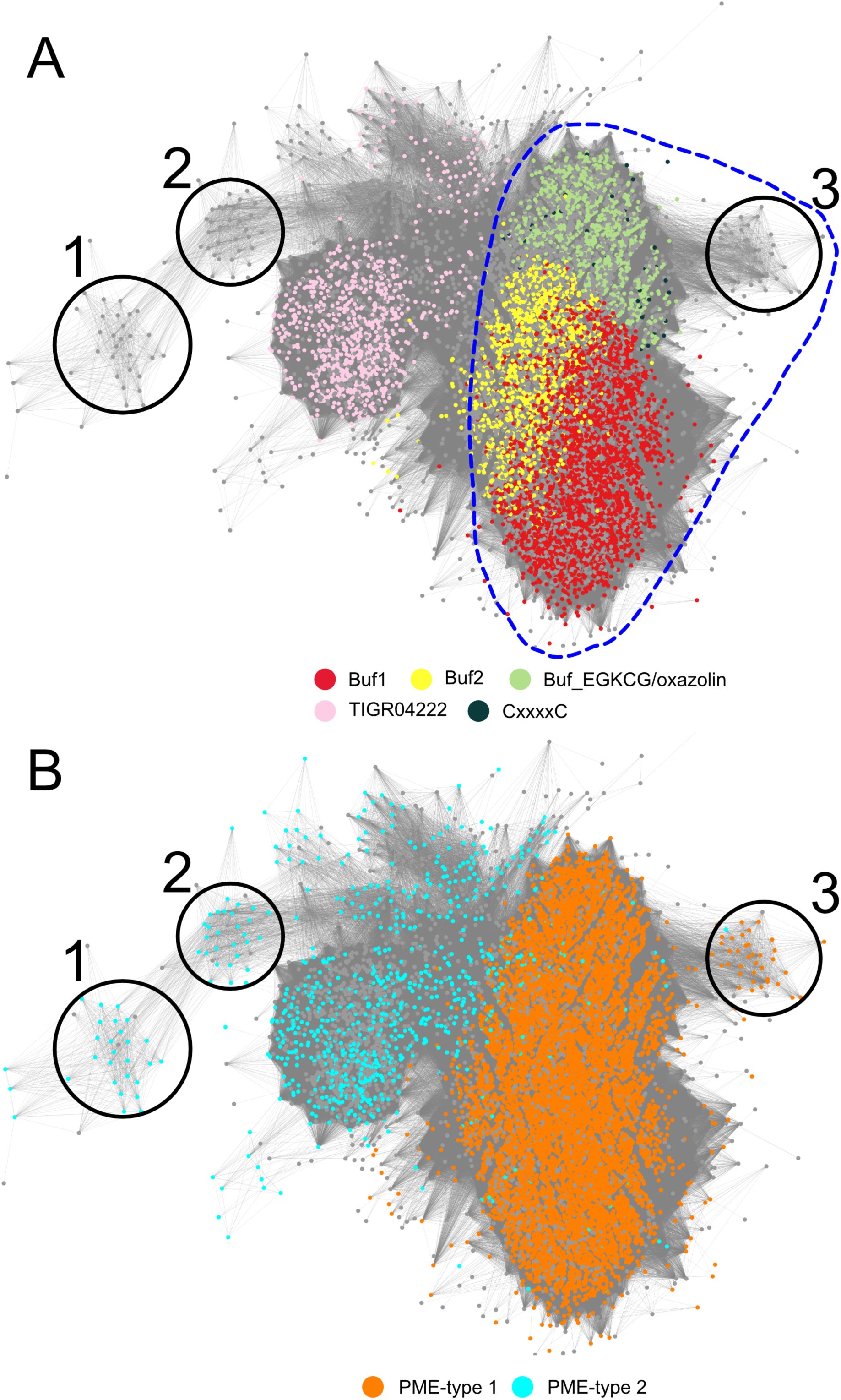
Subgroups of MNIO enzymes are associated with specific precursors and partner proteins. Members of the large sequence cluster of the node network of MNIO enzymes in Fig. 1B are colored according to their genetically linked precursors and partner proteins. In **A**, the colors of the dots represent the MNIO enzymes according to their putative precursor substrates. The pale green dots include both the Buf_EGKCG/oxazolin proteins and the Buf_EGKCG|EF-hand_5 chimeras. The precursors with signal peptides are circled with a stippled blue line. In **B**, the orange and cyan dots represent MNIO enzymes associated with PME-type 1 and with PME-type 2 partner proteins, respectively.

This correlation was confirmed for three small MNIO sequence clusters numbered 1 to 3. Thus, the precursors associated with the MNIO enzymes of clusters #1 and #2 are devoid of predicted Sec signal peptides, and the partners of their MNIO enzymes are PME-type 2 proteins (Fig. 7A,B). In contrast, the precursors associated with MNIO enzymes of cluster #3 harbor predicted Sec signal peptides, and their MNIO enzymes have *bona fide* PME-type 1 proteins. Thus, there is a correlation between the presence of a signal peptide in the precursor and the type of partner protein.

### Co-occurrence analyses

Finally, we mined > 40,000 fully assembled bacterial genomes in the NCBI database to determine the maximal numbers of occurrences of bufferin or bufferin-like BGCs in a single genome. The most extreme case is that of *Legionella pneumophila subsp. fraseri*, with five Buf1 (DUF2282), two Buf2 (BUF_6/12Cys)-, one BUF_EGKCG-, seven MNIO-, six DUF2063- and three DoxX-coding genes (Suppl. Fig. S2). The presence of bufferin precursor genes not associated with MNIO genes suggests that these precursors might be modified by MNIO enzymes encoded at a different locus or that they are not modified. However, we do not know whether all these BGCs are functional.

Our genomic analysis further revealed that very few Buf1, Buf2 or Buf_EGKCG/oxazolin precursor genes are present without MNIO and PME-type 1 genes in the same genome (Table 3). In contrast, a third of genomes harboring TIGR04222 genes are devoid of MNIO genes, revealing a looser association between these two families and further supporting the idea that TIGR04222 proteins form several subfamilies only some of which are likely to be modified by MNIO enzymes.

**Table 3.** Analysis of fully assembled bacterial genomes.

| Type of precursor | Genomes with precursor genes | Max numbers of precursor genes per genome | Numbers of genomes with precursor genes and w/o MNIO genes | Number of genomes with precursor and MNIO genes and w/o PME-type 1 genes |
| --- | --- | --- | --- | --- |
| DUF2282 | 4072 | 5 | 21 | 27 |
| Buf_6-12Cys | 864 | 4 | 0 | 1 |
| Buf_EGKCG | 2583 | 4 | 67 | 56 |
| TIGR04222 | 1372 | 5 | 490 | ND |

*ND, not determined as TIGR04222 genes are not associated with PME-type 1 genes.*

## Discussion

We explored the substrate diversity of MNIO enzymes and described several groups of potential precursors in various eubacterial taxonomic groups. We revealed that the BGCs of the largest groups of RiPP precursors are composed of a small number of core genes. This limited gene set and the overwhelming conservation of Cys-containing motifs indicate that these residues are the most likely targets of MNIO modifications, and that secondary PTMs are rare for the major precursor families. In contrast, the BGCs corresponding to the small clusters of MNIO enzymes are larger and more diverse, with many comprising genes for putative tailoring enzymes. They are likely to code for new MNIO-catalyzed chemistry, including on non-Cys residues, as well as for various secondary modifications. In fact, it is in some of those small MNIO enzyme groups that precursors were found to be modified on Asp or Asn residues or to carry additional PTMs catalyzed by other enzymes (23, 30). In many cases however, peptide precursors are difficult to identify, suggesting that some of these MNIO enzymes may modify *bona fide* proteins, which is likely the case for subsets of TIGR04222 proteins.

We have revealed that the Buf1 bufferin of *C. vibrioides* is involved in copper homeostasis and harbors Cys-derived thiooxazole groups mediating the chelation of Cu(I) and Cu(II), thereby enhancing bacterial growth in conditions of copper excess (22). Buf1 likely promotes survival of environmental bacteria subjected to copper stress in soils or water polluted by metals, notably from anthropic sources (46), or when they are engulfed by amoeba (47). Similarly, the use of copper as an antimicrobial agent in innate immunity (48) might account for the presence of bufferins in bacterial pathogens of mammals. Although we have not identified the chemical modifications carried by Buf2 bufferins, preliminary characterization of that of *C. vibrioides* has shown that it has a similar function as Buf1, its PTMs result in the same mass loss per Cys residue, and its UV/vis spectrum shows a similar 305-nm absorbance maximum (22). Furthermore, the structural models of Buf and Buf2 bufferins are related, and their associated MNIO enzymes belong to the same large sequence cluster, suggesting that they are iso-functional. These elements argue in favor of similar PTMs and functions for both bufferin families.

The PTMs and the function of a member of the MNIO-modified Buf_EGKCG/oxazolin family were recently reported (32). Similar to Buf1, it binds Cu(I) but appears to do so using Cys-derived oxazolone thioamide groups, similar to methanobactins. These proteins constitute the third largest sequence cluster in the SSN node network of RiPP precursors. Interestingly, the MNIO enzymes associated with these precursors are closely related to those associated with bufferin precursors, and their partners are PME-type 1 proteins, like for bufferins. Whether the MNIO enzymes that belong to the large sequence cluster are all iso-functional remains to be seen. Nevertheless, the discrepancy between the PTMs of bufferins and Buf_EGKCG/oxazolins is puzzling and warrants further investigations.

In contrast, TIGR04222 membrane proteins are unrelated putative precursors. Those that harbor C-proximal Cys-rich segments predicted to be exported to the periplasm may be modified by MNIO enzymes. However, one cannot speculate on the nature of their PTMs, as the MNIO enzymes and partner proteins genetically associated with TIGR04222 proteins are somewhat distant from those of bufferins. Similarly, the functions of these proteins in the producing organisms remain to be investigated. Some TIGR04222 proteins are not even associated with MNIOs and/or have no Cys residues. Given their diversity, TIGR04222 proteins are likely to have a variety of functions and to form several distinct families, only some of which may be post-translationally modified by MNIO enzymes.

Methanobactins are involved in copper acquisition and bufferins and Buf_EGKCG/oxazolins in the protection against copper. However, the diversity of the precursors and of their BGCs indicates that many putative RiPPs identified in this study likely play roles beyond adaptation to excess copper. This was already suggested by the denomination ‘*gig*’ (gold-induced genes) operons, which are actually bufferin BGCs, although their role in the protection against gold remains to be shown (49, 50). The identification of fusion proteins between bufferins and metal-binding domains such as MerC further supports a role in the homeostasis of various other metals. As for Buf_EGKC|EF-hand fusion proteins, in addition to a role in copper homeostasis they might contribute to Ca^2+^ homeostasis or signaling, as exemplified by EfhP in *Pseudomonas aeruginosa* (51, 52). Furthermore, the presence of *doxX* genes in many bufferin BGCs suggests protection from other stresses commonly encountered by bacteria. A DoxX homolog in a different genetic context was shown to protect bacteria against oxidative or sulfur stress (53). As for most other types of MNIO-modified RiPPs, their functions remain unexplored thus far.

The presence of signal peptides in a large proportion of MNIO substrates is an original feature among bacterial RiPPs. As the signal peptide directs the precursor to the Sec machinery, there must be a mechanism to avoid premature export before the installation of the PTMs. How recognition between the precursor and the MNIO-partner protein complex operates has been described for methanobactin (24) and pearlin precursors (54), neither of which have signal peptides. In this work, we found out that MNIO enzymes whose substrates are precursors with signal peptides have PME-type 1 proteins, unlike those associated with signal peptide-less precursors. It is tempting to speculate that interactions between the precursor and the PME-type 1 protein might mask the signal peptide until after installation of the PTMs, thereby preventing an early interaction with the Sec machinery. We have preliminary experimental results in our Buf1 model system supporting the idea that the signal peptide is indeed part of the recognition sequence by the BufB-BufC complex. In contrast, many BGCs harbouring TIGR04222 and MNIO genes appear not to harbor partner protein genes. Whether and how the corresponding precursors are modified remains to be established. Future studies will also reveal which MNIO enzymes can work without partner proteins. Of note, the second MNIO enzyme of methanobactin-like BGCs in *Vibrio*, MovX, appears to do so (27). Nevertheless, MovX comprises a winged helix-turn-helix domain in addition to the catalytic domain, which might fulfill a partner function.

Bufferin or Buf_EGKCG/oxazolin precursor genes are rarely present in genomes without MNIO and partner protein genes. Presumably, in these rare occurrences the peptides have activities that do not need PTMs, or they are no longer expressed or functional. We also found genomes with several bufferin-like precursor genes but a single MNIO and PME-type 1 gene pair. It is possible that the lifestyles of these bacteria have fostered RiPP expansion, and that a single MNIO-PME type 1 complex can modify several related precursors. Alternatively, only some of the precursors are modified, and the unmodified ones serve a different purpose. Further studies are needed to address these issues.

Our study suggests that MNIO enzymes modify not only peptides but also proteins that respond to the same signatures as smaller precursors but contain repeated motif or additional domains. In particular, many Cys residues are found in very large precursors of the Buf_EGKCG/oxazolin family. If these residues are all modified before export, there must be a mechanism to thread the precursor through the MNIO active site for its sequential processing, similar to the iterative mechanisms reported for several other families of RiPP precursors (17, 55, 56). The identification of BGCs encoding long precursors followed by two MNIO-coding genes indicates that more than one enzyme might be needed to extensively modify such long proteins.

## Materials and Methods

### Database searches

The non-redundant NCBI protein database was searched with various hidden Markov models (Pfam https://www.ebi.ac.uk/interpro/ (57), TIGRFAMs https://www.jcvi.org/research/tigrfams (58) and home-built HMM models), using the hmmsearch program of the HMMER3 suite (59). Each set of results were clustered by CD-HIT (60) with default parameters to remove redundant sequences sharing at least 90% sequence identity, to avoid overrepresentation of well-studied bacterial genera. The size, the domain composition, and the taxonomy were determined for each protein.

### Sequence similarity network analysis (SSN)

Sequence similarity networks were generated using the Enzyme Function Initiative Enzyme Similarity Tool (EFI-EST) (https://efi.igb.illinois.edu/efi-est/) (61). We followed the guidelines of the EFI-EST site and chose rather stringent parameters (alignment score corresponding to 40% pairwise identity), to better separate the groups of interest. The selected sequence alignment score threshold (AST) is given in each figure. Networks were visualized by Cytoscape (62) by using the organic layout. The ID numbers of all entries found in the SSN node networks are provided in Suppl. Table S3.

### Local database construction and characterization of genetic environments

A local DNA database was constituted by downloading all available bacterial genomic sequences for both Eubacteria and Archeae from the NCBI RefSeq, GenBank and WGS databases. A Python script was written to extract the genetic environments of all MNIO-coding genes by retrieving the sequences of the five genes on either side of the gene of interest in a Rodeo-type approach. The resulting table was used to collect all genes coding for proteins of interest. To define the RiPP BGCs, we imposed the following criteria. All genes encoding proteins known to be relevant to MNIO BCGs (with Pfam short names DUF2282, MbnB_TglH_ChrH (MNIO), DUF2063, DoxX, Sigma70_r4_2 (sigma factor), Sigma70_r2 (sigma factor), NrsF, or NCBIfam name methanobac_OB3b (= MbnA), or corresponding to the new signatures Buf_EGKCG/oxazolin, Buf_6/12Cys, or CxxxxC) in the vicinity of the MNIO genes were considered part of the BGC, irrespective of the intergenic distances. For unrelated genes, we set up a stringent intergenic distance criterion (maximum 10 bp from any relevant gene) for inclusion to ensure that these new genes are in operons with at least one known gene. Although somewhat arbitrary, this choice proved to provide the most information while avoiding too much background noise. For the smaller sequence clusters of MNIO enzymes, RODEO analyses were performed, and all ORFs identified were considered as part of the BCGs irrespective of the intergenic distances. The numbers of occurrences of each type of operonic structures were computed. Prediction of signal-peptides and their cleavage sites was performed using SignalP V6 (63).

### Generation of hmm profiles

We generated hmm profiles for the Buf_6/12Cys, Buf_EGKCG/oxazolin and CxxxxC families according to the procedure described in (64). Briefly, a few dozen proteins (33, 36 and 41 for Buf2, Buf_EGKCG and CxxxxC proteins, respectively) were picked from each cluster of interest of the Representative Node Network 70% displayed with Cytoscape. We chose nodes well distributed across each cluster. Only nodes with a ‘number of IDs in Rep Node’ equal to 1 were kept. Outliers with respect to length were also removed. For each family we first determined the most frequent protein size in the cluster, and we selected sequences from this major subset to generate the seed alignment. The sequences were aligned with MAFFT (L-INS-i), an iterative refinement method incorporating local pairwise alignment information. The non-conserved regions (mostly the N-and C-termini) were edited manually in Jalview, and a hmm profile was generated with this seed alignment using hmmbuild (59). We then performed new SSN analyses with all proteins collected using a given hmm profile to ensure that only *bona fide* members were retained. Outliers found in small, isolated SSN clusters were discarded. The hmm profile files are provided in Supplementary materials (Files S1 to S3).

### Characterization of unknown proteins

To gain insight into proteins of interest devoid of Pfam or TIGRfam domain signatures, we picked several proteins in each cluster of the representative node network and first looked for Interpro signatures. If no Interpro signature was identified, we searched for related proteins using HHpred (65) and selected the best hits. If necessary, we generated AlphaFold2 models of selected members and used DALI (66) to find related structures in the RCSB Protein Data Bank (67).

### Analysis of assembled bacterial genomes

The proteomes of the fully assembled bacterial genomes were downloaded from NCBI (https://www.ncbi.nlm.nih.gov/assembly). The bacterial proteomes were searched with the hmmscan program of the HMMER3 suite (59) using the hmm profile signatures DUF2282, DUF692, DUF2063, DoxX, EF-hand_5, Buf_6/12_Cys, Buf_EGKCG, and TIGR04222 using home-made scripts. The domains of all the corresponding proteins were determined, and the occurrences of each type of domain of interest were computed (Table S4).

## Data summary

A local DNA database was constituted by downloading all available bacterial genomic sequences for both Eubacteria and Archeae from the NCBI RefSeq, GenBank and WGS databases. Protein sequences were collected from the non-redundant NCBI protein database. We also mined the NCBI assembled genomes database. Accession numbers of all the protein sequences used in this work and all genome identification numbers are provided in Supplementary materials in the form of Excel tables.

