## Supplementary material for "Multinuclear non-heme iron dependent oxidative enzymes: Landscape of their substrates, partner proteins and biosynthetic gene clusters": Suppl. Tables S1, S2; Fig; S1, S2; Files S1-S3

**Table S1. Protein domains genetically associated with DUF692 genes**

| Pfam_TIGRfam_Name | Count | % co-occurrence | |
|---|---|---|---|
| DUF692 | 13976 | 102 | |
| DUF2063 | 9696 | 71 | |
| DoxX | 5414 | 40 | |
| DUF2282 | 5268 | 39 | |
| TIGR02937 | 3116 | 23 | RNA polymerase sigma-70 family |
| Sigma70_r2 | 3078 | 23 | |
| Sigma70_r4_2 | 2878 | 21 | |
| ABC_tran | 2708 | 20 | |
| Response_reg | 2394 | 18 | |
| HATPase_c | 1694 | 12 | |
| NrsF | 1596 | 12 | |
| HTH_1 | 1471 | 11 | |
| LysR_substrate | 1466 | 11 | |
| Pyr_redox_2 | 1442 | 11 | |
| TIGR04222 | 1438 | 11 | |
| MFS_1 | 1338 | 10 | |
| BPD_transp_1 | 1046 | 8 | |
| HisKA | 1024 | 8 | |
| TetR_N | 917 | 7 | |
| Pyr_redox_dim | 908 | 7 | |
| HTH_18 | 850 | 6 | |
| GGDEF | 846 | 6 | |
| TIGR00254 | 810 | 6 | |
| EamA | 795 | 6 | |
| HAMP | 760 | 6 | |
| CBS | 748 | 5 | |

*The absolute numbers and percentages of genetic association between these protein domains and MNIO enzymes are provided. The list includes only the proteins or domains coded in the MNIO loci at frequencies of at least 5%. A proprotion greater than 100% is explained by the fact that tandem MNIO genes are present in some BGCs. In red are the domain signatures already known to be found in MNIO-encoding BGCs. Note that not all genetically associated proteins are necessarily involved in RiPP biosynthesis. In particular, the signatures 'ABC_tran', 'Response_reg' and 'HATPAse_c' were not considered relevant, because most bacterial genomes harbor numerous genes coding for paralogs of ABC transporters and two-component systems. The MbnC signature was not found in significant proportions because the methanobactins BGCs represent a very small number of all BGCs collected.*

**Table S2. Potential precursors genetically associated with MNIO enzymes of the largest sequence cluster shown in Fig. 1B.**

| Number of Cys | counts |
|---|---|
| 1 Cys | 27 |
| 2 Cys | 93 |
| 3 Cys | 217 |
| 4 Cys | 107 |
| 5 Cys | 92 |
| 6 Cys | 92 |
| 7 Cys | 39 |
| 8 Cys | 37 |
| 9 Cys | 32 |
| 10 Cys | 17 |
| 11 Cys | 14 |
| 12 Cys | 4 |
| 13 Cys | 3 |
| 14 Cys | 0 |
| 15 Cys | 2 |
| Total | 776 |

*Only putative precursors encoded next to MNIO genes and that do not belong to the previously described families were included.*

**Table S3. Accession numbers of all proteins identified in this work (Excel file)**

**Table S4. Analysis of complete genomes: genome identification, accession numbers of the proteins of interest, and numbers of paralogs in genome (Excel file)**

Fig3G

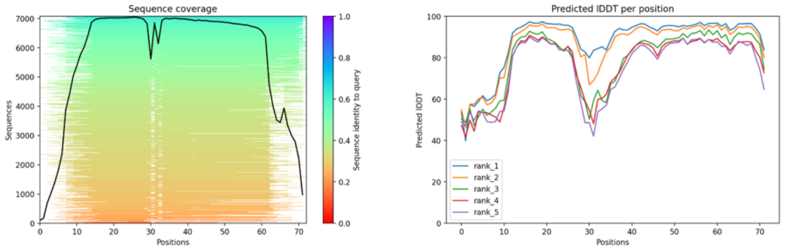


Fig3H_1

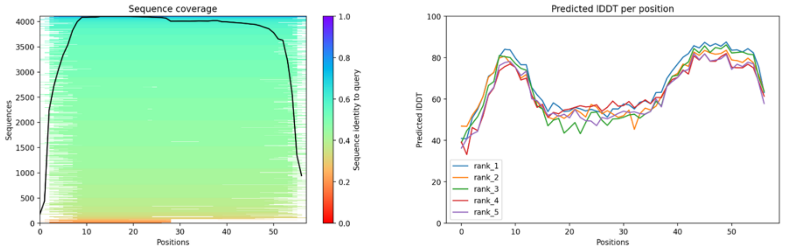


Fig3H_2

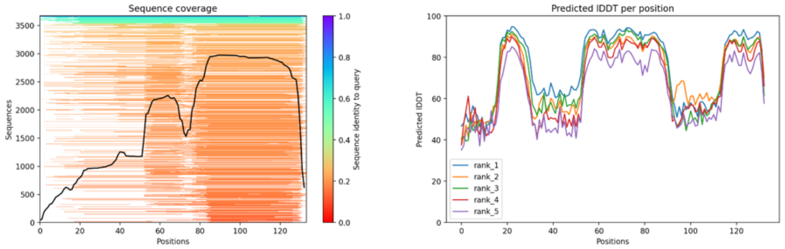


Fig3I

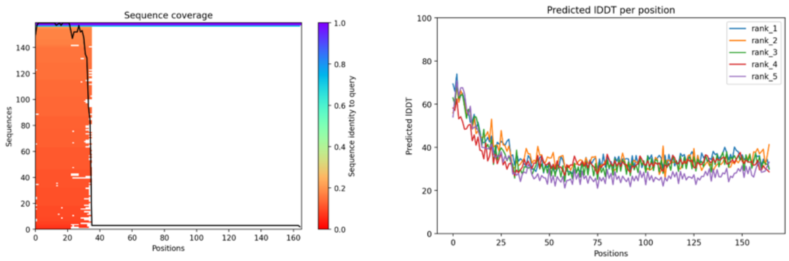


Fig4D_1

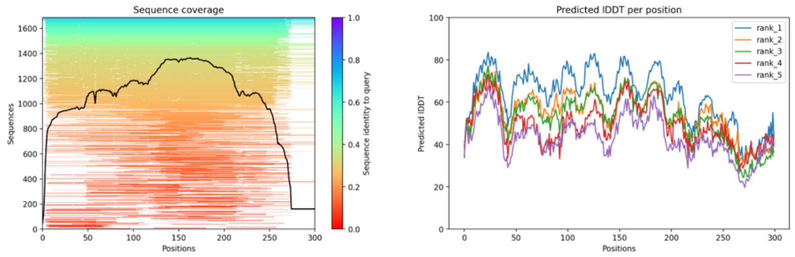

Fig4D_2

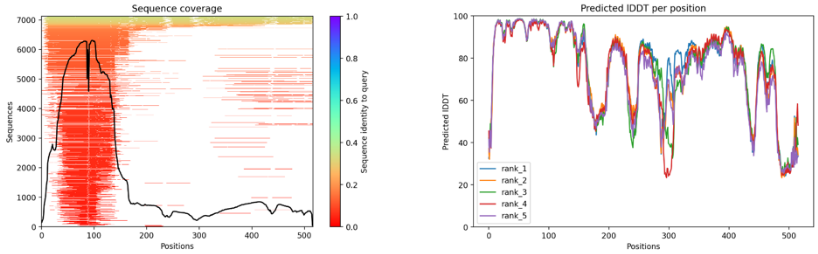


Fig5A_1

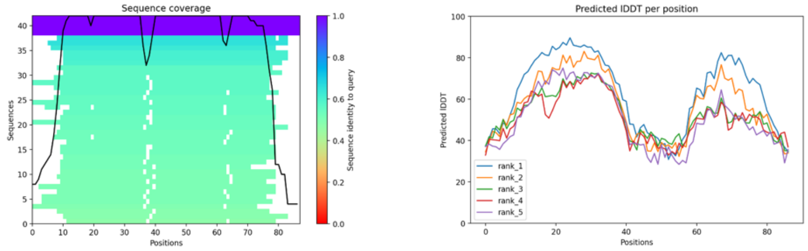


Fig5A_2

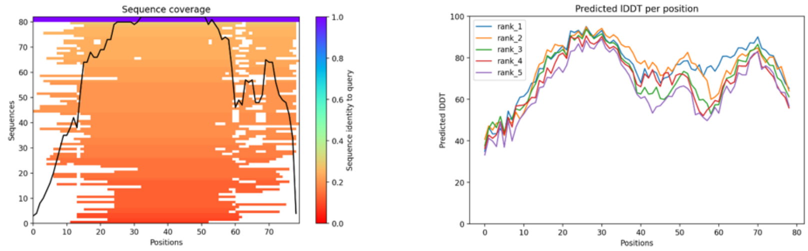


Fig5B_1

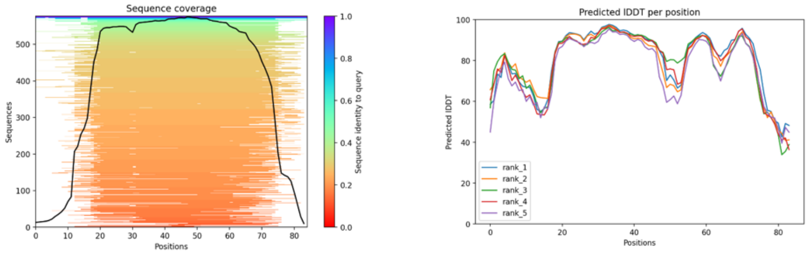


Fig5B_2

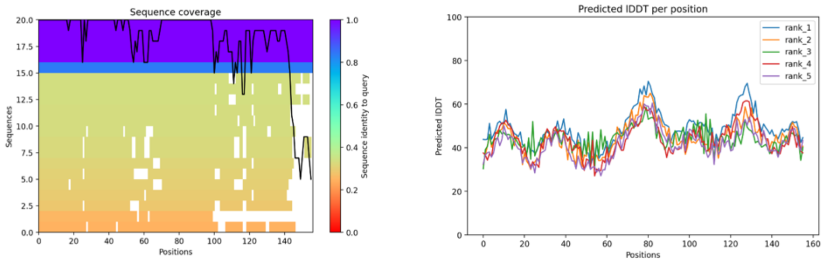

Fig6a

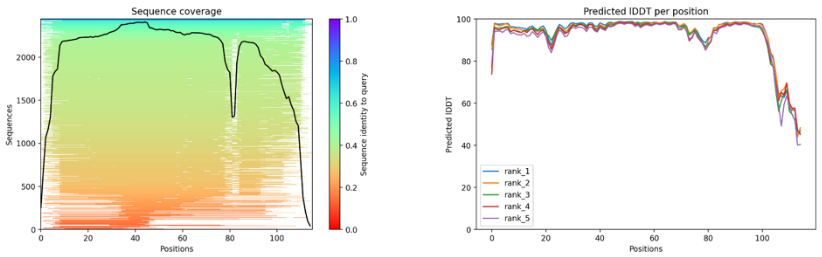


Fig6b

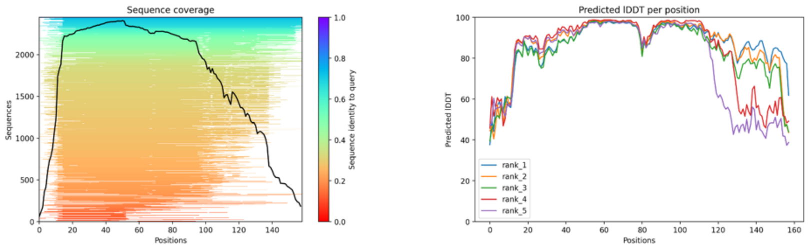


Fig6c

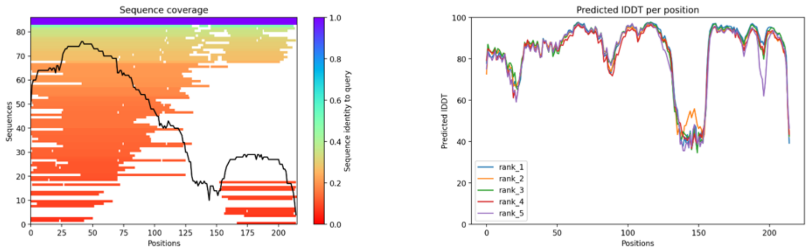


Fig6d

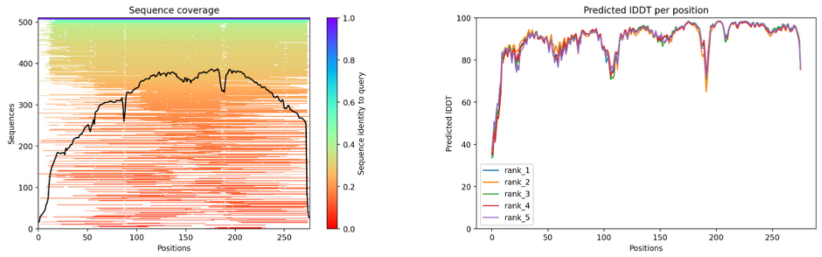


Fig6e

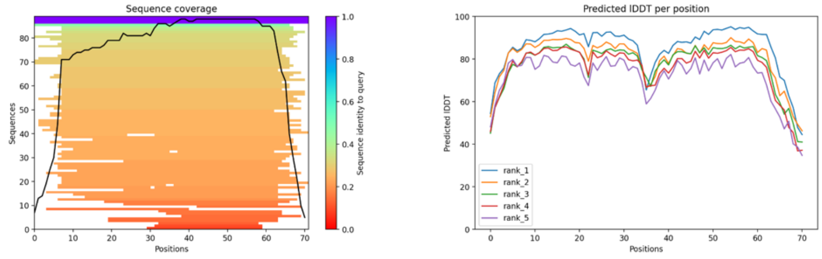


**Fig. S1. Analyses of the structural models.** The sequence coverage and the predicted local distance difference test scores for all positions in the sequences are provided for the Alphafold models shown in the main figures, with the numbers corresponding to those of the figures.

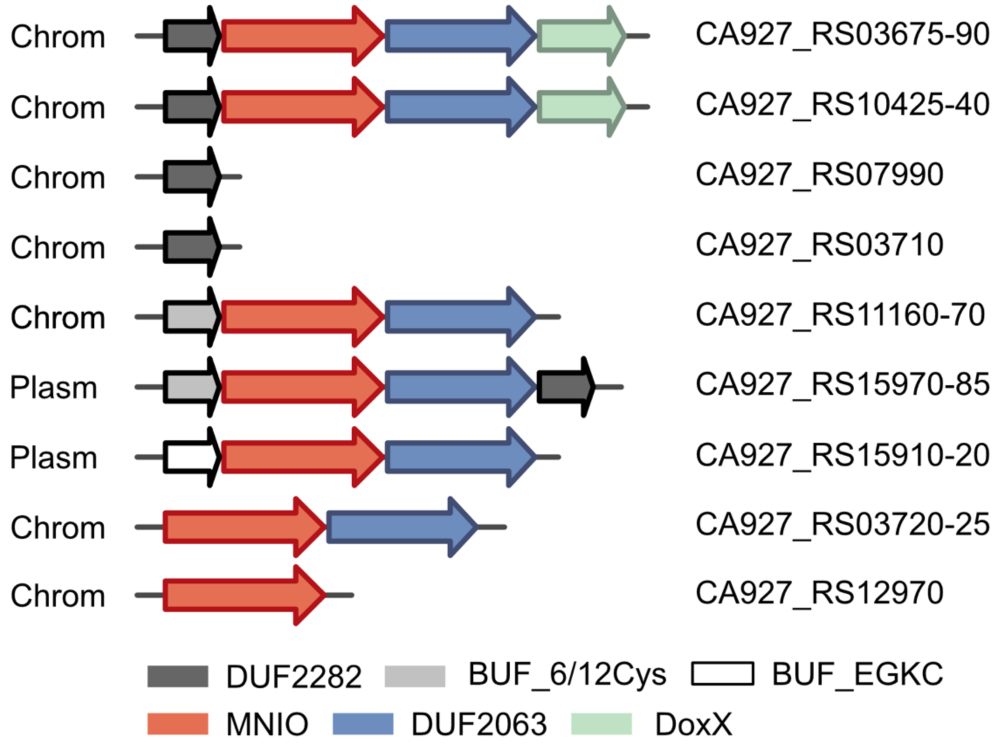


**Figure S2. Bufferin-type BGCs in the genome of *Legionella pneumophila subsp fraseri*.** The locus tags are provided at the right. Two BGCs are found on a mega plasmid. No genes coding for ECF sigma factors and anti-sigma proteins were found, and no other conserved genes were identified. Note that the bufferin associated with locus tag CA927_RS12970 is encoded by a truncated gene (not shown), suggesting loss of function of this BGC. The BGC of CA927_RS07990 also contains genes coding for a YceI protein (small beta barrel protein described to bind hydrophobic molecules) and a cytochrome B protein. The RiPP precursor of the CA927_RS03710 locus is in translational coupling with a guanylate cyclase domain. 'Chrom' and 'Plasm' indicate the location (on the chromosome or on a plasmid) of the various BGCs.

**Files S1-S3. HMM profiles for the new families defined in this work (.hmm files)**

```
HMMER3/f [3.3.2 | Nov 2020]
NAME  Buf2
LENG  50
ALPH  amino
RF    no
MM    no
CONS  yes
CS    no
MAP   yes
DATE  Thu Jun 15 08:38:42 2023
NSEQ  33
EFFN  1.905396
CKSUM 2795863552
STATS LOCAL MSV       -9.0448  0.71916
STATS LOCAL VITERBI   -9.4278  0.71916
STATS LOCAL FORWARD   -4.1249  0.71916
HMM          A        C        D        E        F        G        H
I        K        L        M        N        P        Q        R
S        T        V        W        Y
            m->m     m->i     m->d     i->m     i->i     d->m     d-
>d
  COMPO   2.48064  2.33328  3.16746  2.73946  4.10239  2.04229
3.77237  3.61096  2.32331  3.29754  4.01714  2.84017  4.00921
2.99974  3.23981  2.59035  2.85021  3.10638  5.18199  4.22241
          2.68618  4.42225  2.77519  2.73123  3.46354  2.40513
3.72494  3.29354  2.67741  2.69355  4.24690  2.90347  2.73739
3.18146  2.89801  2.37887  2.77519  2.98518  4.58477  3.61503
          0.01518  4.59089  5.31324  0.61958  0.77255  0.00000
*
      1   2.49722  5.19031  2.77800  2.23539  4.51545  3.50846
3.69882  3.98720  1.75400  3.48684  3.50557  2.98411  3.56312
2.57387  2.92926  2.52704  2.40316  3.57030  5.63497  4.23682      1
k - - -
          2.68618  4.42225  2.77519  2.73123  3.46354  2.40513
3.72494  3.29354  2.67741  2.69355  4.24690  2.90347  2.73739
3.18146  2.89801  2.37887  2.77519  2.98518  4.58477  3.61503
          0.01518  4.59089  5.31324  0.61958  0.77255  0.48576
0.95510
      2   2.76419  4.42140  3.79286  3.26706  3.57993  2.45058
2.97977  2.72792  3.18038  2.66624  3.58765  3.60614  4.16430
3.49409  3.47897  3.03339  3.02469  1.32280  5.05993  3.82094      2
v - - -
          2.68618  4.42225  2.77519  2.73123  3.46354  2.40513
3.72494  3.29354  2.67741  2.69355  4.24690  2.90347  2.73739
3.18146  2.89801  2.37887  2.77519  2.98518  4.58477  3.61503
          0.01518  4.59089  5.31324  0.61958  0.77255  0.48576
0.95510
      3   2.26304  5.23729  2.99848  2.22323  4.57557  3.34406
2.36876  4.04726  1.86718  3.53292  4.28325  2.77800  3.92468
2.58293  2.78494  2.73274  2.99528  3.62504  5.66721  4.27381      3
k - - -
          2.68618  4.42225  2.77519  2.73123  3.46354  2.40513
3.72494  3.29354  2.67741  2.69355  4.24690  2.90347  2.73739
3.18146  2.89801  2.37887  2.77519  2.98518  4.58477  3.61503
```

```
          0.01518  4.59089  5.31324  0.61958  0.77255  0.48576
0.95510
      4   3.67756  0.20013  5.24063  5.17146  5.10198  4.09652
5.67934  4.45504  5.04449  4.28990  5.44847  5.00596  4.84213
5.35764  5.02117  3.94396  4.20188  4.12728  6.27560  5.36666      4
C - - -
          2.68618  4.42225  2.77519  2.73123  3.46354  2.40513
3.72494  3.29354  2.67741  2.69355  4.24690  2.90347  2.73739
3.18146  2.89801  2.37887  2.77519  2.98518  4.58477  3.61503
          0.01518  4.59089  5.31324  0.61958  0.77255  0.48576
0.95510
      5   2.35003  4.68630  3.30810  2.56407  3.56674  3.61897
3.58624  3.08698  2.25856  2.81622  3.06153  2.89271  4.00452
2.91363  3.15251  2.52945  2.86581  2.86934  4.36182  2.72610      5
k - - -
          2.68618  4.42225  2.77519  2.73123  3.46354  2.40513
3.72494  3.29354  2.67741  2.69355  4.24690  2.90347  2.73739
3.18146  2.89801  2.37887  2.77519  2.98518  4.58477  3.61503
          0.01518  4.59089  5.31324  0.61958  0.77255  0.48576
0.95510
      6   3.78144  5.42839  4.59014  4.59699  5.62003  0.16254
5.60919  5.45342  4.86720  4.98412  6.00454  4.75794  4.83053
5.13592  4.96969  3.98112  4.31173  4.84456  6.46989  5.76432      6
G - - -
          2.68618  4.42225  2.77519  2.73123  3.46354  2.40513
3.72494  3.29354  2.67741  2.69355  4.24690  2.90347  2.73739
3.18146  2.89801  2.37887  2.77519  2.98518  4.58477  3.61503
          0.01518  4.59089  5.31324  0.61958  0.77255  0.48576
0.95510
      7   2.33212  4.25334  4.46915  3.88870  3.40352  3.34934
4.34098  1.75882  3.75005  2.09452  3.39112  4.03531  4.35042
3.95411  3.88871  3.08106  3.06932  1.46932  4.95700  3.76768      7
v - - -
          2.68618  4.42225  2.77519  2.73123  3.46354  2.40513
3.72494  3.29354  2.67741  2.69355  4.24690  2.90347  2.73739
3.18146  2.89801  2.37887  2.77519  2.98518  4.58477  3.61503
          0.01518  4.59089  5.31324  0.61958  0.77255  0.48576
0.95510
      8   3.28632  5.48576  2.87501  2.78228  4.40080  3.67547
3.29711  4.51669  3.06425  4.01136  4.92412  0.68107  4.29308
3.43908  3.45282  3.26189  3.60976  4.11591  5.76558  4.25067      8
N - - -
          2.68618  4.42225  2.77519  2.73123  3.46354  2.40513
3.72494  3.29354  2.67741  2.69355  4.24690  2.90347  2.73739
3.18146  2.89801  2.37887  2.77519  2.98518  4.58477  3.61503
          0.01518  4.59089  5.31324  0.61958  0.77255  0.48576
0.95510
      9   1.93453  5.15872  2.74617  2.40346  4.48609  3.23520
3.75798  3.94482  2.38607  3.47437  4.24560  2.84478  3.93748
2.87610  3.02005  1.74092  2.81281  3.54856  5.64872  4.26169      9
s - - -
          2.68618  4.42225  2.77519  2.73123  3.46354  2.40513
3.72494  3.29354  2.67741  2.69355  4.24690  2.90347  2.73739
3.18146  2.89801  2.37887  2.77519  2.98518  4.58477  3.61503
```

```
          0.01518  4.59089  5.31324  0.61958  0.77255  0.48576
0.95510
     10   3.67756  0.20013  5.24063  5.17146  5.10198  4.09652
5.67934  4.45504  5.04449  4.28990  5.44847  5.00596  4.84213
5.35764  5.02117  3.94396  4.20188  4.12728  6.27560  5.36666      10
C - - -
          2.68618  4.42225  2.77519  2.73123  3.46354  2.40513
3.72494  3.29354  2.67741  2.69355  4.24690  2.90347  2.73739
3.18146  2.89801  2.37887  2.77519  2.98518  4.58477  3.61503
          0.01518  4.59089  5.31324  0.61958  0.77255  0.48576
0.95510
     11   2.68284  5.13620  3.26118  2.83747  4.77205  3.08340
3.98231  4.18419  0.89183  3.70626  4.52200  3.29400  4.12245
3.12514  2.86174  2.73006  3.24338  3.76326  5.83313  4.55130      11
k - - -
          2.68618  4.42225  2.77519  2.73123  3.46354  2.40513
3.72494  3.29354  2.67741  2.69355  4.24690  2.90347  2.73739
3.18146  2.89801  2.37887  2.77519  2.98518  4.58477  3.61503
          0.01518  4.59089  5.31324  0.61958  0.77255  0.48576
0.95510
     12   2.74630  4.67704  3.83043  3.79504  5.10362  0.47259
4.97474  4.68202  4.09027  4.35071  5.17751  3.87341  4.19934
4.31268  4.32688  2.58274  3.29666  3.94175  6.38615  5.22646      12
G - - -
          2.68618  4.42225  2.77519  2.73123  3.46354  2.40513
3.72494  3.29354  2.67741  2.69355  4.24690  2.90347  2.73739
3.18146  2.89801  2.37887  2.77519  2.98518  4.58477  3.61503
          0.01518  4.59089  5.31324  0.61958  0.77255  0.48576
0.95510
     13   2.75147  5.06836  3.08706  2.52537  4.34990  3.55179
3.33812  3.78566  2.25428  3.15419  3.47011  3.05388  3.94153
1.79015  2.80159  2.59420  2.25413  3.42498  5.53636  4.18058      13
q - - -
          2.68618  4.42225  2.77519  2.73123  3.46354  2.40513
3.72494  3.29354  2.67741  2.69355  4.24690  2.90347  2.73739
3.18146  2.89801  2.37887  2.77519  2.98518  4.58477  3.61503
          0.01518  4.59089  5.31324  0.61958  0.77255  0.48576
0.95510
     14   2.26198  4.63067  3.55427  3.27334  4.83967  1.53268
4.48091  4.30498  3.42744  3.92634  4.71272  2.94101  4.04303
3.69702  3.81135  1.16481  3.08882  3.68709  6.11704  4.86531      14
s - - -
          2.68618  4.42225  2.77519  2.73123  3.46354  2.40513
3.72494  3.29354  2.67741  2.69355  4.24690  2.90347  2.73739
3.18146  2.89801  2.37887  2.77519  2.98518  4.58477  3.61503
          0.01518  4.59089  5.31324  0.61958  0.77255  0.48576
0.95510
     15   1.87565  5.18916  2.32195  2.12896  4.50736  2.86017
3.72992  3.97334  2.49191  3.48759  3.61509  2.85763  3.92248
2.84087  2.98310  2.40354  2.99101  3.56768  5.64907  4.25384      15
a - - -
          2.68618  4.42225  2.77519  2.73123  3.46354  2.40513
3.72494  3.29354  2.67741  2.69355  4.24690  2.90347  2.73739
3.18146  2.89801  2.37887  2.77519  2.98518  4.58477  3.61503
```

```
          0.06243  4.59089  2.98821  0.61958  0.77255  0.48576
0.95510
     16   3.59424  0.22091  5.16354  5.08244  5.00898  4.02900
5.59708  4.34179  4.94911  4.18587  5.34181  4.92190  4.77376
5.26485  4.93590  3.86110  4.11561  4.02043  6.20378  5.27605      16
C - - -
          2.68618  4.42225  2.77519  2.73123  3.46354  2.40513
3.72494  3.29354  2.67741  2.69355  4.24690  2.90347  2.73739
3.18146  2.89801  2.37887  2.77519  2.98518  4.58477  3.61503
          0.01591  4.54438  5.26672  0.61958  0.77255  0.56747
0.83693
     17   2.29253  5.14181  3.07027  2.58175  4.54843  2.78953
3.79906  3.98780  1.31276  3.51160  4.29920  3.09630  3.98786
2.74433  2.88257  2.64483  3.07126  3.59358  5.66899  4.32362      17
k - - -
          2.68618  4.42225  2.77519  2.73123  3.46354  2.40513
3.72494  3.29354  2.67741  2.69355  4.24690  2.90347  2.73739
3.18146  2.89801  2.37887  2.77519  2.98518  4.58477  3.61503
          0.01591  4.54438  5.26672  0.61958  0.77255  0.56747
0.83693
     18   2.30284  4.47177  3.89397  3.54006  4.67685  2.36072
4.57462  4.10569  3.56493  3.77576  4.57981  3.65847  4.00963
3.83751  3.88606  1.82458  1.02202  3.52206  6.00835  4.80817      18
t - - -
          2.68618  4.42225  2.77519  2.73123  3.46354  2.40513
3.72494  3.29354  2.67741  2.69355  4.24690  2.90347  2.73739
3.18146  2.89801  2.37887  2.77519  2.98518  4.58477  3.61503
          0.01591  4.54438  5.26672  0.61958  0.77255  0.43786
1.03680
     19   1.11079  4.64437  3.50464  3.02129  4.28218  2.47198
4.12990  3.68022  2.67351  3.33813  4.17078  3.39713  4.02010
3.34154  3.40681  2.61475  2.73924  3.02845  5.61090  4.34283      19
a - - -
          2.68618  4.42225  2.77519  2.73123  3.46354  2.40513
3.72494  3.29354  2.67741  2.69355  4.24690  2.90347  2.73739
3.18146  2.89801  2.37887  2.77519  2.98518  4.58477  3.61503
          0.01518  4.59089  5.31324  0.61958  0.77255  0.48576
0.95510
     20   2.74328  5.24765  2.25632  2.13831  4.58753  3.13454
3.33602  4.07175  2.15814  3.55061  4.29082  2.63429  3.90395
2.80064  2.83157  2.31795  2.41133  3.63573  5.68235  4.27130      20
e - - -
          2.68619  4.42226  2.77521  2.73125  3.46355  2.40506
3.72496  3.29355  2.67742  2.69356  4.24691  2.90348  2.73741
3.18148  2.89802  2.37880  2.77521  2.98520  4.58478  3.61504
          0.04229  3.31082  5.31324  0.64354  0.74534  0.48576
0.95510
     21   2.87821  5.27035  2.92830  2.51770  4.59368  3.56025
2.30701  4.06658  2.32327  3.58015  4.36484  1.67848  4.00485
2.93726  2.98723  2.14989  3.11850  3.67109  5.72775  4.34575      23
n - - -
          2.68618  4.42225  2.77519  2.73123  3.46354  2.40513
3.72494  3.29354  2.67741  2.69355  4.24690  2.90347  2.73739
3.18146  2.89801  2.37887  2.77519  2.98518  4.58477  3.61503
```

```
          0.01518  4.59089  5.31324  0.61958  0.77255  0.48576
0.95510
     22   1.89241  5.26465  2.48409  2.11369  4.59528  3.17227
3.72999  4.07576  2.31116  3.56464  4.31441  2.97844  3.92441
2.52555  2.98594  2.18295  3.01011  3.64858  5.70576  4.29489      24
a - - -
          2.68618  4.42225  2.77519  2.73123  3.46354  2.40513
3.72494  3.29354  2.67741  2.69355  4.24690  2.90347  2.73739
3.18146  2.89801  2.37887  2.77519  2.98518  4.58477  3.61503
          0.01518  4.59089  5.31324  0.61958  0.77255  0.48576
0.95510
     23   3.15157  0.52329  4.13434  2.79783  4.53252  3.79157
4.84363  3.76076  3.82326  3.62217  4.61850  4.11590  4.47163
4.23434  4.03622  3.37522  3.58207  3.46084  5.89154  4.74508      25
C - - -
          2.68618  4.42225  2.77519  2.73123  3.46354  2.40513
3.72494  3.29354  2.67741  2.69355  4.24690  2.90347  2.73739
3.18146  2.89801  2.37887  2.77519  2.98518  4.58477  3.61503
          0.01518  4.59089  5.31324  0.61958  0.77255  0.48576
0.95510
     24   2.21161  4.86627  3.19369  2.63562  3.42390  3.14157
3.79342  3.49589  1.52058  3.11948  3.31854  3.14142  3.97650
2.85411  3.03104  2.79939  2.97515  3.19268  5.38407  4.06962      26
k - - -
          2.68618  4.42225  2.77519  2.73123  3.46354  2.40513
3.72494  3.29354  2.67741  2.69355  4.24690  2.90347  2.73739
3.18146  2.89801  2.37887  2.77519  2.98518  4.58477  3.61503
          0.01518  4.59089  5.31324  0.61958  0.77255  0.48576
0.95510
     25   3.78144  5.42839  4.59014  4.59699  5.62003  0.16254
5.60919  5.45342  4.86720  4.98412  6.00454  4.75794  4.83053
5.13592  4.96969  3.98112  4.31173  4.84456  6.46989  5.76432      27
G - - -
          2.68618  4.42225  2.77519  2.73123  3.46354  2.40513
3.72494  3.29354  2.67741  2.69355  4.24690  2.90347  2.73739
3.18146  2.89801  2.37887  2.77519  2.98518  4.58477  3.61503
          0.01518  4.59089  5.31324  0.61958  0.77255  0.48576
0.95510
     26   2.88050  5.20490  3.21284  2.62647  4.52329  3.64321
3.11856  3.95636  1.72129  3.24147  3.48161  3.13619  4.02427
1.59469  2.72575  2.87017  3.09565  3.58981  5.61097  4.28565      28
q - - -
          2.68618  4.42225  2.77519  2.73123  3.46354  2.40513
3.72494  3.29354  2.67741  2.69355  4.24690  2.90347  2.73739
3.18146  2.89801  2.37887  2.77519  2.98518  4.58477  3.61503
          0.01518  4.59089  5.31324  0.61958  0.77255  0.48576
0.95510
     27   2.55665  5.01785  3.08675  3.01830  4.88652  3.48483
4.45172  4.41696  3.39839  4.08057  4.96579  0.63771  4.20633
3.68765  3.78723  3.04677  3.40555  3.90501  6.17883  4.82298      29
N - - -
          2.68618  4.42225  2.77519  2.73123  3.46354  2.40513
3.72494  3.29354  2.67741  2.69355  4.24690  2.90347  2.73739
3.18146  2.89801  2.37887  2.77519  2.98518  4.58477  3.61503
```

          0.01518  4.59089  5.31324  0.61958  0.77255  0.48576
0.95510
     28   1.88929  5.11697  2.84375  2.24928  4.43725  3.31817
3.77190  3.88664  2.55415  3.43351  4.21475  3.03397  3.94548
2.71846  3.03840  1.72593  3.00992  3.31935  5.62289  4.24622      30
s - - -
          2.68618  4.42225  2.77519  2.73123  3.46354  2.40513
3.72494  3.29354  2.67741  2.69355  4.24690  2.90347  2.73739
3.18146  2.89801  2.37887  2.77519  2.98518  4.58477  3.61503
          0.01518  4.59089  5.31324  0.61958  0.77255  0.48576
0.95510
     29   3.67756  0.20013  5.24063  5.17146  5.10198  4.09652
5.67934  4.45504  5.04449  4.28990  5.44847  5.00596  4.84213
5.35764  5.02117  3.94396  4.20188  4.12728  6.27560  5.36666      31
C - - -
          2.68618  4.42225  2.77519  2.73123  3.46354  2.40513
3.72494  3.29354  2.67741  2.69355  4.24690  2.90347  2.73739
3.18146  2.89801  2.37887  2.77519  2.98518  4.58477  3.61503
          0.01518  4.59089  5.31324  0.61958  0.77255  0.48576
0.95510
     30   3.35212  5.49933  3.67262  3.07583  5.04803  3.26808
3.93692  4.39646  0.71165  3.80890  4.66851  3.48510  4.33336
3.06828  2.24593  3.33932  3.53605  4.04711  5.82231  4.65757      32
k - - -
          2.68618  4.42225  2.77519  2.73123  3.46354  2.40513
3.72494  3.29354  2.67741  2.69355  4.24690  2.90347  2.73739
3.18146  2.89801  2.37887  2.77519  2.98518  4.58477  3.61503
          0.01518  4.59089  5.31324  0.61958  0.77255  0.48576
0.95510
     31   3.78144  5.42839  4.59014  4.59699  5.62003  0.16254
5.60919  5.45342  4.86720  4.98412  6.00454  4.75794  4.83053
5.13592  4.96969  3.98112  4.31173  4.84456  6.46989  5.76432      33
G - - -
          2.68618  4.42225  2.77519  2.73123  3.46354  2.40513
3.72494  3.29354  2.67741  2.69355  4.24690  2.90347  2.73739
3.18146  2.89801  2.37887  2.77519  2.98518  4.58477  3.61503
          0.01518  4.59089  5.31324  0.61958  0.77255  0.48576
0.95510
     32   2.99578  5.35680  3.24876  2.68269  4.73944  3.70201
2.82087  4.16390  1.46687  3.62290  4.41632  3.18328  4.08810
1.69544  2.62343  2.96597  2.88254  3.77791  5.71302  4.40893      34
k - - -
          2.68618  4.42225  2.77519  2.73123  3.46354  2.40513
3.72494  3.29354  2.67741  2.69355  4.24690  2.90347  2.73739
3.18146  2.89801  2.37887  2.77519  2.98518  4.58477  3.61503
          0.01518  4.59089  5.31324  0.61958  0.77255  0.48576
0.95510
     33   3.78144  5.42839  4.59014  4.59699  5.62003  0.16254
5.60919  5.45342  4.86720  4.98412  6.00454  4.75794  4.83053
5.13592  4.96969  3.98112  4.31173  4.84456  6.46989  5.76432      35
G - - -
          2.68618  4.42225  2.77519  2.73123  3.46354  2.40513
3.72494  3.29354  2.67741  2.69355  4.24690  2.90347  2.73739
3.18146  2.89801  2.37887  2.77519  2.98518  4.58477  3.61503

```
          0.06243  4.59089  2.98821  0.61958  0.77255  0.48576
0.95510
     34   2.77793  4.17941  4.52112  3.93012  1.88675  3.36346
4.14732  2.50786  3.75660  2.26185  2.92927  3.99085  4.27574
3.91322  3.83073  3.21829  3.00943  2.29031  2.26780  2.82264     36
f - - -
          2.68618  4.42225  2.77519  2.73123  3.46354  2.40513
3.72494  3.29354  2.67741  2.69355  4.24690  2.90347  2.73739
3.18146  2.89801  2.37887  2.77519  2.98518  4.58477  3.61503
          0.01591  4.54438  5.26672  0.61958  0.77255  0.56747
0.83693
     35   2.61924  3.61700  3.89448  3.31826  3.40995  3.76177
4.04809  2.73833  2.64746  1.99645  3.40291  3.62995  4.13880
3.22484  3.49273  2.91897  2.69194  1.61696  4.89591  3.68855     37
v - - -
          2.68621  4.42228  2.77523  2.73126  3.46357  2.40516
3.72468  3.29357  2.67744  2.69358  4.24630  2.90350  2.73725
3.18149  2.89804  2.37885  2.77523  2.98521  4.58480  3.61506
          0.14692  2.02891  5.26672  0.24876  1.51306  0.56747
0.83693
     36   2.18174  5.14571  2.76785  2.24782  4.45963  3.49806
3.68937  3.92388  1.83963  3.29635  4.19505  2.97875  3.89150
2.62393  2.92292  2.50362  2.61308  3.32315  5.59735  3.77419     39
k - - -
          2.68618  4.42225  2.77519  2.73123  3.46354  2.40513
3.72494  3.29354  2.67741  2.69355  4.24690  2.90347  2.73739
3.18146  2.89801  2.37887  2.77519  2.98518  4.58477  3.61503
          0.01591  4.54438  5.26672  0.61958  0.77255  0.56747
0.83693
     37   2.33406  4.44271  3.09957  3.01085  3.58324  3.68896
3.94368  2.92685  2.71384  2.28589  2.14597  3.42052  4.06959
3.27043  3.32249  2.93371  2.45679  2.35892  5.02785  3.79187     40
m - - -
          2.68618  4.42225  2.77519  2.73123  3.46354  2.40513
3.72494  3.29354  2.67741  2.69355  4.24690  2.90347  2.73739
3.18146  2.89801  2.37887  2.77519  2.98518  4.58477  3.61503
          0.01591  4.54438  5.26672  0.61958  0.77255  0.56747
0.83693
     38   2.67746  4.76255  3.29746  2.84313  4.39293  3.43505
4.02342  3.81173  2.59538  3.42883  4.24382  3.27333  3.31982
3.19916  3.25464  1.35537  1.82748  3.40544  5.66171  4.36255     41
s - - -
          2.68618  4.42225  2.77519  2.73123  3.46354  2.40513
3.72494  3.29354  2.67741  2.69355  4.24690  2.90347  2.73739
3.18146  2.89801  2.37887  2.77519  2.98518  4.58477  3.61503
          0.01591  4.54438  5.26672  0.61958  0.77255  0.56747
0.83693
     39   1.83945  5.09646  3.01764  2.15291  4.39329  3.51665
3.71401  3.40883  1.92762  3.38073  4.15693  3.01108  3.60343
2.74020  2.93519  2.53769  2.96256  3.46201  5.56291  4.19061     42
a - - -
          2.68618  4.42225  2.77519  2.73123  3.46354  2.40513
3.72494  3.29354  2.67741  2.69355  4.24690  2.90347  2.73739
3.18146  2.89801  2.37887  2.77519  2.98518  4.58477  3.61503
```

```
          0.01591  4.54438  5.26672  0.61958  0.77255  0.56747
0.83693
     40   2.08436  5.33759  2.23616  1.85910  4.66803  3.49897
3.75782  4.15422  1.84237  3.63525  4.39152  2.96379  3.94287
2.74812  3.03336  2.77671  3.06581  3.72208  5.77188  4.35058      43
k - - -
          2.68618  4.42225  2.77519  2.73123  3.46354  2.40513
3.72494  3.29354  2.67741  2.69355  4.24690  2.90347  2.73739
3.18146  2.89801  2.37887  2.77519  2.98518  4.58477  3.61503
          0.01591  4.54438  5.26672  0.61958  0.77255  0.43786
1.03680
     41   2.04975  5.25137  2.61120  1.67773  4.58266  3.50807
3.72121  4.06213  2.47412  3.55106  4.29955  2.86304  3.91866
2.54257  2.76329  2.63953  2.74901  3.63536  5.69165  4.28307      44
e - - -
          2.68618  4.42225  2.77519  2.73123  3.46354  2.40513
3.72494  3.29354  2.67741  2.69355  4.24690  2.90347  2.73739
3.18146  2.89801  2.37887  2.77519  2.98518  4.58477  3.61503
          0.04995  4.59089  3.25503  0.61958  0.77255  0.48576
0.95510
     42   3.61615  0.21521  5.18394  5.10602  5.03350  4.04676
5.61883  4.37157  4.97432  4.21327  5.36991  4.94407  4.79177
5.28938  4.95844  3.88288  4.13832  4.04853  6.22276  5.29994      45
C - - -
          2.68618  4.42225  2.77519  2.73123  3.46354  2.40513
3.72494  3.29354  2.67741  2.69355  4.24690  2.90347  2.73739
3.18146  2.89801  2.37887  2.77519  2.98518  4.58477  3.61503
          0.01572  4.55666  5.27901  0.61958  0.77255  0.54689
0.86452
     43   2.30141  5.19425  2.37540  2.11771  4.52138  3.32910
3.69233  3.99550  2.03351  2.98751  4.24107  2.85951  3.89424
2.79571  2.92852  2.51759  2.57067  3.57549  5.63803  4.23670      46
k - - -
          2.68618  4.42225  2.77519  2.73123  3.46354  2.40513
3.72494  3.29354  2.67741  2.69355  4.24690  2.90347  2.73739
3.18146  2.89801  2.37887  2.77519  2.98518  4.58477  3.61503
          0.01572  4.55666  5.27901  0.61958  0.77255  0.54689
0.86452
     44   2.06298  5.21626  2.19815  2.43246  4.54632  3.50176
3.71661  4.01887  1.89629  3.51786  4.27262  2.97921  3.91296
2.53496  2.95524  2.63016  2.61896  3.60113  5.66590  4.26506      47
k - - -
          2.68618  4.42225  2.77519  2.73123  3.46354  2.40513
3.72494  3.29354  2.67741  2.69355  4.24690  2.90347  2.73739
3.18146  2.89801  2.37887  2.77519  2.98518  4.58477  3.61503
          0.01572  4.55666  5.27901  0.61958  0.77255  0.54689
0.86452
     45   2.09352  4.98653  2.79198  2.52157  4.24553  3.53424
3.73239  2.88284  2.03877  3.08494  4.05161  3.04843  3.92503
2.36180  2.84633  2.73462  2.94964  2.91102  5.47434  4.12428      48
k - - -
          2.68618  4.42225  2.77519  2.73123  3.46354  2.40513
3.72494  3.29354  2.67741  2.69355  4.24690  2.90347  2.73739
3.18146  2.89801  2.37887  2.77519  2.98518  4.58477  3.61503
```

```
          0.01572  4.55666  5.27901  0.61958  0.77255  0.54689
0.86452
     46   2.90067  5.30406  2.83489  2.29628  4.61217  1.40681
3.48271  4.06772  2.37777  3.58801  4.37948  2.73599  4.00428
2.75880  3.03847  2.86533  2.94355  3.68004  5.75435  4.36828      49
g - - -
          2.68618  4.42225  2.77519  2.73123  3.46354  2.40513
3.72494  3.29354  2.67741  2.69355  4.24690  2.90347  2.73739
3.18146  2.89801  2.37887  2.77519  2.98518  4.58477  3.61503
          0.01572  4.55666  5.27901  0.61958  0.77255  0.54689
0.86452
     47   2.31933  4.49631  3.85853  3.75834  5.01844  0.63265
4.87648  4.49015  3.98343  4.17429  4.96616  3.76945  4.05836
4.18895  4.23023  2.27926  3.10017  3.74472  6.34414  5.18200      50
G - - -
          2.68618  4.42225  2.77519  2.73123  3.46354  2.40513
3.72494  3.29354  2.67741  2.69355  4.24690  2.90347  2.73739
3.18146  2.89801  2.37887  2.77519  2.98518  4.58477  3.61503
          0.01572  4.55666  5.27901  0.61958  0.77255  0.54689
0.86452
     48   2.54741  5.09656  2.88956  2.47380  4.39523  3.51617
3.46921  3.84477  1.79120  3.38263  4.15602  3.00998  3.91052
2.82795  2.81106  2.36449  2.10810  3.17732  5.56316  4.18883      51
k - - -
          2.68618  4.42225  2.77519  2.73123  3.46354  2.40513
3.72494  3.29354  2.67741  2.69355  4.24690  2.90347  2.73739
3.18146  2.89801  2.37887  2.77519  2.98518  4.58477  3.61503
          0.01572  4.55666  5.27901  0.61958  0.77255  0.54689
0.86452
     49   2.59570  4.24157  4.00540  3.04041  2.57557  3.78561
4.07583  2.20712  2.98804  2.43938  3.11490  3.69799  3.40459
3.57737  3.56551  2.90045  2.69634  1.79194  4.84817  3.64770      52
v - - -
          2.68618  4.42225  2.77519  2.73123  3.46354  2.40513
3.72494  3.29354  2.67741  2.69355  4.24690  2.90347  2.73739
3.18146  2.89801  2.37887  2.77519  2.98518  4.58477  3.61503
          0.03811  4.55666  3.61578  0.61958  0.77255  0.54689
0.86452
     50   2.35615  5.00851  3.05171  2.15809  3.62888  3.19169
3.71558  3.49152  2.07703  2.90132  4.07045  3.02520  3.90939
2.70984  2.61386  2.57099  2.71781  3.10371  5.49021  4.13161      53
k - - -
          2.68618  4.42225  2.77519  2.73123  3.46354  2.40513
3.72494  3.29354  2.67741  2.69355  4.24690  2.90347  2.73739
3.18146  2.89801  2.37887  2.77519  2.98518  4.58477  3.61503
          0.01085  4.52940        *  0.61958  0.77255  0.00000
*
//
```

```
HMMER3/f [3.3.2 | Nov 2020]
NAME  Buf_EGKCG/oxazolin
LENG  77
ALPH  amino
RF    no
MM    no
CONS  yes
CS    no
MAP   yes
DATE  Fri Jun 16 10:47:06 2023
NSEQ  36
EFFN  2.043457
CKSUM 3567737997
STATS LOCAL MSV      -10.1463  0.71860
STATS LOCAL VITERBI  -10.7793  0.71860
STATS LOCAL FORWARD   -3.8743  0.71860
HMM          A        C        D        E        F        G        H
I        K        L        M        N        P        Q        R
S        T        V        W        Y
            m->m     m->i     m->d     i->m     i->i     d->m     d-
>d
  COMPO   2.40970  3.06915  2.91752  2.24985  4.01022  2.18801
3.84238  3.66275  2.23752  3.17422  3.84831  3.07604  3.58382
3.03932  3.11110  2.58563  2.95130  3.23025  5.50827  3.94121
          2.68618  4.42225  2.77519  2.73123  3.46354  2.40513
3.72494  3.29354  2.67741  2.69355  4.24690  2.90347  2.73739
3.18146  2.89801  2.37887  2.77519  2.98518  4.58477  3.61503
          0.03122  4.61826  3.86938  0.61958  0.77255  0.00000
*
      1   2.29031  5.22939  2.31498  1.75182  4.55831  3.29494
3.72227  4.03390  2.47625  3.53011  4.28098  2.98614  3.91899
2.66104  2.97004  2.39778  2.69104  3.29875  5.67642  4.27197      1
e - - -
          2.68618  4.42225  2.77519  2.73123  3.46354  2.40513
3.72494  3.29354  2.67741  2.69355  4.24690  2.90347  2.73739
3.18146  2.89801  2.37887  2.77519  2.98518  4.58477  3.61503
          0.01501  4.60205  5.32440  0.61958  0.77255  0.51625
0.90822
      2   2.84717  5.02994  3.11908  2.67956  4.38747  3.58436
3.89784  3.79890  2.48237  3.39767  3.57346  1.16461  4.04476
3.05432  3.06017  2.71968  2.87744  3.45286  5.61788  4.29139      2
n - - -
          2.68618  4.42225  2.77519  2.73123  3.46354  2.40513
3.72494  3.29354  2.67741  2.69355  4.24690  2.90347  2.73739
3.18146  2.89801  2.37887  2.77519  2.98518  4.58477  3.61503
          0.01468  4.62419  5.34653  0.61958  0.77255  0.51625
0.90822
      3   2.50915  4.81904  2.86948  2.98590  4.70606  3.42034
4.30706  4.13167  3.24714  3.76730  4.58821  3.38819  0.91232
3.51120  3.67855  2.64617  2.80023  3.64605  5.99577  4.69507      3
p - - -
          2.68618  4.42225  2.77519  2.73123  3.46354  2.40513
3.72494  3.29354  2.67741  2.69355  4.24690  2.90347  2.73739
3.18146  2.89801  2.37887  2.77519  2.98518  4.58477  3.61503
```

```
          0.01468  4.62419  5.34653  0.61958  0.77255  0.51625
0.90822
      4   2.95101  4.73280  2.98080  3.03796  1.03576  3.23268
4.07979  3.30124  2.90655  2.98956  3.91159  3.46802  4.22190
3.47307  3.55473  3.11977  3.21973  3.06355  5.10717  3.70021      4
f - - -
          2.68618  4.42225  2.77519  2.73123  3.46354  2.40513
3.72494  3.29354  2.67741  2.69355  4.24690  2.90347  2.73739
3.18146  2.89801  2.37887  2.77519  2.98518  4.58477  3.61503
          0.01468  4.62419  5.34653  0.61958  0.77255  0.51625
0.90822
      5   1.45293  5.01568  2.92200  2.47779  4.35056  2.64709
3.82188  3.78336  2.62199  3.36296  4.16180  3.09718  3.97115
2.77763  3.09491  2.43784  2.90130  3.26150  5.58110  4.22855      5
a - - -
          2.68618  4.42225  2.77519  2.73123  3.46354  2.40513
3.72494  3.29354  2.67741  2.69355  4.24690  2.90347  2.73739
3.18146  2.89801  2.37887  2.77519  2.98518  4.58477  3.61503
          0.01468  4.62419  5.34653  0.61958  0.77255  0.46716
0.98558
      6   1.80238  3.62729  4.60860  4.01340  3.32950  4.01516
4.35254  2.12645  3.84920  1.73774  2.26250  4.10655  4.37150
4.01912  3.93654  3.32264  2.98473  2.22148  4.90199  3.72522      6
l - - -
          2.68618  4.42225  2.77519  2.73123  3.46354  2.40513
3.72494  3.29354  2.67741  2.69355  4.24690  2.90347  2.73739
3.18146  2.89801  2.37887  2.77519  2.98518  4.58477  3.61503
          0.01445  4.64004  5.36239  0.61958  0.77255  0.48576
0.95510
      7   2.58835  5.10871  3.05997  2.39349  4.39904  3.54459
3.73691  3.84604  2.02839  3.38959  3.69070  3.03883  3.93652
2.45769  2.88535  2.25761  2.04231  3.16791  5.57561  4.20350      7
k - - -
          2.68618  4.42225  2.77519  2.73123  3.46354  2.40513
3.72494  3.29354  2.67741  2.69355  4.24690  2.90347  2.73739
3.18146  2.89801  2.37887  2.77519  2.98518  4.58477  3.61503
          0.01445  4.64004  5.36239  0.61958  0.77255  0.48576
0.95510
      8   2.76697  5.25938  2.15833  1.76043  4.59367  3.30841
3.72653  4.07462  2.35682  3.56045  4.30522  2.99170  3.23326
2.75222  2.96948  2.40386  2.66481  3.26353  5.69817  4.28916      8
e - - -
          2.68618  4.42225  2.77519  2.73123  3.46354  2.40513
3.72494  3.29354  2.67741  2.69355  4.24690  2.90347  2.73739
3.18146  2.89801  2.37887  2.77519  2.98518  4.58477  3.61503
          0.01445  4.64004  5.36239  0.61958  0.77255  0.48576
0.95510
      9   2.85726  3.67765  4.56681  3.96918  3.27055  4.00450
3.91240  2.58706  3.80063  1.03509  2.86716  4.07836  4.35578
3.97216  3.89485  3.30816  3.08842  2.19040  4.86313  3.45952      9
l - - -
          2.68618  4.42225  2.77519  2.73123  3.46354  2.40513
3.72494  3.29354  2.67741  2.69355  4.24690  2.90347  2.73739
3.18146  2.89801  2.37887  2.77519  2.98518  4.58477  3.61503
```

```
          0.01445  4.64004  5.36239  0.61958  0.77255  0.48576
0.95510
     10   2.76217  5.20915  2.64118  2.40996  4.53030  2.75875
3.73355  3.99920  2.37711  3.50654  4.26260  2.65107  3.93094
2.76860  2.97744  1.81276  2.34460  3.31048  5.66180  4.26476      10
s - - -
          2.68618  4.42225  2.77519  2.73123  3.46354  2.40513
3.72494  3.29354  2.67741  2.69355  4.24690  2.90347  2.73739
3.18146  2.89801  2.37887  2.77519  2.98518  4.58477  3.61503
          0.01445  4.64004  5.36239  0.61958  0.77255  0.48576
0.95510
     11   2.42915  5.15038  2.82296  2.49614  4.47496  2.71485
3.38646  3.93166  2.52824  3.46342  4.23358  2.89442  3.94507
2.74809  3.01391  1.64749  2.52054  3.53786  5.63915  4.25584      11
s - - -
          2.68618  4.42225  2.77519  2.73123  3.46354  2.40513
3.72494  3.29354  2.67741  2.69355  4.24690  2.90347  2.73739
3.18146  2.89801  2.37887  2.77519  2.98518  4.58477  3.61503
          0.01445  4.64004  5.36239  0.61958  0.77255  0.48576
0.95510
     12   2.54416  4.94186  2.76035  3.09320  5.09307  0.61986
4.58693  4.56550  3.67444  4.20786  5.04688  3.48771  4.20016
3.82666  4.11819  3.00486  3.37593  3.96645  6.37024  5.07198      12
G - - -
          2.68618  4.42225  2.77519  2.73123  3.46354  2.40513
3.72494  3.29354  2.67741  2.69355  4.24690  2.90347  2.73739
3.18146  2.89801  2.37887  2.77519  2.98518  4.58477  3.61503
          0.01445  4.64004  5.36239  0.61958  0.77255  0.48576
0.95510
     13   3.15119  4.92011  3.69909  3.28483  3.39503  3.92250
4.03157  3.50801  2.48488  3.13879  4.10433  3.64605  4.37168
3.49318  3.20828  3.29669  2.97736  3.27793  4.92295  0.91372      13
y - - -
          2.68605  4.42231  2.77526  2.73116  3.46360  2.40519
3.72501  3.29360  2.67747  2.69361  4.24528  2.90353  2.73746
3.18152  2.89807  2.37889  2.77519  2.98506  4.58483  3.61509
          0.33466  1.48458  2.85036  0.15763  1.92530  0.48576
0.95510
     14   2.80383  5.12242  3.03158  2.39467  4.40857  3.22231
3.76806  3.84400  2.44187  3.39604  2.83886  2.76762  3.97141
1.48532  2.94650  2.80020  3.03462  3.48326  5.58743  4.23167      15
q - - -
          2.68618  4.42225  2.77519  2.73123  3.46354  2.40513
3.72494  3.29354  2.67741  2.69355  4.24690  2.90347  2.73739
3.18146  2.89801  2.37887  2.77519  2.98518  4.58477  3.61503
          0.01527  4.58545  5.30779  0.61958  0.77255  0.58519
0.81419
     15   3.46340  4.71881  5.40844  4.86142  3.60649  4.87354
5.31780  1.83281  4.73994  1.43067  2.77434  5.00144  5.08523
4.84274  4.80889  4.24352  3.69939  1.03585  5.61589  4.51189      16
v - - -
          2.68618  4.42225  2.77519  2.73123  3.46354  2.40513
3.72494  3.29354  2.67741  2.69355  4.24690  2.90347  2.73739
3.18146  2.89801  2.37887  2.77519  2.98518  4.58477  3.61503
```

```
          0.01527  4.58545  5.30779  0.61958  0.77255  0.58519
0.81419
     16   0.78148  4.51000  4.34248  3.91260  4.08633  3.66460
4.73131  2.60382  3.81620  2.90544  3.99557  4.01405  4.31585
4.10622  4.06297  3.08620  3.20307  2.18893  5.70223  4.49559      17
a - - -
          2.68618  4.42225  2.77519  2.73123  3.46354  2.40513
3.72494  3.29354  2.67741  2.69355  4.24690  2.90347  2.73739
3.18146  2.89801  2.37887  2.77519  2.98518  4.58477  3.61503
          0.06272  4.58545  2.98383  0.61958  0.77255  0.58519
0.81419
     17   2.64355  5.21792  2.16917  1.93374  4.54818  2.80962
3.51416  4.02678  2.44373  3.21273  4.26405  2.96014  3.89387
2.31587  2.93751  2.55291  2.96648  3.60091  5.65800  3.86117      18
e - - -
          2.68618  4.42225  2.77519  2.73123  3.46354  2.40513
3.72494  3.29354  2.67741  2.69355  4.24690  2.90347  2.73739
3.18146  2.89801  2.37887  2.77519  2.98518  4.58477  3.61503
          0.06385  4.53872  2.97261  0.61958  0.77255  0.65935
0.72812
     18   2.01835  4.75489  3.18938  2.63028  3.95796  3.02789
3.25211  3.20013  2.55073  2.99765  3.45719  2.63668  3.94564
2.95812  3.06020  2.34597  2.92448  2.85153  5.28427  3.30122      19
a - - -
          2.68618  4.42225  2.77519  2.73123  3.46354  2.40513
3.72494  3.29354  2.67741  2.69355  4.24690  2.90347  2.73739
3.18146  2.89801  2.37887  2.77519  2.98518  4.58477  3.61503
          0.16906  4.49165  1.93559  0.61958  0.77255  0.56888
0.83507
     19   2.47114  3.59910  2.61335  2.36762  4.16657  2.89156
2.35916  3.59001  2.49294  3.18111  3.98563  3.01569  3.88533
2.84327  2.96129  2.69653  2.71732  2.98197  5.41251  4.06820      20
h - - -
          2.68579  4.42225  2.77471  2.73080  3.46389  2.40508
3.72493  3.29390  2.67699  2.69390  4.24685  2.90375  2.73731
3.18172  2.89836  2.37908  2.77524  2.98505  4.58512  3.61539
          0.63644  0.90100  2.73851  1.29418  0.32038  0.53633
0.87922
     20   2.55451  5.17352  2.24306  2.17906  4.50094  3.05518
3.67112  3.97497  1.88729  3.47162  3.71994  2.94734  3.87248
2.77489  2.90752  2.54236  2.62831  3.55508  5.61766  4.21627      35
k - - -
          2.68618  4.42225  2.77519  2.73123  3.46354  2.40513
3.72494  3.29354  2.67741  2.69355  4.24690  2.90347  2.73739
3.18146  2.89801  2.37887  2.77519  2.98518  4.58477  3.61503
          0.05635  4.48097  3.13570  0.61958  0.77255  0.65040
0.73781
     21   2.83596  5.23229  3.09753  2.30470  4.58537  3.57790
3.70938  4.02540  1.29522  3.50827  4.28500  3.05740  3.68173
2.60919  2.52120  2.68163  3.05321  3.63060  5.62956  3.85807      36
k - - -
          2.68618  4.42225  2.77519  2.73123  3.46354  2.40513
3.72494  3.29354  2.67741  2.69355  4.24690  2.90347  2.73739
3.18146  2.89801  2.37887  2.77519  2.98518  4.58477  3.61503
```

```
          0.01703  4.47732  5.19966  0.61958  0.77255  0.61565
0.77715
     22   2.37288  4.47897  2.56204  2.48601  4.29089  2.77706
3.71082  3.72824  2.22528  3.29255  3.40712  3.01778  3.90447
2.68460  2.95952  2.44558  2.38902  3.10792  5.49826  4.13654      37
k - - -
          2.68618  4.42225  2.77519  2.73123  3.46354  2.40513
3.72494  3.29354  2.67741  2.69355  4.24690  2.90347  2.73739
3.18146  2.89801  2.37887  2.77519  2.98518  4.58477  3.61503
          0.03514  4.52673  3.74164  0.61958  0.77255  0.55712
0.85063
     23   2.08371  5.15200  2.80414  2.36206  4.46812  2.79272
3.69424  3.93304  1.93559  3.44575  3.98159  2.98119  3.59950
2.72306  2.92926  2.56575  2.51311  3.52757  5.60501  4.21328      38
k - - -
          2.68618  4.42225  2.77519  2.73123  3.46354  2.40513
3.72494  3.29354  2.67741  2.69355  4.24690  2.90347  2.73739
3.18146  2.89801  2.37887  2.77519  2.98518  4.58477  3.61503
          0.01578  4.55254  5.27489  0.61958  0.77255  0.44343
1.02675
     24   3.33747  5.99972  1.85796  0.77976  5.29533  3.55883
4.13160  4.84643  3.18759  4.30735  5.15940  3.00965  4.20002
3.29729  3.82058  2.83872  3.63363  4.38436  6.45394  4.90191      39
e - - -
          2.68618  4.42225  2.77519  2.73123  3.46354  2.40513
3.72494  3.29354  2.67741  2.69355  4.24690  2.90347  2.73739
3.18146  2.89801  2.37887  2.77519  2.98518  4.58477  3.61503
          0.01469  4.62342  5.34577  0.61958  0.77255  0.51768
0.90609
     25   2.24889  4.53352  3.89761  3.54946  4.52115  0.80055
4.56130  3.89594  3.55108  3.60927  3.37378  3.70544  4.09170
3.84843  3.85813  2.52649  3.08514  3.42399  5.89490  4.68476      40
g - - -
          2.68618  4.42225  2.77519  2.73123  3.46354  2.40513
3.72494  3.29354  2.67741  2.69355  4.24690  2.90347  2.73739
3.18146  2.89801  2.37887  2.77519  2.98518  4.58477  3.61503
          0.01469  4.62342  5.34577  0.61958  0.77255  0.51768
0.90609
     26   2.84642  5.11987  3.21798  2.63470  4.42578  3.63207
3.77979  3.84167  1.29049  3.39070  3.65717  3.14410  4.01749
2.90896  2.36752  2.54167  2.85683  3.14122  5.57116  4.24835      41
k - - -
          2.68618  4.42225  2.77519  2.73123  3.46354  2.40513
3.72494  3.29354  2.67741  2.69355  4.24690  2.90347  2.73739
3.18146  2.89801  2.37887  2.77519  2.98518  4.58477  3.61503
          0.01469  4.62342  5.34577  0.61958  0.77255  0.51768
0.90609
     27   2.77278  1.06226  4.28748  3.76471  3.51303  3.73689
4.32325  2.88553  3.58446  2.66415  3.19328  3.90224  4.25336
3.86835  3.77407  2.78924  3.09056  2.67158  5.05008  3.41122      42
c - - -
          2.68624  4.42246  2.77528  2.73080  3.46375  2.40476
3.72516  3.29375  2.67719  2.69376  4.24631  2.90349  2.73761
3.18142  2.89783  2.37888  2.77541  2.98540  4.58498  3.61524
```

0.15417  1.97975  5.34577  1.55204  0.23802  0.46631
0.98702
28  2.46843  4.88358  3.16376  2.48485  4.72877  0.99994
4.24744  4.17154  3.15663  3.77843  4.58972  3.34440  4.08732
3.43511  3.59976  2.30445  2.96159  3.68596  5.99207  4.67586  54
g - - -
2.68618  4.42225  2.77519  2.73123  3.46354  2.40513
3.72494  3.29354  2.67741  2.69355  4.24690  2.90347  2.73739
3.18146  2.89801  2.37887  2.77519  2.98518  4.58477  3.61503
0.01445  4.64004  5.36239  0.61958  0.77255  0.48576
0.95510
29  2.65683  5.64532  2.29493  0.86836  4.98910  3.55522
4.01677  4.47738  2.94349  3.98325  4.79303  3.03183  4.12269
3.16251  3.50025  3.04354  3.03686  4.04616  6.14560  4.67612  55
e - - -
2.68618  4.42225  2.77519  2.73123  3.46354  2.40513
3.72494  3.29354  2.67741  2.69355  4.24690  2.90347  2.73739
3.18146  2.89801  2.37887  2.77519  2.98518  4.58477  3.61503
0.01445  4.64004  5.36239  0.61958  0.77255  0.48576
0.95510
30  1.93026  4.61871  4.09763  4.01628  5.14285  0.55428
5.07529  4.58637  4.21539  4.30563  5.12490  3.96754  4.19744
4.41972  4.42096  2.89405  3.25339  3.86281  6.44325  5.33625  56
G - - -
2.68618  4.42225  2.77519  2.73123  3.46354  2.40513
3.72494  3.29354  2.67741  2.69355  4.24690  2.90347  2.73739
3.18146  2.89801  2.37887  2.77519  2.98518  4.58477  3.61503
0.01445  4.64004  5.36239  0.61958  0.77255  0.48576
0.95510
31  2.98819  5.43036  3.61261  2.99821  4.94564  3.88285
3.91527  4.30347  0.79260  3.74496  4.58461  3.42794  4.27842
3.04252  2.26341  2.84718  3.43644  3.94484  5.79748  4.59496  57
k - - -
2.68618  4.42225  2.77519  2.73123  3.46354  2.40513
3.72494  3.29354  2.67741  2.69355  4.24690  2.90347  2.73739
3.18146  2.89801  2.37887  2.77519  2.98518  4.58477  3.61503
0.01445  4.64004  5.36239  0.61958  0.77255  0.48576
0.95510
32  2.74562  0.47398  4.78753  4.63801  4.69850  3.45785
5.20238  4.02652  4.43610  3.92088  4.86887  4.24251  4.28197
4.70617  4.49880  2.68277  3.29733  3.54004  6.14077  4.94758  58
C - - -
2.68618  4.42225  2.77519  2.73123  3.46354  2.40513
3.72494  3.29354  2.67741  2.69355  4.24690  2.90347  2.73739
3.18146  2.89801  2.37887  2.77519  2.98518  4.58477  3.61503
0.01445  4.64004  5.36239  0.61958  0.77255  0.48576
0.95510
33  2.44246  4.57488  3.92036  3.86211  5.13529  0.52764
4.99529  4.61970  4.12419  4.30991  5.10389  3.85826  4.13949
4.32037  4.35278  2.50753  3.18936  3.84960  6.46067  5.30337  59
G - - -
2.68618  4.42225  2.77519  2.73123  3.46354  2.40513
3.72494  3.29354  2.67741  2.69355  4.24690  2.90347  2.73739
3.18146  2.89801  2.37887  2.77519  2.98518  4.58477  3.61503

```
          0.03488  4.64004  3.70400  0.61958  0.77255  0.48576
0.95510
     34   1.61343  5.43207  2.37412  1.94313  4.76910  2.22182
3.88094  4.25549  2.72458  3.75530  4.53395  2.64904  4.02729
3.00784  3.24952  2.89177  3.20208  3.83042  5.91181  4.47836      60
a - - -
          2.68618  4.42225  2.77519  2.73123  3.46354  2.40513
3.72494  3.29354  2.67741  2.69355  4.24690  2.90347  2.73739
3.18146  2.89801  2.37887  2.77519  2.98518  4.58477  3.61503
          0.01475  4.61991  5.34225  0.61958  0.77255  0.52423
0.89651
     35   2.76796  5.27967  2.49952  1.98667  4.62019  2.82932
3.72155  4.10673  2.05475  3.58295  4.32365  2.66759  3.92131
2.55928  2.96596  2.23653  2.70661  3.66827  5.71319  4.29858      61
e - - -
          2.68618  4.42225  2.77519  2.73123  3.46354  2.40513
3.72494  3.29354  2.67741  2.69355  4.24690  2.90347  2.73739
3.18146  2.89801  2.37887  2.77519  2.98518  4.58477  3.61503
          0.01475  4.61991  5.34225  0.61958  0.77255  0.52423
0.89651
     36   2.56927  5.03303  3.10610  2.37327  4.30727  3.39670
3.76538  3.73440  1.59885  3.30879  2.56949  3.08216  3.96076
2.90020  2.97628  2.41432  2.87964  3.38704  5.52180  4.17458      62
k - - -
          2.68580  4.42185  2.77522  2.73066  3.46382  2.40499
3.72523  3.29382  2.67712  2.69383  4.24575  2.90375  2.73747
3.18174  2.89801  2.37893  2.77533  2.98541  4.58505  3.61531
          0.79862  1.11352  1.50666  1.08160  0.41408  0.46244
0.99357
     37   2.35900  5.01813  2.88215  2.10320  4.30274  3.47059
3.66685  3.74587  1.87382  3.29715  3.24654  2.96818  3.46741
2.78848  2.90661  2.66886  2.74631  3.10265  5.49010  4.12078      73
k - - -
          2.68618  4.42225  2.77519  2.73123  3.46354  2.40513
3.72494  3.29354  2.67741  2.69355  4.24690  2.90347  2.73739
3.18146  2.89801  2.37887  2.77519  2.98518  4.58477  3.61503
          0.01839  4.40110  5.12345  0.61958  0.77255  0.52287
0.89849
     38   2.48237  5.22903  3.20645  2.63275  4.59066  3.63904
3.21514  4.01389  1.23070  3.50962  4.30315  3.13700  4.02784
2.87672  2.46470  2.54509  3.11529  3.63807  5.63592  4.32185      74
k - - -
          2.68618  4.42225  2.77519  2.73123  3.46354  2.40513
3.72494  3.29354  2.67741  2.69355  4.24690  2.90347  2.73739
3.18146  2.89801  2.37887  2.77519  2.98518  4.58477  3.61503
          0.01603  4.53698  5.25932  0.61958  0.77255  0.53648
0.87900
     39   2.11354  4.44300  2.35806  2.14974  4.51030  3.07151
3.69662  3.98193  2.12138  3.48255  3.57028  2.62570  3.89949
2.80040  2.93028  2.63610  2.95750  3.56552  5.63164  4.23329      75
a - - -
          2.68618  4.42225  2.77519  2.73123  3.46354  2.40513
3.72494  3.29354  2.67741  2.69355  4.24690  2.90347  2.73739
3.18146  2.89801  2.37887  2.77519  2.98518  4.58477  3.61503
```

0.01532 4.58228 5.30463 0.61958 0.77255 0.49825
0.93545
40 2.21913 5.15768 2.83278 2.46241 4.46865 2.58877
3.49785 3.93070 2.02500 3.44862 3.67124 3.00411 3.21508
2.68005 2.94901 2.62642 2.46340 3.53010 5.61235 4.22359 76
k - - -
2.68618 4.42225 2.77519 2.73123 3.46354 2.40513
3.72494 3.29354 2.67741 2.69355 4.24690 2.90347 2.73739
3.18146 2.89801 2.37887 2.77519 2.98518 4.58477 3.61503
0.01484 4.61380 5.33614 0.61958 0.77255 0.53547
0.88042
41 3.20742 5.71910 2.48056 0.75796 5.03939 3.59174
4.03534 4.54281 2.90093 4.02662 4.85462 3.06211 4.16263
3.18616 2.90647 3.11540 3.47551 4.12191 6.15745 4.71089 77
e - - -
2.68618 4.42225 2.77519 2.73123 3.46354 2.40513
3.72494 3.29354 2.67741 2.69355 4.24690 2.90347 2.73739
3.18146 2.89801 2.37887 2.77519 2.98518 4.58477 3.61503
0.01484 4.61380 5.33614 0.61958 0.77255 0.53547
0.88042
42 2.42390 4.55163 3.90042 3.55604 4.31384 0.84243
4.53123 3.59871 3.52464 3.34032 2.66936 3.73721 4.14438
3.84631 3.81214 2.88812 3.12713 3.24006 5.75160 4.53400 78
g - - -
2.68618 4.42225 2.77519 2.73123 3.46354 2.40513
3.72494 3.29354 2.67741 2.69355 4.24690 2.90347 2.73739
3.18146 2.89801 2.37887 2.77519 2.98518 4.58477 3.61503
0.01484 4.61380 5.33614 0.61958 0.77255 0.45589
1.00481
43 2.58075 5.27170 3.19904 2.65146 4.66383 3.65160
3.80856 4.09585 1.17484 3.58622 4.37189 2.87549 4.05611
2.92487 2.53336 2.36224 3.14711 3.70326 5.70987 4.38271 79
k - - -
2.68618 4.42225 2.77519 2.73123 3.46354 2.40513
3.72494 3.29354 2.67741 2.69355 4.24690 2.90347 2.73739
3.18146 2.89801 2.37887 2.77519 2.98518 4.58477 3.61503
0.01445 4.64004 5.36239 0.61958 0.77255 0.48576
0.95510
44 2.33556 0.82946 3.11919 3.59716 4.36379 3.42507
4.58382 3.68288 3.66379 3.45181 4.32721 3.75321 3.51071
3.92024 3.95430 2.86990 3.11585 3.28437 5.78181 4.57789 80
c - - -
2.68618 4.42225 2.77519 2.73123 3.46354 2.40513
3.72494 3.29354 2.67741 2.69355 4.24690 2.90347 2.73739
3.18146 2.89801 2.37887 2.77519 2.98518 4.58477 3.61503
0.01445 4.64004 5.36239 0.61958 0.77255 0.48576
0.95510
45 2.51698 4.92052 3.12510 2.91107 4.70255 0.89320
3.47612 4.16818 3.15632 3.77374 4.59873 2.91355 4.11415
3.45227 3.58131 2.70820 3.22636 3.70141 5.97736 4.65661 81
g - - -
2.68618 4.42225 2.77519 2.73123 3.46354 2.40513
3.72494 3.29354 2.67741 2.69355 4.24690 2.90347 2.73739
3.18146 2.89801 2.37887 2.77519 2.98518 4.58477 3.61503

```
          0.03493  4.64004  3.70229  0.61958  0.77255  0.48576
0.95510
     46   2.16244  5.41850  2.52300  1.27610  4.74459  3.07468
3.84454  4.23202  2.66483  3.72383  4.49365  2.99947  4.00689
2.69734  3.18295  2.66697  3.16401  3.80482  5.87278  4.44163     82
e - - -
          2.68618  4.42225  2.77519  2.73123  3.46354  2.40513
3.72494  3.29354  2.67741  2.69355  4.24690  2.90347  2.73739
3.18146  2.89801  2.37887  2.77519  2.98518  4.58477  3.61503
          0.07508  4.61986  2.77289  0.61958  0.77255  0.52431
0.89639
     47   2.21949  5.00299  2.73918  2.53743  4.31741  1.79471
3.77478  3.75017  2.36366  3.32619  3.55013  3.06121  3.93926
2.91167  3.03579  2.47768  2.98023  3.39093  5.54095  4.18654     83
g - - -
          2.68618  4.42225  2.77519  2.73123  3.46354  2.40513
3.72494  3.29354  2.67741  2.69355  4.24690  2.90347  2.73739
3.18146  2.89801  2.37887  2.77519  2.98518  4.58477  3.61503
          0.01566  4.56044  5.28278  0.61958  0.77255  0.62600
0.76513
     48   2.41832  4.97719  3.18729  2.62180  4.26329  3.58314
3.77740  3.67127  1.37343  3.26000  3.19982  3.12776  3.98027
2.92288  2.80856  2.62317  2.76180  3.34014  5.48128  4.16171     84
k - - -
          2.68611  4.42229  2.77523  2.73115  3.46357  2.40508
3.72498  3.29358  2.67733  2.69358  4.24666  2.90350  2.73743
3.18150  2.89804  2.37886  2.77523  2.98522  4.58481  3.61507
          0.25795  3.25333  1.66747  1.06205  0.42426  0.62600
0.76513
     49   2.43157  1.73019  3.33674  2.59235  3.86440  3.50620
3.89769  3.22546  2.42024  2.91596  3.78645  3.26949  3.97362
3.14333  3.12126  2.59969  2.93962  2.94767  5.23581  3.98060     91
c - - -
          2.68618  4.42225  2.77519  2.73123  3.46354  2.40513
3.72494  3.29354  2.67741  2.69355  4.24690  2.90347  2.73739
3.18146  2.89801  2.37887  2.77519  2.98518  4.58477  3.61503
          0.01921  4.35753  5.07988  0.61958  0.77255  0.83614
0.56806
     50   2.49328  4.61018  3.26621  2.75686  3.95310  1.69439
3.86754  3.33856  2.73621  3.00219  2.96464  3.20458  3.93605
2.78532  3.14539  2.56637  2.69075  3.03593  5.30246  4.02413     92
g - - -
          2.68618  4.42225  2.77519  2.73123  3.46354  2.40513
3.72494  3.29354  2.67741  2.69355  4.24690  2.90347  2.73739
3.18146  2.89801  2.37887  2.77519  2.98518  4.58477  3.61503
          0.01885  4.37623  5.09857  0.61958  0.77255  0.86160
0.54902
     51   1.85044  5.19357  2.47066  1.78175  4.53363  3.07118
3.73418  4.00416  2.54340  3.52195  4.29721  2.92503  3.29630
2.85737  3.04875  2.60172  3.01460  3.59415  5.68910  4.28150     93
e - - -
          2.68611  4.42230  2.77496  2.73110  3.46359  2.40514
3.72500  3.29359  2.67746  2.69360  4.24657  2.90352  2.73745
3.18152  2.89806  2.37892  2.77512  2.98524  4.58482  3.61509
```

```
          0.28028  1.79265  2.55219  0.54036  0.87356  0.86160
0.54902
     52   2.41645  5.08375  2.51230  2.20236  4.39903  2.86768
3.63280  3.86266  2.05196  3.37920  4.14056  2.90879  3.82880
2.74336  2.87716  2.38441  2.71279  3.15125  5.54212  4.15032      97
k - - -
          2.68618  4.42225  2.77519  2.73123  3.46354  2.40513
3.72494  3.29354  2.67741  2.69355  4.24690  2.90347  2.73739
3.18146  2.89801  2.37887  2.77519  2.98518  4.58477  3.61503
          0.02033  4.30171  5.02406  0.61958  0.77255  0.70729
0.67920
     53   2.56123  5.00709  3.10288  2.67689  4.58318  2.63148
3.86320  4.00408  1.16430  3.54450  4.35340  3.15053  3.98932
3.00593  2.84892  2.66395  3.09256  3.59219  5.69697  4.39102      98
k - - -
          2.68618  4.42225  2.77519  2.73123  3.46354  2.40513
3.72494  3.29354  2.67741  2.69355  4.24690  2.90347  2.73739
3.18146  2.89801  2.37887  2.77519  2.98518  4.58477  3.61503
          0.01827  4.40726  5.12961  0.61958  0.77255  0.74713
0.64193
     54   2.24238  3.82478  2.88968  2.49435  4.18938  2.79695
3.70287  3.61559  2.23567  3.20152  3.53042  2.88644  3.21813
2.84192  2.95944  2.36724  2.65366  3.27486  5.42789  4.08074      99
k - - -
          2.68619  4.42230  2.77525  2.73129  3.46359  2.40509
3.72500  3.29359  2.67738  2.69360  4.24533  2.90352  2.73745
3.18152  2.89806  2.37875  2.77525  2.98524  4.58482  3.61508
          0.37593  2.88141  1.35755  1.16713  0.37289  0.67860
0.70791
     55   2.27881  5.04483  2.68492  2.04704  4.35009  2.90636
3.38207  3.80927  2.15211  3.33311  3.72628  2.88964  3.80715
2.72289  2.85233  2.49274  2.86634  3.41448  5.50244  4.11532     106
e - - -
          2.68618  4.42225  2.77519  2.73123  3.46354  2.40513
3.72494  3.29354  2.67741  2.69355  4.24690  2.90347  2.73739
3.18146  2.89801  2.37887  2.77519  2.98518  4.58477  3.61503
          0.02232  4.20910  4.93145  0.61958  0.77255  0.96494
0.47966
     56   3.01360  5.56193  2.07000  1.06525  4.88355  2.74332
3.86224  4.39811  2.84759  3.89278  4.72682  2.81260  3.94731
3.02156  3.42834  2.91019  3.29916  3.96423  6.04746  4.55702     107
e - - -
          2.68618  4.42225  2.77519  2.73123  3.46354  2.40513
3.72494  3.29354  2.67741  2.69355  4.24690  2.90347  2.73739
3.18146  2.89801  2.37887  2.77519  2.98518  4.58477  3.61503
          0.02184  4.23093  4.95327  0.61958  0.77255  0.99293
0.46282
     57   2.31634  4.72645  3.09706  2.69862  4.29326  1.49611
3.86728  3.69744  2.46775  3.30767  4.14274  3.13933  3.90599
3.04638  2.55619  2.73105  2.97238  3.31922  5.52723  4.24369     108
g - - -
          2.68618  4.42225  2.77519  2.73123  3.46354  2.40513
3.72494  3.29354  2.67741  2.69355  4.24690  2.90347  2.73739
3.18146  2.89801  2.37887  2.77519  2.98518  4.58477  3.61503
```

```
          0.02184  4.23093  4.95327  0.61958  0.77255  0.99293
0.46282
     58   2.61373  5.06127  2.99235  2.51085  4.44278  3.48236
3.68998  3.86589  1.53073  3.39018  4.19512  3.01678  3.91264
2.52748  2.67096  2.12647  3.01137  3.49336  5.54213  4.22122    109
k - - -
          2.68618  4.42225  2.77519  2.73123  3.46354  2.40513
3.72494  3.29354  2.67741  2.69355  4.24690  2.90347  2.73739
3.18146  2.89801  2.37887  2.77519  2.98518  4.58477  3.61503
          0.04415  4.23093  3.55262  0.61958  0.77255  0.99293
0.46282
     59   2.51980  1.42499  3.69790  3.25891  3.63788  3.34622
3.09436  3.15840  3.09602  2.87752  3.78153  3.48548  3.94683
3.47331  3.36043  2.46342  2.89198  2.85803  5.10168  3.79745    110
c - - -
          2.68618  4.42225  2.77519  2.73123  3.46354  2.40513
3.72494  3.29354  2.67741  2.69355  4.24690  2.90347  2.73739
3.18146  2.89801  2.37887  2.77519  2.98518  4.58477  3.61503
          0.02232  4.20910  4.93145  0.61958  0.77255  0.84939
0.55805
     60   2.35005  4.61695  3.04824  2.61646  4.45432  1.29056
4.06726  3.88823  2.97699  3.51112  4.32972  3.18451  3.39842
3.26666  3.40037  2.20782  2.95899  3.41747  5.73866  4.44185    111
g - - -
          2.68566  4.42124  2.77496  2.73051  3.46388  2.40476
3.72529  3.29388  2.67728  2.69389  4.24637  2.90361  2.73774
3.18180  2.89827  2.37889  2.77537  2.98552  4.58511  3.61537
          0.74073  0.66053  5.01183  1.15252  0.37956  0.65129
0.73683
     61   2.73907  5.23797  2.54398  2.04245  4.57247  3.12971
3.47401  4.05449  1.63304  3.53640  4.28628  2.92657  3.87919
2.62034  2.92349  2.53485  2.86651  3.62389  5.67139  4.25997    120
k - - -
          2.68618  4.42225  2.77519  2.73123  3.46354  2.40513
3.72494  3.29354  2.67741  2.69355  4.24690  2.90347  2.73739
3.18146  2.89801  2.37887  2.77519  2.98518  4.58477  3.61503
          0.01785  4.43026  5.15261  0.61958  0.77255  0.36965
1.17433
     62   2.38610  5.15915  2.87913  2.50411  4.52806  3.31355
3.76707  3.98966  1.38661  3.50826  4.27275  3.03881  3.45591
2.88097  2.99301  2.35588  2.88007  3.57946  5.66994  4.28599    121
k - - -
          2.68618  4.42225  2.77519  2.73123  3.46354  2.40513
3.72494  3.29354  2.67741  2.69355  4.24690  2.90347  2.73739
3.18146  2.89801  2.37887  2.77519  2.98518  4.58477  3.61503
          0.01475  4.61986  5.34221  0.61958  0.77255  0.52431
0.89639
     63   2.23062  5.18742  2.19039  2.33715  4.50739  2.88308
3.71106  3.97611  2.19970  3.48201  3.49951  2.99686  3.91346
2.49615  2.94555  2.53962  2.96979  3.13720  5.63555  4.23997    122
d - - -
          2.68618  4.42225  2.77519  2.73123  3.46354  2.40513
3.72494  3.29354  2.67741  2.69355  4.24690  2.90347  2.73739
3.18146  2.89801  2.37887  2.77519  2.98518  4.58477  3.61503
```

```
          0.01475  4.61986  5.34221  0.61958  0.77255  0.52431
0.89639
     64   2.06085  5.21011  2.98381  2.12355  4.53740  2.36434
3.73024  4.00614  1.98503  3.51046  4.26655  3.00530  3.61834
2.68262  2.96020  2.64076  2.88340  3.59350  5.66243  4.26700    123
k - - -
          2.68618  4.42225  2.77519  2.73123  3.46354  2.40513
3.72494  3.29354  2.67741  2.69355  4.24690  2.90347  2.73739
3.18146  2.89801  2.37887  2.77519  2.98518  4.58477  3.61503
          0.01475  4.61986  5.34221  0.61958  0.77255  0.46239
0.99365
     65   3.48778  6.08990  2.23013  0.56154  5.40151  3.63044
4.25810  4.97407  3.35272  4.44776  5.34348  3.10414  4.29711
3.44131  3.98321  3.34124  3.79763  4.52320  6.54903  5.02884    124
E - - -
          2.68618  4.42225  2.77519  2.73123  3.46354  2.40513
3.72494  3.29354  2.67741  2.69355  4.24690  2.90347  2.73739
3.18146  2.89801  2.37887  2.77519  2.98518  4.58477  3.61503
          0.01445  4.64004  5.36239  0.61958  0.77255  0.48576
0.95510
     66   2.35381  4.73691  3.42352  3.24615  4.86680  0.75331
4.52106  4.32152  3.46835  3.95476  4.77132  3.55063  4.12317
3.14168  3.83184  2.52977  3.20272  3.74736  6.15748  4.90115    125
g - - -
          2.68618  4.42225  2.77519  2.73123  3.46354  2.40513
3.72494  3.29354  2.67741  2.69355  4.24690  2.90347  2.73739
3.18146  2.89801  2.37887  2.77519  2.98518  4.58477  3.61503
          0.01445  4.64004  5.36239  0.61958  0.77255  0.48576
0.95510
     67   3.37470  5.52649  3.75863  3.11002  5.06958  3.97404
3.93786  4.40945  0.71826  3.81540  4.67326  3.51163  4.35650
3.06604  2.18795  2.83932  3.54974  4.06314  5.82851  4.66545    126
k - - -
          2.68618  4.42225  2.77519  2.73123  3.46354  2.40513
3.72494  3.29354  2.67741  2.69355  4.24690  2.90347  2.73739
3.18146  2.89801  2.37887  2.77519  2.98518  4.58477  3.61503
          0.01445  4.64004  5.36239  0.61958  0.77255  0.48576
0.95510
     68   2.13445  0.56399  4.84262  4.66353  4.72407  3.43789
5.23011  3.77522  4.48411  3.76273  4.73905  4.23467  4.26004
4.70901  4.54330  2.94949  3.24897  3.34843  6.19415  5.07214    127
C - - -
          2.68618  4.42225  2.77519  2.73123  3.46354  2.40513
3.72494  3.29354  2.67741  2.69355  4.24690  2.90347  2.73739
3.18146  2.89801  2.37887  2.77519  2.98518  4.58477  3.61503
          0.01445  4.64004  5.36239  0.61958  0.77255  0.48576
0.95510
     69   2.67206  4.57929  3.82887  3.49568  4.43734  0.81493
4.51908  3.83649  3.49343  3.53909  2.93572  3.69429  4.12474
3.81222  3.80013  2.37195  3.12372  3.40481  5.82685  4.59351    128
g - - -
          2.68605  4.42234  2.77517  2.73121  3.46363  2.40513
3.72504  3.29363  2.67695  2.69364  4.24620  2.90356  2.73749
3.18156  2.89810  2.37888  2.77529  2.98528  4.58486  3.61512
```

```
          0.08354  2.58416  5.36239  1.27692  0.32697  0.48576
0.95510
     70   3.90954  5.97321  3.14415  0.29962  5.46672  3.99553
4.73833  5.11437  3.75074  4.61808  5.63198  3.70485  4.65402
3.98375  4.19498  3.84682  4.24203  4.73822  6.46355  5.29710    134
E - - -
          2.68618  4.42225  2.77519  2.73123  3.46354  2.40513
3.72494  3.29354  2.67741  2.69355  4.24690  2.90347  2.73739
3.18146  2.89801  2.37887  2.77519  2.98518  4.58477  3.61503
          0.01445  4.64004  5.36239  0.61958  0.77255  0.48576
0.95510
     71   2.17532  4.74377  3.51158  3.35497  4.90859  0.69405
4.59741  4.33024  3.08249  3.99549  4.82798  3.62640  4.16096
3.84546  3.84072  2.91368  3.24579  3.76164  6.19319  4.96439    135
g - - -
          2.68618  4.42225  2.77519  2.73123  3.46354  2.40513
3.72494  3.29354  2.67741  2.69355  4.24690  2.90347  2.73739
3.18146  2.89801  2.37887  2.77519  2.98518  4.58477  3.61503
          0.01445  4.64004  5.36239  0.61958  0.77255  0.48576
0.95510
     72   3.10348  5.24920  3.45289  2.97966  4.87526  3.73844
4.00156  4.24857  0.77693  3.75560  4.59809  3.40688  4.22647
3.14443  2.68636  2.69605  3.00058  3.85688  5.84767  4.61472    136
k - - -
          2.68618  4.42225  2.77519  2.73123  3.46354  2.40513
3.72494  3.29354  2.67741  2.69355  4.24690  2.90347  2.73739
3.18146  2.89801  2.37887  2.77519  2.98518  4.58477  3.61503
          0.01477  4.61826  5.34061  0.61958  0.77255  0.48576
0.95510
     73   3.72701  0.18888  5.28578  5.22343  5.15685  4.13654
5.72757  4.52227  5.10042  4.35137  5.51147  5.05549  4.88254
5.41203  5.07121  3.99314  4.25296  4.19063  6.31774  5.42009    137
C - - -
          2.68618  4.42225  2.77519  2.73123  3.46354  2.40513
3.72494  3.29354  2.67741  2.69355  4.24690  2.90347  2.73739
3.18146  2.89801  2.37887  2.77519  2.98518  4.58477  3.61503
          0.01477  4.61826  5.34061  0.61958  0.77255  0.48576
0.95510
     74   2.10351  4.70149  3.44368  3.19435  4.66988  0.87293
3.49493  4.13147  3.34961  3.77033  4.59298  3.51252  4.09155
3.64767  3.72256  2.57463  3.14812  3.61890  5.98269  4.71400    138
g - - -
          2.68618  4.42225  2.77519  2.73123  3.46354  2.40513
3.72494  3.29354  2.67741  2.69355  4.24690  2.90347  2.73739
3.18146  2.89801  2.37887  2.77519  2.98518  4.58477  3.61503
          0.01477  4.61826  5.34061  0.61958  0.77255  0.48576
0.95510
     75   1.71289  5.03477  3.02646  2.31457  4.45666  1.96930
3.56692  3.89699  2.70737  3.47146  4.27067  3.12358  3.99767
3.03733  3.17973  2.21193  3.06776  3.51214  5.68147  4.32579    139
a - - -
          2.68618  4.42225  2.77519  2.73123  3.46354  2.40513
3.72494  3.29354  2.67741  2.69355  4.24690  2.90347  2.73739
3.18146  2.89801  2.37887  2.77519  2.98518  4.58477  3.61503
```

```
          0.07292  4.61826  2.80583  0.61958  0.77255  0.48576
0.95510
     76   2.34259  5.17607  2.45522  2.26519  4.50056  3.10874
3.68431  3.97266  2.23780  3.47292  3.33633  2.46084  3.88617
2.78846  2.92056  2.22630  2.82068  3.55588  5.62189  4.22267    140
s - - -
          2.68618  4.42225  2.77519  2.73123  3.46354  2.40513
3.72494  3.29354  2.67741  2.69355  4.24690  2.90347  2.73739
3.18146  2.89801  2.37887  2.77519  2.98518  4.58477  3.61503
          0.06507  4.53313  2.95163  0.61958  0.77255  0.58755
0.81122
     77   2.63504  5.18734  3.15430  2.46654  4.52820  3.59710
3.71961  3.95541  1.27294  3.45826  3.83837  3.08701  3.98090
2.71857  2.46916  2.61736  3.05775  3.57930  5.59508  4.26872    141
k - - -
          2.68618  4.42225  2.77519  2.73123  3.46354  2.40513
3.72494  3.29354  2.67741  2.69355  4.24690  2.90347  2.73739
3.18146  2.89801  2.37887  2.77519  2.98518  4.58477  3.61503
          0.01140  4.47947        *  0.61958  0.77255  0.00000
*
//
```

```
HMMER3/f [3.3.2 | Nov 2020]
NAME  CxxxxC
LENG  63
ALPH  amino
RF    no
MM    no
CONS  yes
CS    no
MAP   yes
DATE  Fri Jun 16 17:44:00 2023
NSEQ  41
EFFN  1.303772
CKSUM 3237953633
STATS LOCAL MSV       -9.0431  0.71890
STATS LOCAL VITERBI   -9.4798  0.71890
STATS LOCAL FORWARD   -4.2647  0.71890
HMM          A        C        D        E        F        G        H
I        K        L        M        N        P        Q        R
S        T        V        W        Y
            m->m     m->i     m->d     i->m     i->i     d->m     d-
>d
  COMPO   2.49304  3.00630  2.85614  2.73092  3.89519  2.12588
3.90188  3.35404  2.36105  2.88986  4.07203  3.16611  3.69950
2.94405  2.83056  2.69631  2.88568  2.90050  5.40993  3.80849
          2.68618  4.42225  2.77519  2.73123  3.46354  2.40513
3.72494  3.29354  2.67741  2.69355  4.24690  2.90347  2.73739
3.18146  2.89801  2.37887  2.77519  2.98518  4.58477  3.61503
          0.01950  4.34304  5.06539  0.61958  0.77255  0.00000
*
      1   2.72719  4.93205  2.79986  2.29003  4.52163  1.53663
3.91957  3.97109  2.79123  3.54868  4.36101  2.82269  3.92619
3.08431  3.26267  1.99381  2.91907  3.54469  5.75401  4.39568      1
g - - -
          2.68618  4.42225  2.77519  2.73123  3.46354  2.40513
3.72494  3.29354  2.67741  2.69355  4.24690  2.90347  2.73739
3.18146  2.89801  2.37887  2.77519  2.98518  4.58477  3.61503
          0.01950  4.34304  5.06539  0.61958  0.77255  0.48576
0.95510
      2   3.35622  5.05844  4.12323  4.09300  5.17664  0.27053
5.15697  4.91769  4.33623  4.50026  5.49794  4.29007  4.46094
4.62394  4.49544  3.54533  3.87254  4.35447  6.12093  5.30320      2
G - - -
          2.68618  4.42225  2.77519  2.73123  3.46354  2.40513
3.72494  3.29354  2.67741  2.69355  4.24690  2.90347  2.73739
3.18146  2.89801  2.37887  2.77519  2.98518  4.58477  3.61503
          0.01950  4.34304  5.06539  0.61958  0.77255  0.48576
0.95510
      3   2.86193  5.26576  3.31239  2.82738  4.67774  3.70823
3.53070  4.10063  0.89757  3.57613  4.44145  3.26763  4.13965
2.95048  2.42050  3.10898  3.32170  3.76105  5.63396  4.39793      3
k - - -
          2.68618  4.42225  2.77519  2.73123  3.46354  2.40513
3.72494  3.29354  2.67741  2.69355  4.24690  2.90347  2.73739
3.18146  2.89801  2.37887  2.77519  2.98518  4.58477  3.61503
```

```
          0.01950  4.34304  5.06539  0.61958  0.77255  0.48576
0.95510
      4   3.24744  0.34056  4.82837  4.69212  4.61500  3.74725
5.24095  3.87147  4.53691  3.74811  4.89238  4.56363  4.48535
4.86314  4.56776  3.51648  3.75421  3.57504  5.89397  4.89140      4
C - - -
          2.68618  4.42225  2.77519  2.73123  3.46354  2.40513
3.72494  3.29354  2.67741  2.69355  4.24690  2.90347  2.73739
3.18146  2.89801  2.37887  2.77519  2.98518  4.58477  3.61503
          0.01950  4.34304  5.06539  0.61958  0.77255  0.48576
0.95510
      5   0.50500  4.50288  4.01767  3.86247  4.59783  3.30970
4.81674  3.80115  3.89336  3.65783  4.65549  3.86845  4.09515
4.19228  4.09055  2.85876  3.17008  3.36690  5.97224  4.83843      5
A - - -
          2.68618  4.42225  2.77519  2.73123  3.46354  2.40513
3.72494  3.29354  2.67741  2.69355  4.24690  2.90347  2.73739
3.18146  2.89801  2.37887  2.77519  2.98518  4.58477  3.61503
          0.01950  4.34304  5.06539  0.61958  0.77255  0.48576
0.95510
      6   1.77734  4.38446  3.67374  3.26513  4.24922  3.23267
4.28734  3.14042  3.24269  3.31314  4.18231  3.48155  3.91957
3.55555  3.57566  1.10256  2.90490  3.13222  5.63403  4.40039      6
s - - -
          2.68618  4.42225  2.77519  2.73123  3.46354  2.40513
3.72494  3.29354  2.67741  2.69355  4.24690  2.90347  2.73739
3.18146  2.89801  2.37887  2.77519  2.98518  4.58477  3.61503
          0.01950  4.34304  5.06539  0.61958  0.77255  0.48576
0.95510
      7   3.35622  5.05844  4.12323  4.09300  5.17664  0.27053
5.15697  4.91769  4.33623  4.50026  5.49794  4.29007  4.46094
4.62394  4.49544  3.54533  3.87254  4.35447  6.12093  5.30320      7
G - - -
          2.68618  4.42225  2.77519  2.73123  3.46354  2.40513
3.72494  3.29354  2.67741  2.69355  4.24690  2.90347  2.73739
3.18146  2.89801  2.37887  2.77519  2.98518  4.58477  3.61503
          0.01950  4.34304  5.06539  0.61958  0.77255  0.48576
0.95510
      8   3.48549  5.40647  3.68944  3.25316  4.90578  3.89071
4.12519  4.37424  0.53539  3.84657  4.80652  3.64650  4.37718
3.30688  2.65070  3.51095  3.71446  4.06438  5.78082  4.67739      8
K - - -
          2.68618  4.42225  2.77519  2.73123  3.46354  2.40513
3.72494  3.29354  2.67741  2.69355  4.24690  2.90347  2.73739
3.18146  2.89801  2.37887  2.77519  2.98518  4.58477  3.61503
          0.01950  4.34304  5.06539  0.61958  0.77255  0.48576
0.95510
      9   3.24744  0.34056  4.82837  4.69212  4.61500  3.74725
5.24095  3.87147  4.53691  3.74811  4.89238  4.56363  4.48535
4.86314  4.56776  3.51648  3.75421  3.57504  5.89397  4.89140      9
C - - -
          2.68618  4.42225  2.77519  2.73123  3.46354  2.40513
3.72494  3.29354  2.67741  2.69355  4.24690  2.90347  2.73739
3.18146  2.89801  2.37887  2.77519  2.98518  4.58477  3.61503
```

```
          0.01950  4.34304  5.06539  0.61958  0.77255  0.48576
0.95510
     10   3.35622  5.05844  4.12323  4.09300  5.17664  0.27053
5.15697  4.91769  4.33623  4.50026  5.49794  4.29007  4.46094
4.62394  4.49544  3.54533  3.87254  4.35447  6.12093  5.30320      10
G - - -
          2.68618  4.42225  2.77519  2.73123  3.46354  2.40513
3.72494  3.29354  2.67741  2.69355  4.24690  2.90347  2.73739
3.18146  2.89801  2.37887  2.77519  2.98518  4.58477  3.61503
          0.01950  4.34304  5.06539  0.61958  0.77255  0.48576
0.95510
     11   2.48397  4.44945  3.43280  3.13585  4.42920  2.88869
4.27690  3.84676  3.21642  3.51618  4.34440  3.03043  3.90774
3.52388  3.56521  2.30416  0.99827  3.35250  5.75980  4.52041      11
t - - -
          2.68618  4.42225  2.77519  2.73123  3.46354  2.40513
3.72494  3.29354  2.67741  2.69355  4.24690  2.90347  2.73739
3.18146  2.89801  2.37887  2.77519  2.98518  4.58477  3.61503
          0.01950  4.34304  5.06539  0.61958  0.77255  0.48576
0.95510
     12   2.74715  4.80305  3.12173  1.46818  4.02596  3.56164
3.83150  3.17444  2.64437  2.56101  3.88517  3.14759  3.98267
3.00991  3.07145  2.82754  2.83137  2.55231  5.39021  4.07982      12
e - - -
          2.68618  4.42225  2.77519  2.73123  3.46354  2.40513
3.72494  3.29354  2.67741  2.69355  4.24690  2.90347  2.73739
3.18146  2.89801  2.37887  2.77519  2.98518  4.58477  3.61503
          0.01950  4.34304  5.06539  0.61958  0.77255  0.48576
0.95510
     13   3.48549  5.40647  3.68944  3.25316  4.90578  3.89071
4.12519  4.37424  0.53539  3.84657  4.80652  3.64650  4.37718
3.30688  2.65070  3.51095  3.71446  4.06438  5.78082  4.67739      13
K - - -
          2.68618  4.42225  2.77519  2.73123  3.46354  2.40513
3.72494  3.29354  2.67741  2.69355  4.24690  2.90347  2.73739
3.18146  2.89801  2.37887  2.77519  2.98518  4.58477  3.61503
          0.01950  4.34304  5.06539  0.61958  0.77255  0.48576
0.95510
     14   2.56120  4.42658  3.48043  2.90767  3.40125  3.62175
3.85678  1.94229  2.53423  2.62876  3.21596  3.32767  4.00048
2.99210  2.39688  2.86515  2.89434  2.63729  4.99941  3.76029      14
i - - -
          2.68618  4.42225  2.77519  2.73123  3.46354  2.40513
3.72494  3.29354  2.67741  2.69355  4.24690  2.90347  2.73739
3.18146  2.89801  2.37887  2.77519  2.98518  4.58477  3.61503
          0.01950  4.34304  5.06539  0.61958  0.77255  0.48576
0.95510
     15   4.18963  5.26606  5.11024  4.92502  1.15712  4.80429
3.63222  3.80574  4.73656  3.04072  4.36611  4.41321  5.08471
4.53749  4.64518  4.19975  4.39932  3.75018  3.72801  0.88582      15
y - - -
          2.68618  4.42225  2.77519  2.73123  3.46354  2.40513
3.72494  3.29354  2.67741  2.69355  4.24690  2.90347  2.73739
3.18146  2.89801  2.37887  2.77519  2.98518  4.58477  3.61503
```

```
          0.01950  4.34304  5.06539  0.61958  0.77255  0.48576
0.95510
     16   2.17473  4.95019  2.83329  1.85450  4.48379  1.97249
3.85034  3.93315  2.69337  3.50006  4.29844  3.01691  3.90154
2.99967  3.17335  2.25626  3.00898  3.51258  5.69774  4.33144      16
e - - -
          2.68618  4.42225  2.77519  2.73123  3.46354  2.40513
3.72494  3.29354  2.67741  2.69355  4.24690  2.90347  2.73739
3.18146  2.89801  2.37887  2.77519  2.98518  4.58477  3.61503
          0.01950  4.34304  5.06539  0.61958  0.77255  0.48576
0.95510
     17   2.58088  5.11117  3.05573  2.43998  4.43965  3.53232
3.68417  3.86117  1.58697  3.38461  4.17991  3.02993  3.92862
2.04036  2.68907  2.77499  3.00684  3.06643  5.54029  4.20949      17
k - - -
          2.68618  4.42225  2.77519  2.73123  3.46354  2.40513
3.72494  3.29354  2.67741  2.69355  4.24690  2.90347  2.73739
3.18146  2.89801  2.37887  2.77519  2.98518  4.58477  3.61503
          0.01950  4.34304  5.06539  0.61958  0.77255  0.48576
0.95510
     18   1.78871  4.69938  3.17428  2.61677  3.91578  3.51858
3.74372  3.21784  2.43825  2.95363  3.79723  3.10822  3.91298
2.62952  2.84205  2.63451  2.67738  2.61068  5.24187  3.94564      18
a - - -
          2.68618  4.42225  2.77519  2.73123  3.46354  2.40513
3.72494  3.29354  2.67741  2.69355  4.24690  2.90347  2.73739
3.18146  2.89801  2.37887  2.77519  2.98518  4.58477  3.61503
          0.01950  4.34304  5.06539  0.61958  0.77255  0.48576
0.95510
     19   2.65495  5.23294  2.37317  1.91708  4.56545  3.42631
3.53867  4.05079  1.89922  3.53153  4.28252  2.52865  3.85608
2.69577  2.92779  2.62214  2.76665  3.61791  5.66697  4.24934      19
k - - -
          2.68618  4.42225  2.77519  2.73123  3.46354  2.40513
3.72494  3.29354  2.67741  2.69355  4.24690  2.90347  2.73739
3.18146  2.89801  2.37887  2.77519  2.98518  4.58477  3.61503
          0.01950  4.34304  5.06539  0.61958  0.77255  0.48576
0.95510
     20   3.13565  4.49142  4.80567  4.24002  3.35254  4.37631
4.74736  1.78217  4.10326  1.08818  3.19703  4.42861  4.01122
4.27114  4.23320  3.70843  3.37174  1.80545  5.21662  4.08616      20
l - - -
          2.68618  4.42225  2.77519  2.73123  3.46354  2.40513
3.72494  3.29354  2.67741  2.69355  4.24690  2.90347  2.73739
3.18146  2.89801  2.37887  2.77519  2.98518  4.58477  3.61503
          0.01950  4.34304  5.06539  0.61958  0.77255  0.48576
0.95510
     21   2.59908  5.11681  2.41519  2.23731  4.44064  3.43455
3.62698  3.75936  2.10318  3.22812  4.16367  2.59524  3.82991
2.50457  2.85981  2.48996  2.55429  3.29640  5.56206  4.16395      21
k - - -
          2.68618  4.42225  2.77519  2.73123  3.46354  2.40513
3.72494  3.29354  2.67741  2.69355  4.24690  2.90347  2.73739
3.18146  2.89801  2.37887  2.77519  2.98518  4.58477  3.61503
```

```
          0.01950  4.34304  5.06539  0.61958  0.77255  0.48576
0.95510
     22   2.48940  5.01621  2.93120  2.42650  4.32625  3.09265
2.26818  3.77146  2.44630  3.32129  4.10493  2.65234  3.86241
2.50313  2.91894  2.35163  2.68149  3.39631  5.51232  4.14325      22
h - - -
          2.68618  4.42225  2.77519  2.73123  3.46354  2.40513
3.72494  3.29354  2.67741  2.69355  4.24690  2.90347  2.73739
3.18146  2.89801  2.37887  2.77519  2.98518  4.58477  3.61503
          0.01950  4.34304  5.06539  0.61958  0.77255  0.48576
0.95510
     23   3.13227  5.67045  0.89559  2.28750  5.03845  3.37993
3.99402  4.58323  3.05946  4.08961  4.95450  2.28080  4.03358
3.17035  3.67626  3.02179  3.21432  4.13010  6.23895  4.70973      23
d - - -
          2.68618  4.42225  2.77519  2.73123  3.46354  2.40513
3.72494  3.29354  2.67741  2.69355  4.24690  2.90347  2.73739
3.18146  2.89801  2.37887  2.77519  2.98518  4.58477  3.61503
          0.01950  4.34304  5.06539  0.61958  0.77255  0.48576
0.95510
     24   2.86476  5.03642  2.78462  2.29333  4.40635  3.45471
3.56194  3.85305  2.69361  3.44602  4.29774  3.06152  1.29153
3.07276  3.10182  2.88547  2.96625  3.50248  5.65352  4.30940      24
p - - -
          2.68618  4.42225  2.77519  2.73123  3.46354  2.40513
3.72494  3.29354  2.67741  2.69355  4.24690  2.90347  2.73739
3.18146  2.89801  2.37887  2.77519  2.98518  4.58477  3.61503
          0.01950  4.34304  5.06539  0.61958  0.77255  0.48576
0.95510
     25   2.95061  5.21841  2.77984  2.54340  4.60227  3.48306
3.87230  4.13116  2.56230  3.65216  4.49397  2.56561  4.02263
1.13290  2.92824  2.80641  3.23036  3.73818  5.75246  4.38429      25
q - - -
          2.68618  4.42225  2.77519  2.73123  3.46354  2.40513
3.72494  3.29354  2.67741  2.69355  4.24690  2.90347  2.73739
3.18146  2.89801  2.37887  2.77519  2.98518  4.58477  3.61503
          0.01950  4.34304  5.06539  0.61958  0.77255  0.48576
0.95510
     26   3.00041  5.53644  1.25942  2.12239  4.82973  2.18902
3.86647  4.33685  2.81518  3.83595  4.65240  2.74748  3.97514
3.01679  3.38378  2.90970  3.27245  3.91391  5.99953  4.06856      26
d - - -
          2.68619  4.42226  2.77520  2.73124  3.46355  2.40502
3.72495  3.29355  2.67742  2.69356  4.24691  2.90347  2.73740
3.18147  2.89802  2.37888  2.77520  2.98519  4.58478  3.61504
          0.06834  2.81759  5.06539  0.41986  1.07043  0.48576
0.95510
     27   2.80621  5.10010  3.16726  2.56708  4.42477  3.57155
3.38771  3.84116  1.82904  2.72855  4.16013  3.06932  3.94836
2.07829  2.13973  2.80168  3.01824  3.20579  5.50799  4.19874      28
k - - -
          2.68618  4.42225  2.77519  2.73123  3.46354  2.40513
3.72494  3.29354  2.67741  2.69355  4.24690  2.90347  2.73739
3.18146  2.89801  2.37887  2.77519  2.98518  4.58477  3.61503
```

```
          0.01950  4.34304  5.06539  0.61958  0.77255  0.48576
0.95510
     28   3.66836  4.97658  5.19522  4.67876  3.19387  4.87304
5.17123  2.38359  4.42889  0.56549  2.76946  4.93627  5.03185
4.56569  4.51506  4.28697  3.89790  2.58023  5.36227  4.27179     29
L - - -
          2.68618  4.42225  2.77519  2.73123  3.46354  2.40513
3.72494  3.29354  2.67741  2.69355  4.24690  2.90347  2.73739
3.18146  2.89801  2.37887  2.77519  2.98518  4.58477  3.61503
          0.01950  4.34304  5.06539  0.61958  0.77255  0.48576
0.95510
     29   2.83677  4.43773  4.35190  4.02646  3.90302  3.76904
4.79533  2.30762  3.91400  2.70906  3.85894  4.11335  4.39226
4.24185  4.11034  2.91822  3.23044  0.75415  5.60735  4.34189     30
v - - -
          2.68618  4.42225  2.77519  2.73123  3.46354  2.40513
3.72494  3.29354  2.67741  2.69355  4.24690  2.90347  2.73739
3.18146  2.89801  2.37887  2.77519  2.98518  4.58477  3.61503
          0.01950  4.34304  5.06539  0.61958  0.77255  0.48576
0.95510
     30   3.16428  5.09380  3.70663  3.08894  3.74319  3.84821
3.82647  3.77326  2.32898  3.27888  4.23195  3.46670  4.23893
3.09916  0.99511  3.22219  3.36697  3.51954  5.10207  2.63539     31
r - - -
          2.68618  4.42225  2.77519  2.73123  3.46354  2.40513
3.72494  3.29354  2.67741  2.69355  4.24690  2.90347  2.73739
3.18146  2.89801  2.37887  2.77519  2.98518  4.58477  3.61503
          0.01950  4.34304  5.06539  0.61958  0.77255  0.48576
0.95510
     31   0.80970  4.29406  3.96252  3.61091  4.34044  3.17675
4.52266  3.50800  3.56241  3.38576  4.27242  3.64493  3.92751
3.84579  3.82284  2.50266  2.56529  2.77591  5.75880  4.57440     32
a - - -
          2.68618  4.42225  2.77519  2.73123  3.46354  2.40513
3.72494  3.29354  2.67741  2.69355  4.24690  2.90347  2.73739
3.18146  2.89801  2.37887  2.77519  2.98518  4.58477  3.61503
          0.01950  4.34304  5.06539  0.61958  0.77255  0.48576
0.95510
     32   3.62737  5.38013  4.17354  3.61542  4.86109  3.98875
4.28659  4.46062  2.58540  3.89009  4.89883  3.92761  4.48762
3.50219  0.45014  3.70264  3.87328  4.16871  5.76267  4.71093     33
R - - -
          2.68618  4.42225  2.77519  2.73123  3.46354  2.40513
3.72494  3.29354  2.67741  2.69355  4.24690  2.90347  2.73739
3.18146  2.89801  2.37887  2.77519  2.98518  4.58477  3.61503
          0.01950  4.34304  5.06539  0.61958  0.77255  0.48576
0.95510
     33   3.57839  5.69886  0.42649  2.76099  5.15862  3.64405
4.44917  4.87224  3.63624  4.40164  5.41486  3.33250  4.33018
3.70565  4.19804  3.50372  3.93459  4.46785  6.22315  4.98548     34
D - - -
          2.68618  4.42225  2.77519  2.73123  3.46354  2.40513
3.72494  3.29354  2.67741  2.69355  4.24690  2.90347  2.73739
3.18146  2.89801  2.37887  2.77519  2.98518  4.58477  3.61503
```

```
         0.01950  4.34304  5.06539  0.61958  0.77255  0.48576
0.95510
     34  3.35622  5.05844  4.12323  4.09300  5.17664  0.27053
5.15697  4.91769  4.33623  4.50026  5.49794  4.29007  4.46094
4.62394  4.49544  3.54533  3.87254  4.35447  6.12093  5.30320      35
G - - -
         2.68618  4.42225  2.77519  2.73123  3.46354  2.40513
3.72494  3.29354  2.67741  2.69355  4.24690  2.90347  2.73739
3.18146  2.89801  2.37887  2.77519  2.98518  4.58477  3.61503
         0.01950  4.34304  5.06539  0.61958  0.77255  0.48576
0.95510
     35  3.02239  5.29522  3.21496  2.68885  4.68798  3.66985
3.73409  4.09060  1.09466  3.33498  4.37740  2.76198  4.06748
2.49348  2.44142  2.99772  3.21922  3.73290  5.62747  4.36654      36
k - - -
         2.68618  4.42225  2.77519  2.73123  3.46354  2.40513
3.72494  3.29354  2.67741  2.69355  4.24690  2.90347  2.73739
3.18146  2.89801  2.37887  2.77519  2.98518  4.58477  3.61503
         0.01950  4.34304  5.06539  0.61958  0.77255  0.48576
0.95510
     36  3.24744  0.34056  4.82837  4.69212  4.61500  3.74725
5.24095  3.87147  4.53691  3.74811  4.89238  4.56363  4.48535
4.86314  4.56776  3.51648  3.75421  3.57504  5.89397  4.89140      37
C - - -
         2.68618  4.42225  2.77519  2.73123  3.46354  2.40513
3.72494  3.29354  2.67741  2.69355  4.24690  2.90347  2.73739
3.18146  2.89801  2.37887  2.77519  2.98518  4.58477  3.61503
         0.01950  4.34304  5.06539  0.61958  0.77255  0.48576
0.95510
     37  3.35622  5.05844  4.12323  4.09300  5.17664  0.27053
5.15697  4.91769  4.33623  4.50026  5.49794  4.29007  4.46094
4.62394  4.49544  3.54533  3.87254  4.35447  6.12093  5.30320      38
G - - -
         2.68618  4.42225  2.77519  2.73123  3.46354  2.40513
3.72494  3.29354  2.67741  2.69355  4.24690  2.90347  2.73739
3.18146  2.89801  2.37887  2.77519  2.98518  4.58477  3.61503
         0.01950  4.34304  5.06539  0.61958  0.77255  0.48576
0.95510
     38  2.64448  4.28416  3.66836  3.09688  3.40992  3.65915
3.92740  2.54640  2.82987  1.83714  3.24647  3.45933  3.31075
3.22452  3.34238  2.78211  2.34540  2.34050  4.87900  3.65979      39
l - - -
         2.68618  4.42225  2.77519  2.73123  3.46354  2.40513
3.72494  3.29354  2.67741  2.69355  4.24690  2.90347  2.73739
3.18146  2.89801  2.37887  2.77519  2.98518  4.58477  3.61503
         0.01950  4.34304  5.06539  0.61958  0.77255  0.48576
0.95510
     39  2.57396  5.01011  2.76041  2.34098  4.30310  3.23190
3.44709  3.74478  2.39487  3.30405  4.09356  2.95831  3.86698
2.81799  2.95233  1.89257  2.16356  3.37831  5.50546  4.13701      40
s - - -
         2.68624  4.42231  2.77525  2.73129  3.46359  2.40507
3.72500  3.29360  2.67746  2.69360  4.24695  2.90352  2.73721
3.18152  2.89806  2.37892  2.77492  2.98503  4.58482  3.61509
```

0.04466 3.28691 5.06539 1.42501 0.27510 0.48576
0.95510
40 2.26617 4.48077 3.37430 2.86567 4.23027 1.22882
4.15015 3.58674 3.08494 3.29434 4.14365 3.34726 3.92185
3.39366 3.45771 2.50290 2.93158 2.62621 5.58531 4.33391 52
g - - -
2.68618 4.42225 2.77519 2.73123 3.46354 2.40513
3.72494 3.29354 2.67741 2.69355 4.24690 2.90347 2.73739
3.18146 2.89801 2.37887 2.77519 2.98518 4.58477 3.61503
0.01950 4.34304 5.06539 0.61958 0.77255 0.48576
0.95510
41 2.57568 4.41320 2.87737 2.11009 4.42641 3.33895
3.62774 3.89383 2.11582 3.39886 4.15283 2.74185 3.83290
2.20065 2.52419 2.63303 2.77573 3.48308 5.55104 3.70833 53
e - - -
2.68618 4.42225 2.77519 2.73123 3.46354 2.40513
3.72494 3.29354 2.67741 2.69355 4.24690 2.90347 2.73739
3.18146 2.89801 2.37887 2.77519 2.98518 4.58477 3.61503
0.01950 4.34304 5.06539 0.61958 0.77255 0.48576
0.95510
42 2.37286 4.64132 3.17078 2.88813 4.47618 1.13073
4.11173 3.86955 2.53182 3.51735 4.35266 3.26896 3.94208
3.31731 3.30074 2.72877 2.76644 3.42028 5.74834 4.47674 54
g - - -
2.68618 4.42225 2.77519 2.73123 3.46354 2.40513
3.72494 3.29354 2.67741 2.69355 4.24690 2.90347 2.73739
3.18146 2.89801 2.37887 2.77519 2.98518 4.58477 3.61503
0.01950 4.34304 5.06539 0.61958 0.77255 0.48576
0.95510
43 2.54990 4.08256 4.28870 3.69412 2.84797 3.78225
3.60988 1.85712 3.54453 1.96545 3.18937 3.82962 4.14771
3.73501 3.66666 3.07866 2.68370 2.04836 4.69557 2.93735 55
i - - -
2.68618 4.42225 2.77519 2.73123 3.46354 2.40513
3.72494 3.29354 2.67741 2.69355 4.24690 2.90347 2.73739
3.18146 2.89801 2.37887 2.77519 2.98518 4.58477 3.61503
0.01950 4.34304 5.06539 0.61958 0.77255 0.48576
0.95510
44 2.44576 5.13197 2.73033 2.23300 4.46207 3.20108
3.41768 3.93751 2.15130 3.43046 4.17699 2.41534 3.82763
2.58080 2.61392 2.45710 2.83137 3.51462 5.57310 3.56812 56
k - - -
2.68618 4.42225 2.77519 2.73123 3.46354 2.40513
3.72494 3.29354 2.67741 2.69355 4.24690 2.90347 2.73739
3.18146 2.89801 2.37887 2.77519 2.98518 4.58477 3.61503
0.01950 4.34304 5.06539 0.61958 0.77255 0.48576
0.95510
45 2.31919 4.84817 3.04314 2.45380 4.10013 2.87062
3.69676 3.51687 2.45869 3.12096 3.93435 2.76007 3.00478
2.62848 2.96121 2.59797 2.72151 2.47943 5.36533 4.03113 57
a - - -
2.68618 4.42225 2.77519 2.73123 3.46354 2.40513
3.72494 3.29354 2.67741 2.69355 4.24690 2.90347 2.73739
3.18146 2.89801 2.37887 2.77519 2.98518 4.58477 3.61503

0.01950  4.34304  5.06539  0.61958  0.77255  0.48576
0.95510
46  2.48036  5.08289  2.56468  2.27407  4.39528  3.43900
3.53136  3.85882  2.15522  3.37409  3.66428  2.70557  3.29852
2.63197  2.86652  2.54970  2.43776  3.17342  5.53560  4.14520  58
k - - -
2.68618  4.42225  2.77519  2.73123  3.46354  2.40513
3.72494  3.29354  2.67741  2.69355  4.24690  2.90347  2.73739
3.18146  2.89801  2.37887  2.77519  2.98518  4.58477  3.61503
0.01950  4.34304  5.06539  0.61958  0.77255  0.48576
0.95510
47  2.33970  5.10874  2.77385  2.07275  4.42930  3.26410
3.62944  3.89829  2.12405  3.21743  4.15685  2.85557  3.19816
2.48601  2.86320  2.63195  2.61773  3.37970  5.55629  4.16039  59
e - - -
2.68618  4.42225  2.77519  2.73123  3.46354  2.40513
3.72494  3.29354  2.67741  2.69355  4.24690  2.90347  2.73739
3.18146  2.89801  2.37887  2.77519  2.98518  4.58477  3.61503
0.04754  4.34304  3.39836  0.61958  0.77255  0.48576
0.95510
48  2.37554  5.06600  2.64895  2.16522  4.37295  3.43247
3.63551  3.83239  2.35327  3.35654  4.12142  2.74305  3.43587
2.68426  2.88008  2.25807  2.42494  3.16384  5.52610  4.13889  60
e - - -
2.68597  4.42237  2.77523  2.73135  3.46366  2.40524
3.72506  3.29366  2.67716  2.69363  4.24701  2.90358  2.73724
3.18145  2.89803  2.37880  2.77503  2.98530  4.58489  3.61515
0.69997  1.28202  1.48754  0.73706  0.65108  0.52464
0.89592
49  2.58970  4.73813  3.02240  2.19743  3.96474  3.42892
3.62939  3.18623  2.30534  2.84989  3.65707  2.97293  3.81720
2.36028  2.80868  2.63805  2.76563  2.88961  5.24590  3.48462  66
e - - -
2.68618  4.42225  2.77519  2.73123  3.46354  2.40513
3.72494  3.29354  2.67741  2.69355  4.24690  2.90347  2.73739
3.18146  2.89801  2.37887  2.77519  2.98518  4.58477  3.61503
0.02575  4.06781  4.79016  0.61958  0.77255  0.67401
0.71265
50  2.53791  4.49541  3.23006  2.61690  3.66642  3.50145
3.73014  2.64824  2.40064  2.72231  3.58928  3.13564  3.72028
2.70968  3.03850  2.56674  2.60745  2.33251  5.05095  3.62641  67
v - - -
2.68618  4.42225  2.77519  2.73123  3.46354  2.40513
3.72494  3.29354  2.67741  2.69355  4.24690  2.90347  2.73739
3.18146  2.89801  2.37887  2.77519  2.98518  4.58477  3.61503
0.02413  4.13208  4.85443  0.61958  0.77255  0.65940
0.72808
51  2.08487  4.58204  3.08081  2.50304  3.77043  3.48577
3.71092  3.01432  2.51201  2.65473  3.67280  3.08909  3.32842
2.92157  2.92175  2.65957  2.70109  2.52474  5.12792  3.84196  68
a - - -
2.68618  4.42225  2.77519  2.73123  3.46354  2.40513
3.72494  3.29354  2.67741  2.69355  4.24690  2.90347  2.73739
3.18146  2.89801  2.37887  2.77519  2.98518  4.58477  3.61503

```
          0.02322  4.16998  4.89233  0.61958  0.77255  0.41476
1.08028
     52   2.46159  4.40290  2.67444  2.29499  4.39263  3.16193
3.62393  3.85683  2.30561  3.37078  4.12846  2.63045  3.82574
2.32293  2.65077  2.45860  2.61263  3.24805  5.53111  4.14017      69
e - - -
          2.68618  4.42225  2.77519  2.73123  3.46354  2.40513
3.72494  3.29354  2.67741  2.69355  4.24690  2.90347  2.73739
3.18146  2.89801  2.37887  2.77519  2.98518  4.58477  3.61503
          0.01996  4.31979  5.04214  0.61958  0.77255  0.49238
0.94462
     53   2.55384  4.64250  2.63368  2.34728  4.34038  3.26966
3.63474  3.61205  2.31548  2.88742  4.09467  2.57515  3.36170
2.55344  2.76764  2.25007  2.88028  3.31441  5.50278  4.12162      70
s - - -
          2.68618  4.42225  2.77519  2.73123  3.46354  2.40513
3.72494  3.29354  2.67741  2.69355  4.24690  2.90347  2.73739
3.18146  2.89801  2.37887  2.77519  2.98518  4.58477  3.61503
          0.01973  4.33127  5.05361  0.61958  0.77255  0.47529
0.97207
     54   2.66580  4.67977  2.60004  2.19884  4.41612  2.42911
3.63937  3.88141  2.32714  3.39431  4.15327  2.85530  3.83783
2.67210  2.75776  2.17495  2.71192  3.47604  5.55409  4.16110      71
s - - -
          2.68618  4.42225  2.77519  2.73123  3.46354  2.40513
3.72494  3.29354  2.67741  2.69355  4.24690  2.90347  2.73739
3.18146  2.89801  2.37887  2.77519  2.98518  4.58477  3.61503
          0.01950  4.34304  5.06539  0.61958  0.77255  0.48576
0.95510
     55   3.06357  5.32339  3.40427  2.76656  4.76609  3.73037
3.71631  4.13545  1.12311  3.56662  4.39219  2.99875  4.09569
2.59669  1.99523  3.03845  2.99832  3.77822  5.62038  4.39624      72
k - - -
          2.68618  4.42225  2.77519  2.73123  3.46354  2.40513
3.72494  3.29354  2.67741  2.69355  4.24690  2.90347  2.73739
3.18146  2.89801  2.37887  2.77519  2.98518  4.58477  3.61503
          0.01950  4.34304  5.06539  0.61958  0.77255  0.48576
0.95510
     56   2.66007  2.31489  4.29655  3.70247  3.07087  3.79312
4.11105  2.26614  3.55388  1.91401  2.76535  3.84071  4.15811
3.53080  3.67773  2.75246  2.89270  2.15580  4.71445  3.53054      73
l - - -
          2.68618  4.42225  2.77519  2.73123  3.46354  2.40513
3.72494  3.29354  2.67741  2.69355  4.24690  2.90347  2.73739
3.18146  2.89801  2.37887  2.77519  2.98518  4.58477  3.61503
          0.01950  4.34304  5.06539  0.61958  0.77255  0.48576
0.95510
     57   1.95957  3.54159  3.88265  3.33854  3.54027  3.53246
4.10932  2.71510  3.26351  2.60717  3.51952  3.58966  4.03436
3.52998  3.53951  2.33669  2.22225  1.86888  5.02160  3.81569      74
v - - -
          2.68618  4.42225  2.77519  2.73123  3.46354  2.40513
3.72494  3.29354  2.67741  2.69355  4.24690  2.90347  2.73739
3.18146  2.89801  2.37887  2.77519  2.98518  4.58477  3.61503
```

0.01950 4.34304 5.06539 0.61958 0.77255 0.48576
0.95510
58 2.71763 5.19017 2.67528 1.90256 4.51404 2.49001
3.66976 3.99026 2.44103 3.48849 4.24740 2.60879 3.85742
2.69931 2.74614 2.19979 2.85897 3.57161 5.63808 4.22909 75
e - - -
2.68618 4.42225 2.77519 2.73123 3.46354 2.40513
3.72494 3.29354 2.67741 2.69355 4.24690 2.90347 2.73739
3.18146 2.89801 2.37887 2.77519 2.98518 4.58477 3.61503
0.01950 4.34304 5.06539 0.61958 0.77255 0.48576
0.95510
59 2.73834 4.71173 3.20088 3.13373 4.77790 0.69118
4.47816 4.37511 3.48434 4.00675 4.86606 2.81816 4.05264
3.75501 3.82531 2.87123 3.22501 3.78434 6.03149 4.78832 76
G - - -
2.68618 4.42225 2.77519 2.73123 3.46354 2.40513
3.72494 3.29354 2.67741 2.69355 4.24690 2.90347 2.73739
3.18146 2.89801 2.37887 2.77519 2.98518 4.58477 3.61503
0.01950 4.34304 5.06539 0.61958 0.77255 0.48576
0.95510
60 2.92429 4.37579 4.33466 3.76165 3.45822 4.07295
4.40810 2.13816 3.57049 1.94014 3.37628 4.01964 4.42048
3.88084 3.08625 3.38170 3.16440 1.15648 5.12248 3.92576 77
v - - -
2.68618 4.42225 2.77519 2.73123 3.46354 2.40513
3.72494 3.29354 2.67741 2.69355 4.24690 2.90347 2.73739
3.18146 2.89801 2.37887 2.77519 2.98518 4.58477 3.61503
0.01950 4.34304 5.06539 0.61958 0.77255 0.48576
0.95510
61 2.23174 0.75859 4.48745 4.24615 4.30663 3.27948
4.87375 3.31732 4.06275 3.31528 4.33634 3.98237 4.06765
4.33979 4.17065 2.78753 3.05137 2.96788 5.80762 4.64536 78
c - - -
2.68618 4.42225 2.77519 2.73123 3.46354 2.40513
3.72494 3.29354 2.67741 2.69355 4.24690 2.90347 2.73739
3.18146 2.89801 2.37887 2.77519 2.98518 4.58477 3.61503
0.01950 4.34304 5.06539 0.61958 0.77255 0.48576
0.95510
62 2.23632 4.47863 3.64781 3.55497 4.80805 0.65500
4.70444 4.23948 3.77210 3.96378 4.82486 3.66120 4.00008
4.02600 4.02581 2.74018 3.08887 3.61442 6.09850 4.95191 79
G - - -
2.68618 4.42225 2.77519 2.73123 3.46354 2.40513
3.72494 3.29354 2.67741 2.69355 4.24690 2.90347 2.73739
3.18146 2.89801 2.37887 2.77519 2.98518 4.58477 3.61503
0.01950 4.34304 5.06539 0.61958 0.77255 0.48576
0.95510
63 2.73250 5.22676 3.13914 2.58547 4.60631 3.59626
3.69490 4.02157 1.82098 3.49564 4.29452 2.96130 3.98714
1.56995 2.34040 2.78510 3.09901 3.64432 5.60031 4.30045 80
q - - -
2.68618 4.42225 2.77519 2.73123 3.46354 2.40513
3.72494 3.29354 2.67741 2.69355 4.24690 2.90347 2.73739
3.18146 2.89801 2.37887 2.77519 2.98518 4.58477 3.61503

```
          0.01317  4.33671        *  0.61958  0.77255  0.00000
*
//
```
